# Plural molecular and cellular mechanisms of pore domain *KCNQ2* encephalopathy

**DOI:** 10.1101/2024.01.04.574177

**Authors:** Timothy J. Abreo, Emma C. Thompson, Anuraag Madabushi, Heun Soh, Nissi Varghese, Carlos G. Vanoye, Kristen Springer, Kristen L. Park, Jim Johnson, Scotty Sims, Zhigang Ji, Ana G. Chavez, Miranda J. Jankovic, Bereket Habte, Aamir R. Zuberi, Cathleen Lutz, Zhao Wang, Vaishnav Krishnan, Lisa Dudler, Stephanie Einsele-Scholz, Jeffrey L. Noebels, Alfred L. George, Atul Maheshwari, Anastasios V. Tzingounis, Edward C. Cooper

## Abstract

*KCNQ2* variants in children with neurodevelopmental impairment are difficult to assess due to their heterogeneity and unclear pathogenic mechanisms. We describe a child with neonatal-onset epilepsy, developmental impairment of intermediate severity, and *KCNQ2* G256W heterozygosity. Analyzing prior KCNQ2 channel cryoelectron microscopy models revealed G256 as a node of an arch-shaped non-covalent bond network linking S5, the pore turret, and the ion path. Co-expression with G256W dominantly suppressed conduction by wild-type subunits in heterologous cells. Ezogabine partly reversed this suppression. G256W/+ mice have epilepsy leading to premature deaths. Hippocampal CA1 pyramidal cells from G256W/+ brain slices showed hyperexcitability. G256W/+ pyramidal cell KCNQ2 and KCNQ3 immunolabeling was significantly shifted from axon initial segments to neuronal somata. Despite normal mRNA levels, G256W/+ mouse KCNQ2 protein levels were reduced by about 50%. Our findings indicate that G256W pathogenicity results from multiplicative effects, including reductions in intrinsic conduction, subcellular targeting, and protein stability. These studies provide evidence for an unexpected and novel role for the KCNQ2 pore turret and introduce a valid animal model of *KCNQ2* encephalopathy. Our results, spanning structure to behavior, may be broadly applicable because the majority of *KCNQ2* encephalopathy patients share variants near the selectivity filter.

## Introduction

*KCNQ2* variants are among the most frequent diagnostic findings from genetic tests for epilepsy in young children (Symonds et al., 2019; Truty et al., 2019). Such test results often raise new questions about pathogenicity and developmental prognosis. This uncertainty is partly due to the great diversity of different alleles known among individuals seeking care (n = 1954 in NCBI ClinVar; Landrum et al., 2018). A second contributor is the broad phenotypic spectrum associated with *KCNQ2* variants (Weckhuysen and George, 2022). At the milder end, individuals may have seizures restricted to the first weeks or months of life and good later development—a disorder called self-limited familial neonatal epilepsy (SLFNE; Ronen et al., 1993; Scheffer et al., 2017). At the most highly impaired end, individuals have treatment-refractory, neonatal-onset seizures accompanied by lifelong profound global disability, a disorder first called “*KCNQ2* encephalopathy” (Weckhuysen et al., 2012), with electroclinical features akin to the older diagnostic group with very poor prognosis, Ohtahara syndrome (Beal et al., 2012; Olson et al., 2017). Case series reveal a middle group with de novo missense or small indel variants where seizures may remit early, and considerable childhood development of motor, receptive language, and social abilities takes place, albeit with delay. For many such affected people, spoken language remains absent, and other features including autism spectrum disorder, recurring if infrequent convulsions, and inability to perform activities of daily life independently impose significant limitations (Weckhuysen et al., 2013; Millichap et al., 2016). This has led to the concept of a *KCNQ2* developmental and epileptic encephalopathy (DEE) spectrum (Dirkx et al., 2020; Berg et al., 2021). Efforts to correlate developmental prognosis to variant functional consequences within critical structural domains have been made (Millichap and Cooper, 2012; Millichap *et al*., 2016; Goto et al., 2019; Zhang et al., 2020; Brunger et al., 2023), but prediction can likely be helped by more confluent biological evidence.

Voltage-gated K^+^ (Kv) channels contain voltage-sensing (VSD) and pore-gating (PGD) structural domains. During channel activity, conformational changes in the two domains are coupled giving rise to ion current. Here we analyze KCNQ2 G256W, a PGD variant found in an infant whose neonatal seizures remitted in early infancy and who subsequently gained neurodevelopmental milestones on an intermediate severity trajectory. G256W maps to the PGD turret near the top of the S5 helix. The G256 location is far from the primary channel components needed for ion flow (voltage sensor, pore gate, and selectivity filter), raising questions about pathogenicity and (if pathogenic) mechanisms. Prior biophysical studies of the K^+^ channel PGD turret have highlighted its roles in display of negative electrostatic surface charge, and its binding of pore-blocker venom toxins (Miller et al., 1985; MacKinnon and Yellen, 1990; Hille et al., 1999; Banerjee et al., 2013; Zhao et al., 2019). By examining and comparing recent models of KCNQ1, 2, and KCNQ4 generated by cryoelectron microscopy (cryoEM), we found evidence that the KCNQ2 G256 residue contributes to a previously unstudied KCNQ channel turret role, stabilizing the open selectivity filter from its extracellular side. We analyzed KCNQ2 G256W pathogenicity via a multilevel experimental program including expression in heterologous cells and Crispr/Cas9-generated knock-in mice. We also made mice with a neighboring frameshift variant, and directly compared this model of the milder SLFNE phenotype with G256W in vivo. Unlike many other Kv currents, KCNQ2 mediated M-currents are non-inactivating (Brown and Adams, 1980). Our results led us to conclude that the biological role of this distinctive I_M_ property--absence of inactivation--is central to the pathophysiology of PGD variants like G256W.

## Results

### Clinical and developmental history of individual 1 index child

A non-dysmorphic female was born at term in the United States to parents of one older well-developing child. The neonate was vigorous with Apgar scores of 7 and 9, weight of 3253 gm (51.3%), length of 48 cm (26.8%), and orbitofrontal head circumference of 34.5 cm, (69.7%). Epileptic seizures began 3 hours after birth, and included arching, facial flushing, head deviation, eye rolling, and upper extremity flexion with tensing. The mother reported events during the third trimester concerning for in utero seizures. The fetus would maintain a rigid posture for 20-30 min, and could not be repositioned, causing significant maternal pain. Each episode was followed by a period of decreased fetal movement. The newborn was treated with levetiracetam (20 mg/kg BID). As seizures persisted, the following day phenobarbital was added (2 x 10 mg/kg). Seizures continued, despite levetiracetam, phenobarbital, and subsequently, lorazepam. Brain magnetic resonance imaging and screening metabolic labs showed no abnormalities.

The neonate was transferred to a tertiary care hospital for further evaluation and management. Chromosomal microarray, mitochondrial genome, CSF metabolic studies, and sequencing of several individual genes (*ARX, STXBP1, PDHA1, SCN1A*) revealed no abnormalities. Despite antiseizure medication escalations, frequent generalized tonic and focal tonic seizures continued. Review of 158 hours of video-electroencephalography (V-EEG) obtained between age 5 and 21 days, showed 73 seizures where onset, evolution, and offset were individually discernable. There were 2 periods, lasting 15 and 24 min, where several seizures overlapped. Of the well-resolved seizure onsets, 31 (42.4%) were diffuse and bilateral; 42 (57.6%) were unilateral (20 left-sided). Figure 1 illustrates a seizure of bilateral onset where the initial ictal change was diffuse voltage attenuation (Figure 1A, red arrow), a feature described previously (Ronen *et al*., 1993). Marked postictal voltage amplitude attenuation followed 67 of 73 seizures (91.8%, Figure 1B; Figure 1–Figure supplement 1-Movie). Each period of postictal attenuation was followed by recovery periods with increased discontinuity lasting for up to 5 min after seizures, especially generalized ones (Figure 1–Figure supplement 1-Movie). The interictal EEG background was abnormal due to multifocal spikes, poor organization, and excessive discontinuity for age. Unlike in Ohtahara syndrome, however, the EEG showed sleep state dependence, and included some variability and periods of continuity (Figure 1–Figure supplement 2). The infant was discharged from the hospital on (mg/kg/day) phenobarbital (1.6), levetiracetam (51) and topiramate (9.3), still having multiple daily tonic seizures. At home (age four weeks), new seizure types emerged including myoclonic and epileptic spasms, and additional EEG monitoring showed electrodecrements and multifocal epileptiform activity. Although hypsarrhythmia was not seen, treatment with adrenocorticotropic hormone (ACTH, 120 units/m^2^/day) was attempted at one month of age. This was accompanied by reduced seizure frequency, followed by cessation of clinical seizures within several days. Sanger sequencing of *KCNQ2* at two months of age identified a novel heterozygous variant, c.766G>T (p.Gly256Trp), classified as of uncertain clinical significance. Parental testing was not performed. Beginning at three months of age, the three antiseizure medicines were reduced sequentially and stopped. The infant remained seizure-free and completed tapering by eight months of age. Over the next dozen years, she experienced infrequent convulsions (6 in total), some provoked by febrile illness or overnight travel.

**Figure 1.**
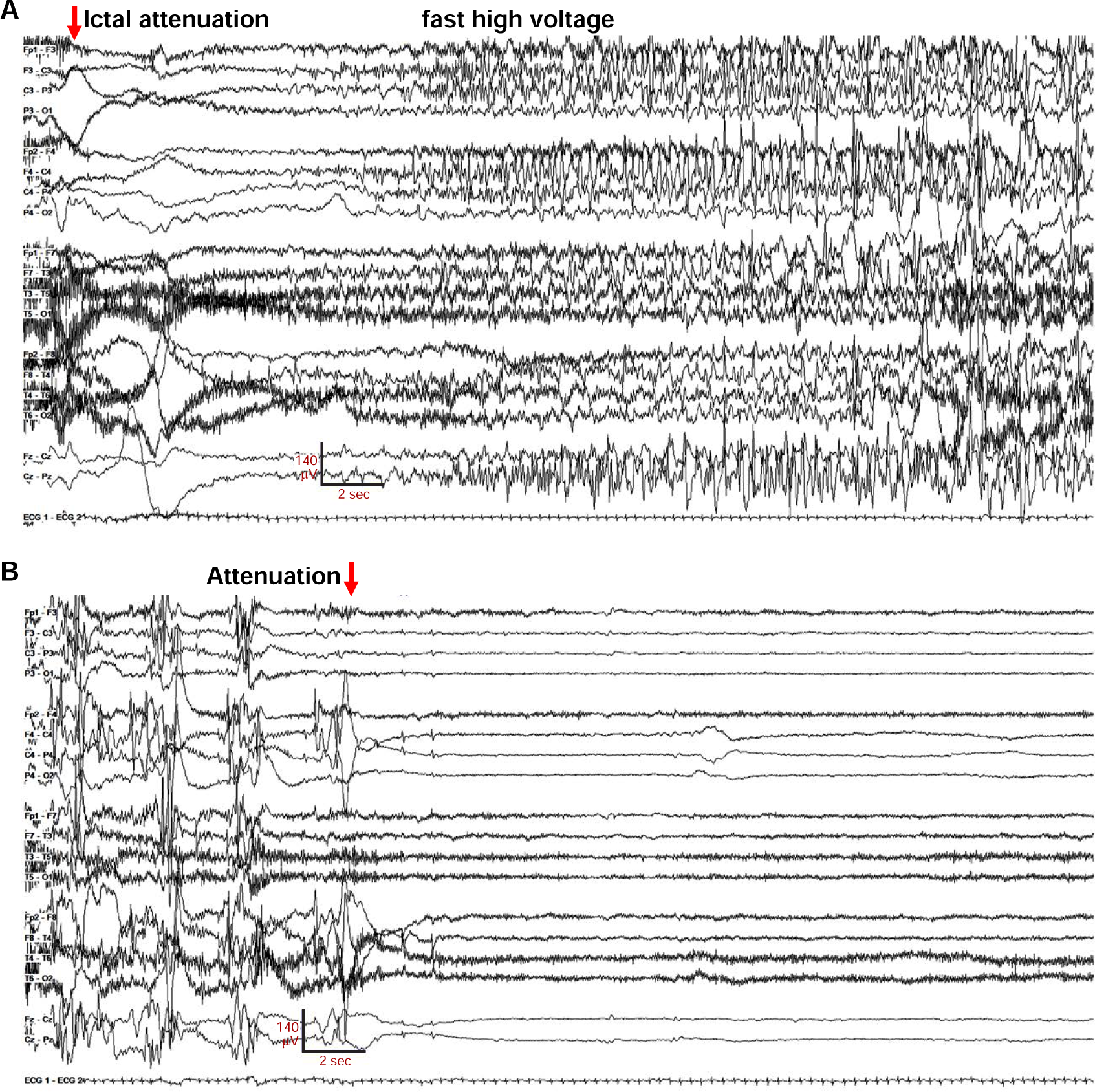
EEG of a bilateral onset seizure in Gly256Trp/+ individual 1, age 16 days. The recording is continuous from **A** to **B**. **A.** Seizure onset with diffuse, bilateral amplitude EEG attenuation (red arrow), which is obscured in several electrodes by high frequency muscle artifact (muscle artifact is better seen in Figure 1—figure supplement 1, movie). **B.** Seizure electrographic evolution to post-ictal voltage attenuation (red arrow). Settings: LFF 3 Hz, HFF 70 Hz, sensitivity 7uV/mm, 35 sec/panel.

Developmental delay was clinically apparent in infancy. Assessed by her neurologist at 9 months of age, she was functioning at a level expected typically at 5-6 month. She sat at 1 year, self-fed at 18 months, crawled at 20 months, and walked independently at 42 months. She received diagnoses of cortical visual impairment (1 year) and autism spectrum disorder (3 years). At last examination (age 12), she could use an adaptive communication device to request things (“eat sandwich”) and use modified sign language in 2-3 word combinations (“different music please”). She could respond to motor instructions, tap people when she wanted something, and point to items in a book. She is not able to manipulate clothing fasteners or descend stairs without supervision. Developmental challenges have included behavioral outbursts, self-injury, severe constipation, and sleep.

At 15 months of age, parents and physicians began a trial of ezogabine for potential beneficial effects on development. Within 2 weeks the parental global impression was of improved development, but no formal assessment was performed. An attempt to wean and discontinue ezogabine was made after the United States Food and Drug Administration published a notice warning of potential ezogabine-induced risks of skin discoloration, retinal abnormalities, and vision loss (FDA, 2013). Subsequent worsening of irritability, insomnia, and developmental skills with less ezogabine led caregivers to resume the prior dose. When the manufacturer subsequently announced plans for ezogabine market withdrawal, the drug was more slowly tapered and discontinued. A dilated ophthalmologic examination prior to ezogabine discontinuation was normal (29 month exposure).

### Individuals 2-4 with mosaic and heterozygous KCNQ2 G256W

A Korean collaborative team (Jang et al., 2019) described Individual 2, a child with neonatal seizures beginning on the second day of life and de novo *KCNQ2* c.766G>T (p.Gly256Trp). The seizure types included focal clonic, tonic, and epileptic spasms. When last seen at age 0.8 years, the child’s diagnosis was Ohtahara syndrome, but no additional information was presented. Very recently, a G256W occurrence was identified in Germany (NCBI, 2023). The proband male child (Individual 3) experienced seizures beginning at day of life 3 that remitted with antiseizure medicine. Medication was discontinued at one year of age but was resumed in childhood for infrequent convulsions. At last contact (age 12 years), he had delay in fine motor, gross motor, and language development, stereotypies, and was diagnosed with autism. One parent (Individual 4) had a history of seizures confined to the infantile period, never received antiseizure medication, and experienced subsequent normal development. Sequencing of Individual 3 at age 21 mo revealed *KCNQ2* c.766G>T (p.Gly256Trp) heterozygosity. The parent’s test showed the same variant in 320 of 831 reads (38.5%), indicative of post-zygotic mosaicism (Weckhuysen *et al*., 2013; Myers et al., 2018). These G256W recurrences in individuals manifesting characteristic *KCNQ2* DEE findings provide clinical evidence further supporting pathogenicity. The clinical information highlights electroclinical features we have modelled experimentally.

### G256W lies atop a dome-shaped hydrogen bond network linking helix S5 to the turret and selectivity filter

The KCNQ2 pore domain contains many known pathogenic missense variants and very few non-pathogenic missense variants, leading to low regional mutational tolerance (Traynelis et al., 2017; Karczewski et al., 2020). G256 lies near the beginning of the loop between the S5 and S6 transmembrane helices (Figure 2A). Seeking structural insights into pathogenic mechanisms of the heterozygous (i.e., G256W/+) substitution, we analyzed P-loop regions of cryoelectron microscopic structures from KCNQ2 and several homologues (Sun and MacKinnon, 2020; Li et al., 2021; Zheng et al., 2022). Like other K^+^ channels, the KCNQ2 P-loop has three distinct subsegments: the turret, the pore helix that partially penetrates the membrane, and the selectivity filter (SF, Figure 2A-E). Canonical SF residues T(I/V)GYG line the ion path perpendicular to the membrane surface. The SF polypeptide bends to parallel the membrane, forming a segment (herein termed the SF bridge) that extends radially from the ion path to S6 (Doyle et al., 1998; Hoshi and Armstrong, 2013). G256 faces the extracellular aqueous environment at the periphery and apex of the PGD, over 22 Å away from the SF (Figure 2E-F, Figure 2—Figure supplement 1-Movie). We wondered how substitution at G256 might alter pore function. We compared structural models of KCNQ2, KCNQ4, and KCNQ1 with those of the more distantly related channels KcsA, fly *Shaker,* and human Kv1.2 and made phylogenetic sequence comparisons (Figure 2—Figure supplement 2). Unlike in KCNQ1, KcsA, *Shaker*, or Kv1.2, KCNQ2 structural models and their density maps revealed G256 as the apical node of an H-bond network arching from S5 to the KCNQ2 SFB segment (Figure 2G-I, Figure 2—Figure supplement 3-Movie). Glycine uniquely confers main chain flexibility due to its lack of steric hindrance (Carugo and Djinovic-Carugo, 2013). KCNQ2 G256 contributes to a tight peptide turn through torsion angles (psi -64.4° and phi -74.3°) rarely found at non-Gly residues. The model predicts a Gly256-Glu257 ω-bond deviates from planarity, which is unusual (MacArthur and Thornton, 1996), contributing to a G256 carbonyl to N258 amide H-bond (Figure 2G-I). The N258 side chain extends away from G256 towards the SFB, where it contributes to a network of bonds including three turret (N258, H260 and D262) and three SFB residues (K283, Y284, and Q286).

**Figure 2.**
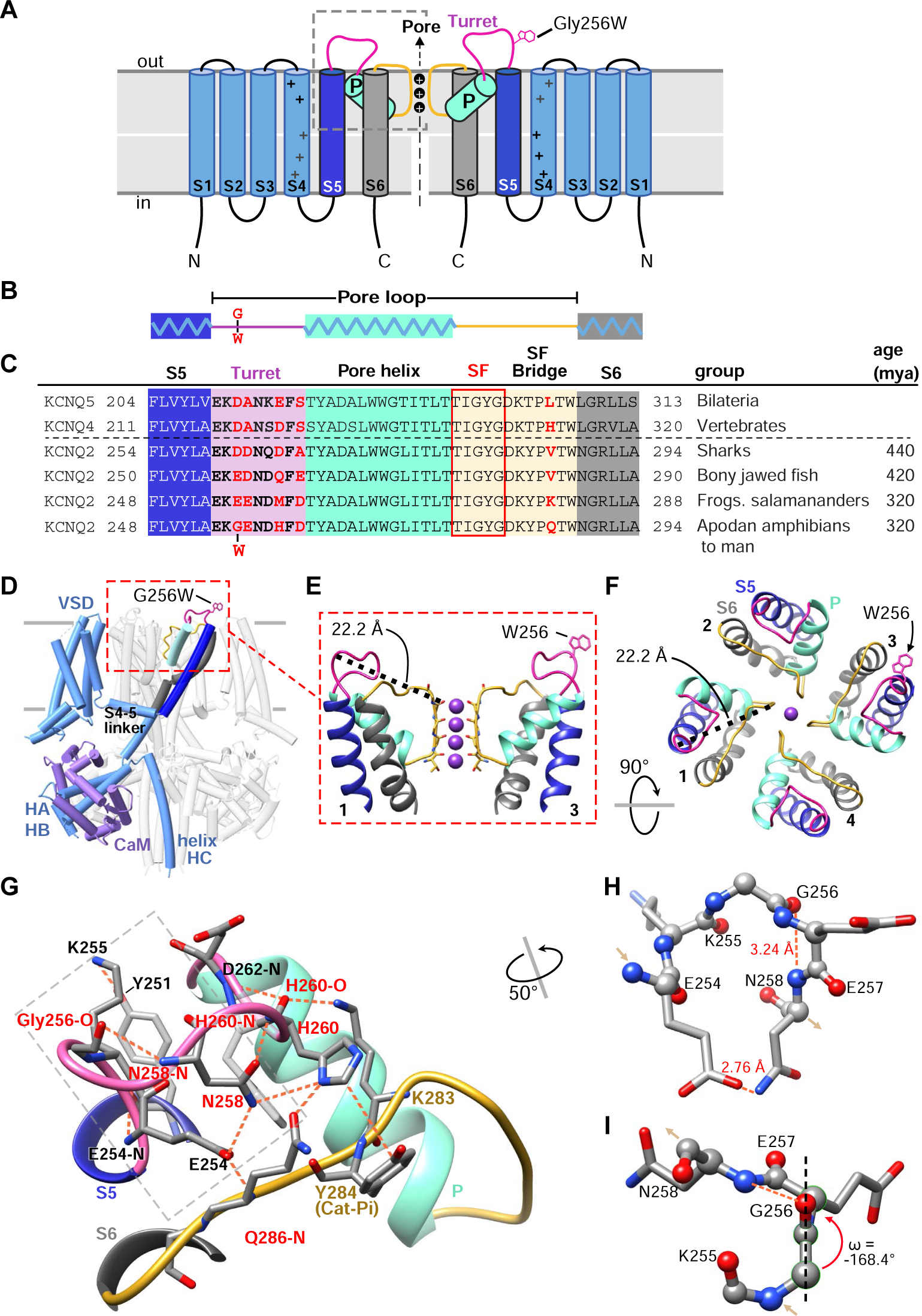
Gly256 is linked to the selectivity filter bridge segment via a hydrogen bond network among residues distinct to KCNQ2. **A, B**. Cartoons showing KCNQ2 membrane topology, including transmembrane segments S1-S6 and the P-loop (turret segment, purple; H5 or P-helix, cyan; and selectivity filter segment, yellow). Positions of the K^+^ selective pore, and the G256W substitution within the turret are indicated. **C**. Alignment of human KCNQ4 and KCNQ5 sequences with KCNQ2 sequences of major vertebrate groups. Background colors match panels A-B, and the five selectivity filter lining residues are boxed in red. At four aligned positions within the turret and one in the SFB, KCNQ2 substitutions have evolved in amphibians and tetrapods (residues highlighted in red). **D**. Rendering of the WT KCNQ2-calmodulin tetrameric structure obtained by cryoEM (PDB 7cr3), highlighting one subunit and the position of the G256W substitution near the channel’s extracellular domain apex. The Trp256 sidechain is at scale but its rotamer is chosen arbitrarily. The subunit closest to the viewer is partially deleted to reveal the highlighted subunit more clearly. **E**. Ribbon rendering of the extracellular part of the PGD. For clarity, only two opposing side subunits are shown (as schematically in **A**). A Trp side chain is added at one Gly256 α-carbon. The distance between the G256 α-carbon and Y280 carbonyl oxygen at the selectivity filter mouth is labeled. **F**. Top down view of the KCNQ2 regions as in panel **E**, but showing 4 subunits. The Trp rotamer is different from panels **D-E**. The S5, S6 and P-helices are labeled. **G**. Hydrogen bonding network of the KCNQ2 turret. All predicted bonds are shown as dashed orange lines. The network extends from the S5 helix (Y251) via the labelled turret residue atoms to bonds involving residues of the SFB. As in **C**, five residues that diverge invertebrates are colored red. **H, I.** The turret peptide region near G256, which is boxed with a grey dashed line in **G**. The main chain is shown as ball-and-stick; side chains as stick. A tight turn occurs at K255 to N258, stabilized by hydrogen bonding between the G256 carbonyl oxygen and N258 amide. **I.** The G256-E257 peptide deviates from planarity (ω = +/-180°) by 11.6° (∼2.6 sd). In and out arrows indicate N and C termini, respectively. Abbreviations: mya, million years ago; VSD, voltage-sensor domain; HA-HB, the cytoplasmic helices A and B; CaM, calmodulin.

Phylogenetic comparisons indicate that four turret and SFB residues (G256, H260, D262 and Q286) within this bond network co-evolved in *KCNQ2* genes of fish and amphibians, and were subsequently conserved across reptiles, birds, and mammals, divergent from other *KCNQ* subtypes (Figure 2C). KCNQ4, an evolutionary ancestor of both KCNQ2 and KCNQ3 (Cooper, 2011), exhibits a very similar turret fold and a bond network linking S5, turret, and SFB (Figure 2— Figure supplement 2C-D). Turret sequences and structures of KCNQ1,KcsA, *Shaker,* and Kv1.2 are less like KCNQ2 and lack direct bonds to the SFB. The turret sequence of KCNQ3, the most recently arising KCNQ2 paralogue, includes 10 inserted residues that are absent from KCNQ5, 4, and 2 (Figure 2—Figure supplement 2F). This evidence of unique divergence and later conservation suggested that despite its small side chain, water-facing location, and distance from the pore, KCNQ2 G256 could be intolerant to substitution. Beyond the loss of unique Gly physicochemical features, introducing a large, planar, hydrophobic W256 side chain might cause disruptive local consequences due to its preference for burial at membrane boundaries (Khemaissa et al., 2022). We tested these predictions with experiments in vitro and using mice bearing the heterozygous G256W substitution.

### G256W co-expression suppresses currents of KCNQ2/KCNQ3 channels in Chinese hamster ovary (CHO) cells

We made whole cell patch clamp recordings from CHO cells co-expressing wild-type (WT) KCNQ2 and KCNQ3 using an automated, 384-well system (Vanoye et al., 2022). We compared three expression conditions: WT KCNQ2 and KCNQ3, G256W and KCNQ3 WT, and co-expression of a 1:1 ratio of WT KCNQ2 and G256W with WT KCNQ3 to mimic the heterozygous genotype. A simplified random association model assuming equal expression and assembly predicts that the last condition results in a mixed population of subunit stoichiometries containing 0, 1, or 2 G256W subunits (Figure 3A). The WT channels gave currents with slow voltage-dependent activation and no inactivation (Figure 3B). Cells expressing either KCNQ3 only (Vanoye *et al*., 2022), or the combination of KCNQ2 G256W and WT KCNQ3 (Figure 3B) exhibited no detectable currents. When WT KCNQ2 and G256W cDNAs were co-expressed at a 1:1 ratio to mimic heterozygosity, currents at 40 mV were significantly reduced to 44.3 ± 8% compared to WT only controls (Figure 3B-C). This reduction appeared linear with respect to time and voltage, as conductance-voltage fits showed no significant changes in V_1/2_ (Figure 3E), activation slope, or time constants.

**Figure 3.**
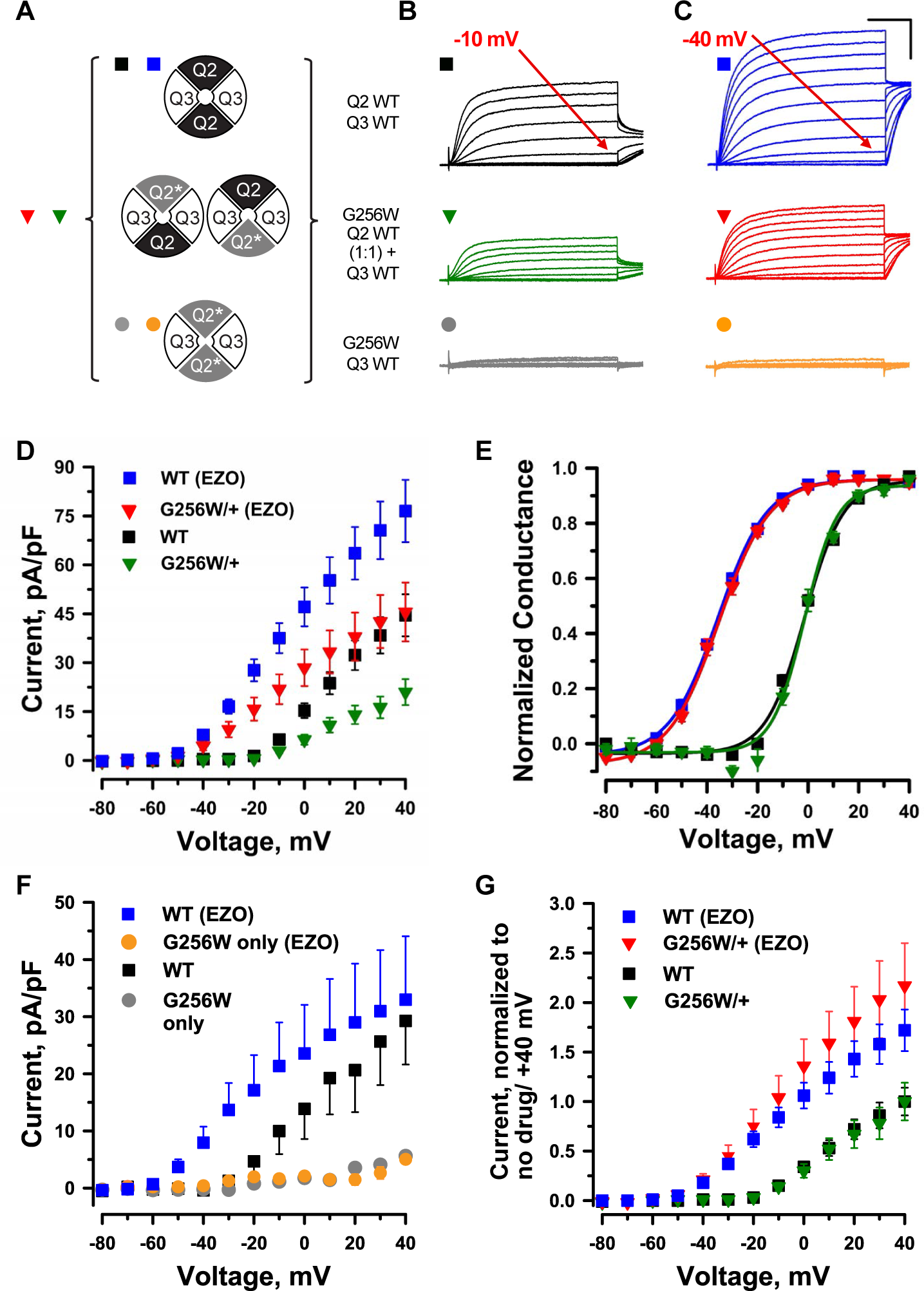
KCNQ2 G256W co-expression suppresses current in KCNQ2/KCNQ3 heteromeric channels. **A.** Cartoon showing the expected combinations of WT and G256W subunits under heterozygosity based on a simple random association model and preferred 2:2 stoichiometry for KCNQ2 and KCNQ3. **B-G.** In vitro dissection of effects of G256W heterozygosity on currents. **B-C.** Mean current families are shown for the indicated combinations of expression of KCNQ2 and KCNQ3 prior to and after addition of 10 μM ezogabine (n = 60, 50; 40, 31; 28, 24 for the upper, middle, and lower conditions). Scale**D-E.** Current/voltage and conductance/voltage relationships for the indicated WT only and G256W/WT electroporations into KCNQ3 stable expressing cells. **F.** Current/voltage relationship for G256W (‘homozygous”) heteromeric channels, compared with the subset of WT control cells studied in parallel by automated patch recording. **G.** Replot of data from panel **D**. At each voltage, mean current is normalized to mean current at +40 mV in absence of ezogabine.

Ezogabine (retigabine) increases currents through neuronal KCNQ channels by shifting activation to more hyperpolarized voltages, enhancing activation kinetics, and increasing maximal current density (Tatulian et al., 2001; Gunthorpe et al., 2012). During its period of commercial availability, ezogabine was used as targeted therapy in patients with *KCNQ2* DEE arising from loss-of-function variants (Millichap and Cooper, 2012; Weckhuysen *et al*., 2013; Millichap *et al*., 2016; Nissenkorn et al., 2021), including in individual 1. We compared the effects of ezogabine (10 µM) on WT KCNQ2/KCNQ3 heteromers and channels from cells expressing G256W. In cells expressing G256W and KCNQ3, ezogabine treatment had no effect (Figure 3C, F). In cells expressing WT KCNQ2, G256W and WT KCNQ3, mimicking heterozygosity and predicted to yield a mixed population of channels (Figure 3A), ezogabine significantly increased currents and shifted activation voltage dependence (Figure 3C, D, E). Currents from the heterozygous G256W condition after ezogabine exceeded those of WT KCNQ2/KCNQ3 control cells absent ezogabine. We compared the relative effect of ezogabine in control and G256/+ cells by dividing post-by pre-treatment currents. Ezogabine enhancement in cells mimicking theG256W/+ genotype was equal to that of WT controls (Figure 3G). This is especially notable, as 25% of the channels assembled in the G256W/+ mimicking cells are predicted to include 2 G256W subunits and therefore be ezogabine-unresponsive. This suggested ezogabine treatment increased current in cells including one G256W subunit as fully as in WT KCNQ2/KCNQ3 channels.

### G256W co-expression suppresses homomeric KCNQ2 currents in Chinese hamster ovary (CHO) cells

In parallel, we studied the effect of G256W through manual whole-cell patch recordings. We used a co-transfected GFP marker to select for high expression cells for patching. Mean current densities were higher than observed for the automated patch recording. However, as in the automated patch system, co-expression of G256W with WT KCNQ3 resulted in little or no current. Co-transfection of G256W and WT KCNQ2 cDNA with KCNQ3 (1:1:2) to mimic heterozygosity reduced current to 41% +/- 0.1 % of WT controls (Figure 3—Figure supplement 1). Because KCNQ2 subunits appear to be expressed as KCNQ2 homomeric channels in some neurons (Cooper et al., 2000; Hadley et al., 2003; Martire et al., 2004; Schwarz et al., 2006; Varghese et al., 2023) we analyzed the impact of G256W in this subunit configuration. WT KCNQ2 expressed alone gave robust currents, but G256W currents were undetectable (Figure 3—Figure supplement 2). Co-expression of WT KCNQ2 and G256W cDNA (1:1) has been modeled as yielding a mix of tetramers including WT subunits only, G256W only, and between 1 to 3 G256W subunits, with the ratios of these stoichiometries predicted by the binomial distribution (Figure 3—Figure supplement 2, panel D). The current density (+40 mV) under 1:1 co-expression to mimic heterozygosity was 30.1% +/- 0.056 % of WT control, a greater relative reduction than seen for heteromeric channels, as predicted by with the assembly models. Heterozygous co-expression of G256W did not change the midpoint voltage or steepness of voltage-dependent activation in either the heteromeric or homomeric subunit composition experiments.

### A neonatal seizure in a heterozygous G256W mouse at P10

To better understand the consequences of the G256W/+ expression in vivo under control of the native *Kcnq2* promoter, we introduced the variant into C57BL6/J mice using Crispr/Cas9. Initial progeny included mice heterozygous for the intended variant (G256W/+) and a mouse heterozygous for a 7 base deletion in codons 254-256 (E254*fs*; Figure 4). The deletion preserved the WT exon 5-6 splice boundary (Figure 4—Figure supplement 1), yielding a transcript with 15 novel sense codons and an in-frame stop codon spanning the exon 5-6 junction. The E254*fs* transcript is predicted to yield a truncated protein and be targeted for nonsense mediated mRNA decay., Western blots using antibodies against the KCNQ2 N-terminus did not reveal bands for the protein product (Figure 4—Figure supplement 2). Truncating variants are found in a majority of SLFNE families (59.5%; Goto *et al*., 2019). To enable direct comparison between a characteristic SLFNE variant type and G256W in vivo, we purified both lines by breeding against WT C57BL6/J and studied them in parallel.

**Figure 4.**
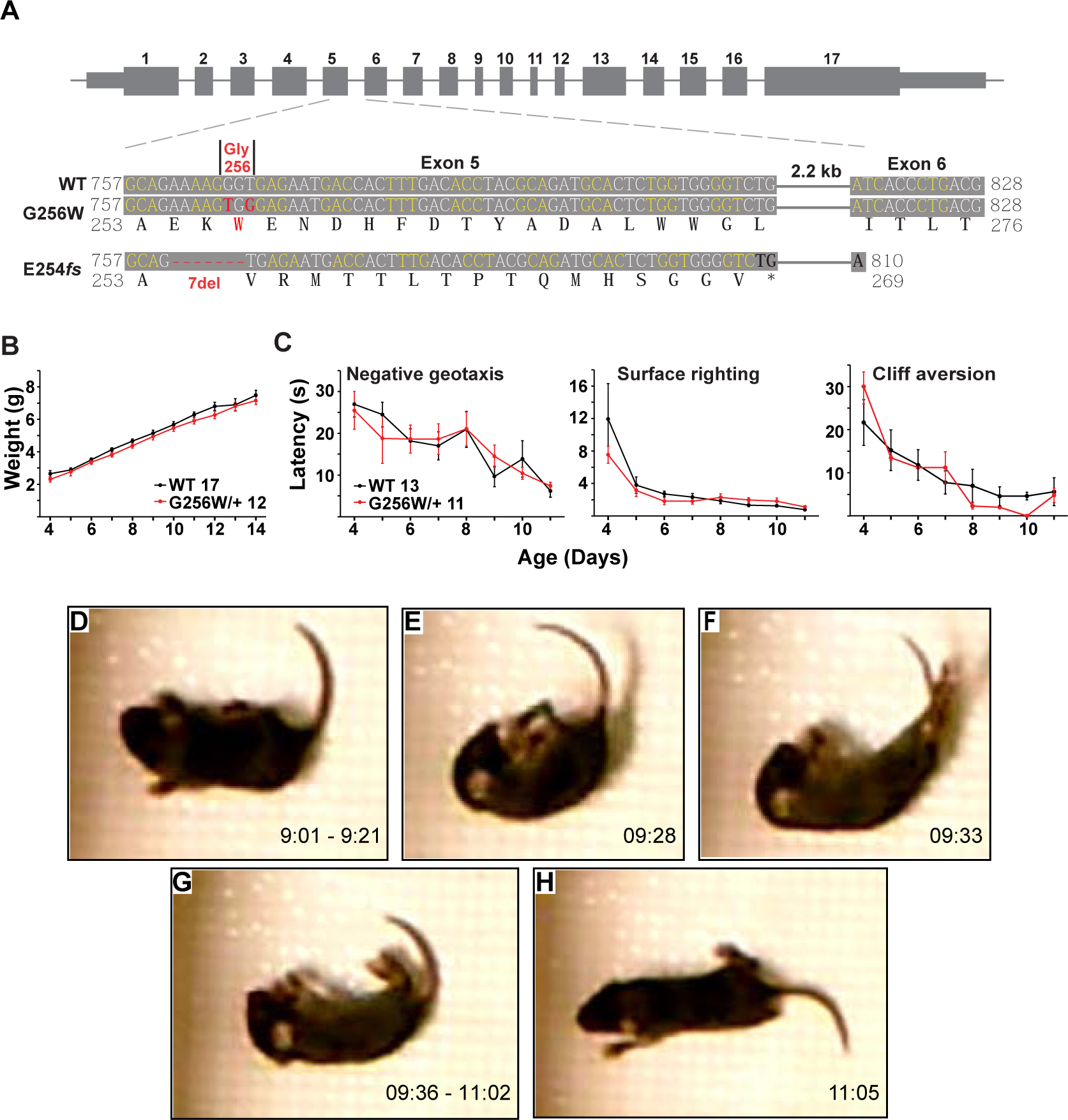
Immature heterozygous G256W mice exhibit normal development and have infrequent epileptic seizures. **A.** Upper, map of the *Kcnq2* constructs. Lower, sequence alignments for the region between the middle of exon 5 and the beginning of exon 6. Although the human G256W variant is a single base substitution, Crispr/Cas9 editing introduced two substitutions, since the WT G256 codons differ between mouse (GGT) and human (GGG). Also aligned is the DNA and protein sequences of the frameshift mutation. **B.** WT and G256W/+ mice showed no difference in weight gain during development. **C.** WT and G256W/+ mice performed similarly in the developmental milestone assays for negative geotaxis, surface righting, and cliff aversion. **D-H.** Screenshots of stages of a generalized seizure in a P10 G256W/+ mouse (see also: Figure 4**—**Figure supplement 3-Movie). **D.** Onset with immobility and myoclonic tail and forelimb shaking. **E.** Abrupt fall to side with flexion posturing. **F.** Evolution to hindlimb and tail extension posture. **G.** Immobility with flaccid appearance, interrupted by brief episodes of tail, individual limb myoclonus or clonus. **H.** Arouses, quickly regains upright posture, then normal mobility. Labels: time in 15 min source video.

In crosses of WT females and heterozygous males, G256W/+ and E254*fs*/+ mice represented ∼50% of live births (230/481 and 147/309, respectively). For E254*fs* het x het crosses, 12 WT, 33 E254*fs*/+, and 3 E254*fs*/E254*fs* mice were born. The E254*fs*/E254*fs* pups appeared stillborn or died at P0, as seen for homozygotes with other *Kcnq2* null alleles (Watanabe et al., 2000; Yang et al., 2003). For G256W het x het crosses, 16 WT, 24 G256W/+, and 5 G256W/G256W mice were born. All G256W/G256W pups were dead at initial observation on P0 or died within hours. G256W/+ pups showed no differences in weight nor in the screens for progress in several assays of motor development (Figure 4B-C). However, we video recorded a convulsive seizure in a P10 G256W/+ female (Figure 4D-H). The seizure evolved from behavioral arrest and myoclonic jerks, followed by loss of postural control and hindlimb/tail extensor posturing. After 90 sec of immobility with brief episodes of myoclonus, the mouse regained awareness and upright posture. The seizure lasted approximately 120 seconds in total (Figure 4—Figure supplement 3-Movie). Between five and nine G256W/+ mice and 26 control mice were video recorded on alternate days between P8 and P14 for 15 min per day, and only one seizure was observed.

### CA1 pyramidal neurons in P12-15 heterozygous G256W mice show increased firing and reduced spike frequency adaptation

We made whole cell recordings of CA1 pyramidal cells in acute horizontal hippocampal slices from P12-15 WT and G256W/+ littermates. Compared to WT neurons, positive current injections in G256W/+ neurons evoked significantly greater numbers of action potentials (Figure 5). There were no differences in resting membrane potential, several action potential biophysical parameters, or somatic input resistance between WT and G256W/+ mice (Figure 5—Figure supplement 1).

**Figure 5.**
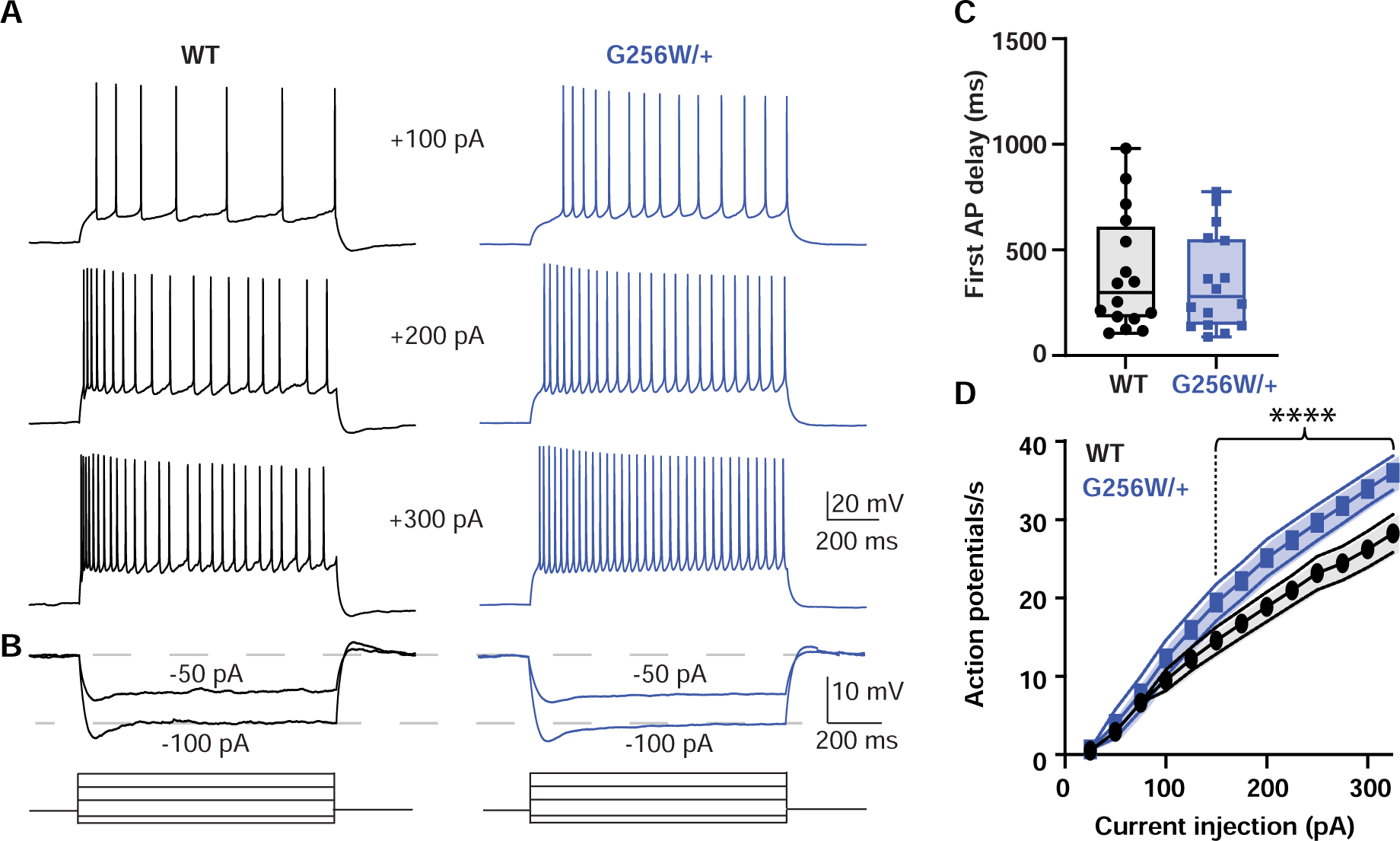
Heterozygous G256W mice have increased CA1 pyramidal cell excitability. **A.** Representative voltage responses to increasing current injection steps (step duration 1 sec) in CA1 pyramidal neurons from WT and G256W/+ mice. The resting membrane potential was held at -65 mV. **B.** Representative voltage responses to decreasing current injections steps (1s) in CA1 pyramidal neurons from WT and G256W/+ mice. **C**. Time to first action potential following step stimulus is not significantly different between groups (3 animals per group; WT and G256W/+, n=16 cells each).**D.** Summary graph showing the effect of one copy of G256W on the action potential count (3 animals per group; WT and G256W/+, n=16 cells each, F(12,180)=5.8, **** is P<0.0001).. Data are presented as mean and s.e.m.

### Adult heterozygous G256W mice experience fatal and nonfatal seizures

Convulsive seizures were observed in adult G256W/+ mice occasionally during routine animal care but never in co-housed WT littermates. To learn electrophysiological correlates, we performed electrocorticography on eight adult heterozygous mice for a total of 1440 hours. The five electrographic seizures captured in two mice were stereotyped. All exhibited generalized onset and stereotyped evolution: an isolated herald spike or polyspike (Figure 6A and 6B, arrows), rapid transition to fast, high-amplitude spiking lasting 20-30 seconds, and strong amplitude attenuation lasting 15-30 seconds, and slow recovery (Figure 6A). One mouse was recorded for 150 hours over 10 days without seizures, then had four seizures within 72 hours, the last of which was fatal. The final ictal EEG followed the pattern of prior seizures, but the strongly attenuated EEG diminished progressively and did not recover (Figure 6B-C). One fatal seizure was video recorded (Figure 6—Figure supplement 1-Movie). In this mouse, seizure began with wild running and jumping (for 5-10 sec), followed by a brief arrest (and start of the video recording). Running resumed for 3 sec, followed by abrupt loss of postural control, extensor posturing, and respiratory arrest. The experimenter attempted cardiac resuscitation with anterior chest compressions, but the mouse did not revive. In other seizure instances, mice were observed to revive, either without intervention or with chest compressions or a noxious limb pinch stimulus. Kaplan-Meier analysis showed that ∼22% of non-censored G256W/+ mice died prematurely, a significant increase compared to WT littermates, with a median age of death of 180 days (Figure 6D). E254*fs*/+ mice experienced no early mortality, in agreement with findings for other SLFNE models (Watanabe *et al*., 2000; Singh et al., 2008; Robbins et al., 2013; Figure 6—Figure supplement 2). Many G256W/+ premature deaths during long-term colony housing were unwitnessed, but the mouse bodies we recovered after death exhibited the stereotyped posture we observed in the two seizure videos, with flexed forelimb and hindlimb extension posture captured by rigor mortis.

**Figure 6.**
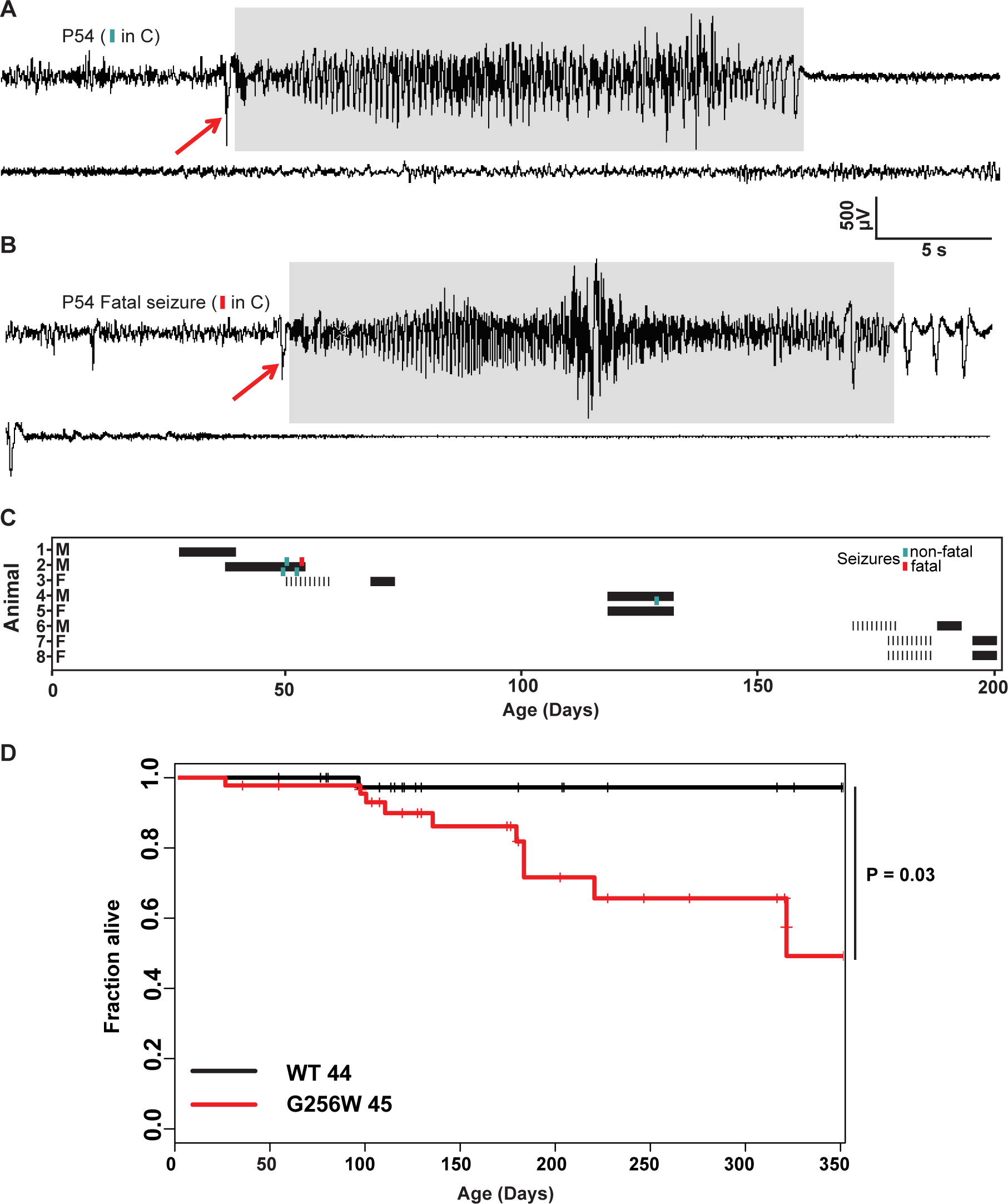
Convulsive seizures in adult heterozygous G256W mice show stereotyped electrographic features and reduce survival. **A-B.** EEGs of non-fatal and subsequent fatal seizure captured in a P54 male G256W/+ mouse (animal 2, panel C). Electrographic seizures were characterized by fast spiking, high amplitude activity lasting 15-20 s (highlighted in gray). **C.** Summary showing the sex, ages, duration of recordings and timing of seizures in 8 animals undergoing EEG. Turqoise hashmarks denote a survived seizure, red hashmark denote a fatal seizure. Black bars are periods on EEG; some recording were performed on a 6 hr/day schedule. **D.** Survival curve of WT vs G256W/+ mice, hashmarks indicate censored mice. G256W/+ mice had signifcant mortality, P = 0.0348 Cox propotional hazards model.

### Distinct patterns of altered *Kcnq2* and *Kcnq3* mRNA expression in heterozygous E254*fs* and G256W mice

We used RT-qPCR to measure *Kcnq2* and *Kcnq3* mRNA levels in hippocampus and neocortex at P21 and P100. There were no differences in *Kcnq2* mRNA expression between WT and G256W/+ mice (Figure 7A). Because E254*fs* transcripts should be eliminated by nonsense mediated decay, we expected E254*fs*/+ mice to have about 50% less *Kcnq2* mRNA than WT and G256W/+ mice. However, using an assay that did not discriminate between transcripts from WT and E254*fs* alleles, *Kcnq2* mRNA was reduced by only 25% (+/- 3%) at P21 and by 35% (+/- 1%) at P100 in E254*fs*/+ mice. The *Kcnq2* mRNA levels in E254*fs*/+ mice were significantly greater than 50% of WT (one sample t-test). This result could reflect incompletely efficient nonsense mediated decay of E254*fs* transcripts (Zetoune et al., 2008; Dyle et al., 2020), or compensation achieved via increased *Kcnq2* gene expression or mRNA stability. To attempt to discriminate among those possibilities, we performed additional experiments using a TaqMan assay detecting only the WT *Kcnq2* allele (McGuigan et al., 2002). In this assay, E254*fs*/+ and G256W/+ mice both showed approximately 50% losses of mRNA. In E254*fs*/+ mice, total *Kcnq2* mRNA was significantly greater than WT *Kcnq2* mRNA (one sample t-test; Figure 7B). Taken together, these results suggest that nonsense mediated decay of E254*fs* transcripts is incomplete, and that expression of the *Kcnq2* WT allele is unchanged in both lines of heterozygous mice. In contrast, RT-qPCR revealed a significant increase in *Kcnq3* mRNA in E254*fs*/+ hippocampus at P21, as observed previously for *Kcnq3* mRNA *Kcnq2* homozygous null mouse models (Robbins *et al*., 2013). In addition, *Kcnq3* mRNA was increased in 3 of 4 G256W/+ experiments (P21 cortex, P21 hippocampus and P100 hippocampus but not P100 cortex). Consistent with the qPCR results, Sanger sequencing of cDNA amplified from P21 hippocampus of G256W/+ and E254*fs*/+ mice showed peaks corresponding to the G256W and E254*fs* alleles (Figure 7—Figure supplement 1, panel B-C).

**Figure 7.**
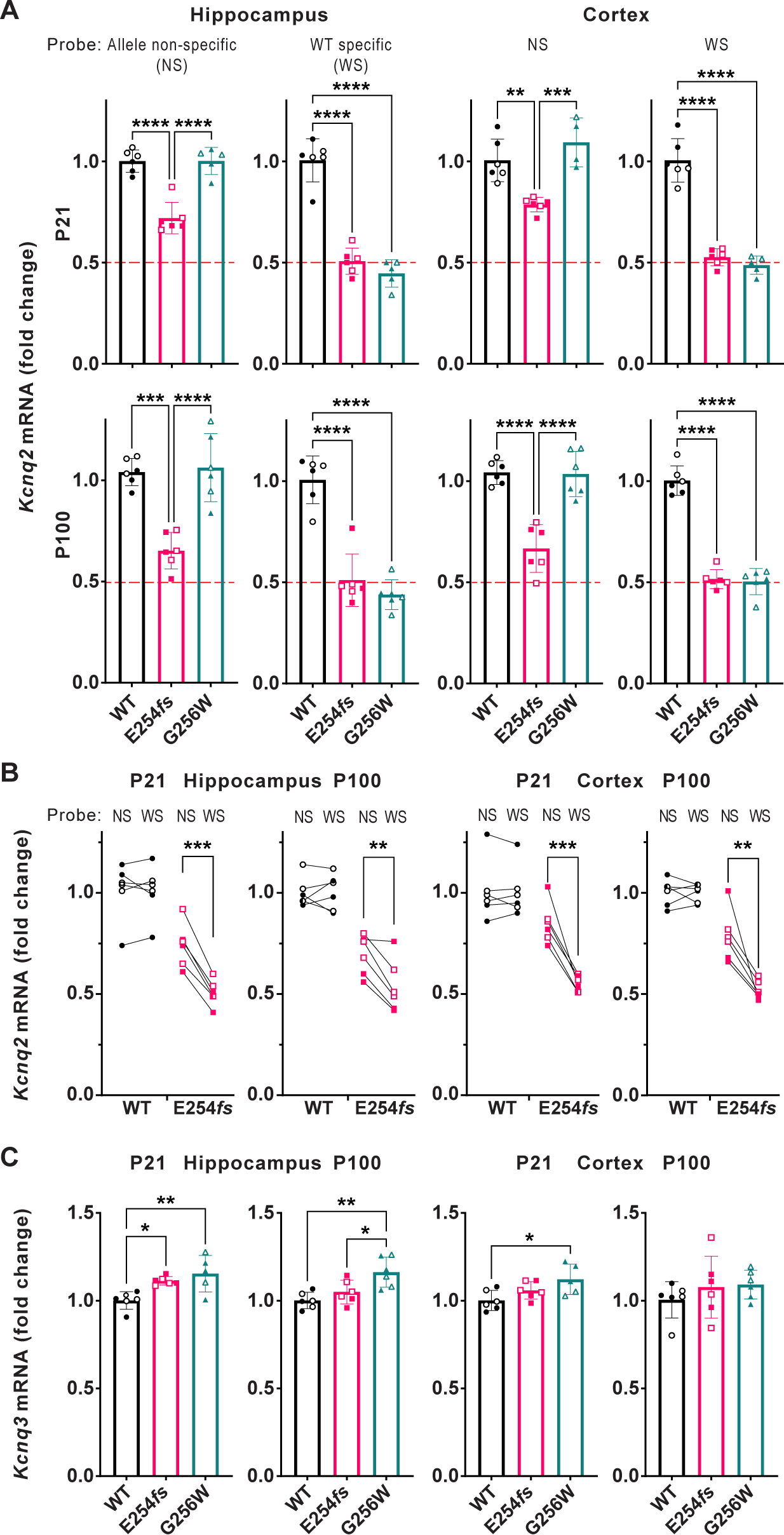
RT-qPCR shows incompletely efficient nonsense mediated decay of *Kcnq2* E254*fs/+* mRNA and increased *Kcnq3* mRNA in E254*fs/+* and G256W/+ mice. **A.** Upper, *Kcnq2* mRNA levels in P21 hippocampus and neocortex. Levels were analyzed using TaqMan probes binding to both WT and variant *Kcnq2* alleles (allele non-selective, NS), or binding only to WT only (WT selective, WS). Total *Kcnq2* mRNA in E254*fs/+* samples were significantly higher than 50% of WT (hippocampus: P = 1.28x10^-5^, neocortex: P = 3.5x10^-7^, one sample t-test). Lower, *Kcnq2* mRNA levels in P100 hippocampus and neocortex, using the probes as above. Total *Kcnq2* mRNA levels in E254*fs/+* samples were significantly higher than the expected 50% of WT in hippocampus (p = 0.0003) and neocortex: (p = 0.0007). **B.** Total and WT *Kcnq2* mRNA, tested in parallel, by individual. Age, sex (male, open symbols), brain region, genotype, and probe are indicated. In all four tissues tested, E254*fs/+* mice have greater total *Kcnq2* mRNA than WT *Kcnq2* mRNA (P21 hippocampus, P = 0.0001; P100 hippocampus, P = 0.005; P21 neocortex, P = 0.0007; P100 neocortex, P = 0.0053; pairwise t-test). **C.** In P21 G256W/+ mice, *Kcnq3* mRNA was significantly increased: 1.15-fold (+/- 0.10, P = 0.0043) in the hippocampus and 1.12-fold (+/- 0.09, P = 0.00213) in the neocortex. In P100 E254*fs*/+ mice, *Kcnq3* mRNA significantly increased (1.11 +/- 0.02-fold, P = 0.0245) in the hippocampus only. One way ANOVA, * = p≤0.05, ** = p≤0.005, *** = p≤0.0005. (See Supplemental Data for statistical test calculations).

### Heterozygous G256W mice show diminished KCNQ2 and KCNQ3 targeting to axon initial segments and axons, and reduced levels of KCNQ2 protein

KCNQ2 is highly concentrated at many axon initial segments and nodes of Ranvier, where it colocalizes with voltage-gated sodium (NaV) channels, Ankyrin-G, and other Ankyrin-G interacting proteins (Devaux et al., 2004; Pan et al., 2006). In many but not all neurons, KCNQ3 is also colocalized at these subdomains and forms heteromeric channel with KCNQ2 (Shah et al., 2008; Klinger et al., 2011; Battefeld et al., 2014; Martinello et al., 2015; Jing et al., 2022). We compared KCNQ2 and KCNQ3 subcellular localization in WT, E254*fs*/+ and G256W/+ mice by immunofluorescence labeling and confocal imaging of hippocampal sections, using antibodies directed against sequence from the KCNQ2 N-terminal region and pairs of samples processed and imaged in parallel. In CA1, where abundant pyramidal cell AISs are found crossing the border between stratum pyramidale and stratum oriens (Figure 8-Movie), sections from G256W/+ mice showed diminished AIS labeling and increased neuronal somata labeling for both KCNQ2 and KCNQ3. In the CA1 of E254*fs*/+ mice, the intensity ratios of AIS and somatic labeled regions did not differ from WT (Figure 8—Figure supplement 1-Movie). In CA3, images of G256W/+ mice showed less intense KCNQ2 and KCNQ3 labeling within stratum lucidum (where mossy fibers are marked by PanNav) and increased labeling in stratum pyramidale (Figure 8—Figure supplement 2-Movie).

**Figure 8.**
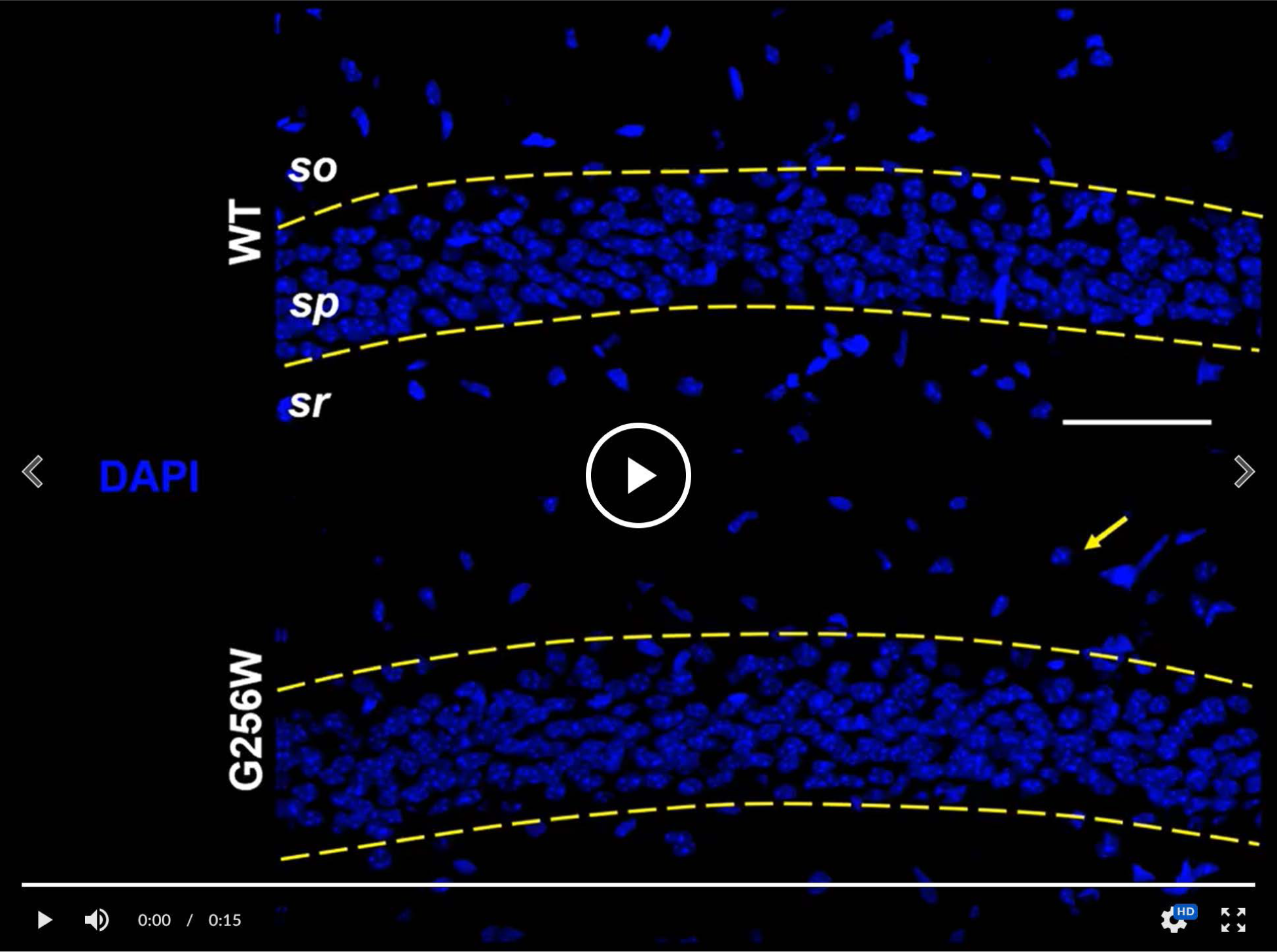
Heterozygous G256W mice show reduced KCNQ2 and KCNQ3 labeling of CA1 pyramidal cell AISs and increased labeling of neuronal somata. Identically processed age P21 tissue sections of WT (upper) and G256W/+ (lower) mice; area CA1B was imaged under identical settings. Confocal image stacks are shown as maximal intensity projections. In the animation, channels for the indicated markers are allowed to fade into the next, enabling evaluation of colabeling. DAPI marks cell nuclei. AnkG strongly marks AISs and lightly labels somata and proximal apical dendrites. An arrow highlights one stratum oriens interneuron somatically labeled for KCNQ2 only. Labels: DAPI, 4’,6-diamidine-2’-phenylindole; so, Stratum oriens; sp, Stratum pyramidale, sr, Stratum radiatum. Scale: 50 μm. <u>Link to movie F8</u>

To test these results obtained by visual inspection of paired samples, we performed a quantitative, blinded analysis of subcellular labeling of WT, G256W/+, and E254*fs*/+ mouse hippocampal sections in CA1, CA3, and the dentate gyrus. We marked axonal and soma containing regions of interest (ROIs) and quantitated relative axon (or AIS) to somatic containing ROI intensities. Comparisons between these ratios reflect relative axonal (or AIS) and somatic protein concentrations. The approach does not quantify at the level of individual axons and somata, and linearity of labeling intensity vs. protein was not established. Because these limitations applied equally across genotypes, however, the methods allowed differences between the genotypes to be detected and tested for significance. For G256W/+ mice, the ratio of pyramidal cell AIS/somatic ROI intensity was significantly reduced for both KCNQ2 and KCNQ3 in the CA1 and CA3 (Figure 9A-B). In CA3, this was driven mainly by increased pyramidal cell somatic labeling, as the long CA3 AISs (Kosaka, 1980) are more sparse than in CA1 and were infrequently captured longitudinally in coronal sections. Also, G256W/+ mice had a significant reduction in the CA3 stratum lucidum to stratum pyramidale ROI ratio, suggesting potential diminished forward trafficking to the mossy fibers and/or CA3 pyramidal cell axon collaterals (Figure 9C). The very thin AISs of dentate granule cells lie mostly within the granule cell layer (GCL; Martinello *et al*., 2015), so measurement of ratios between tissue layers is an imperfect proxy for (soma vs. AIS) subcellular distribution. The KCNQ2 labeling ratio for dentate PML (axon-enriched) vs. GCL (somata and AISs) was not significant different between genotypes, but the KCNQ3 ratio showed significant reduction (Figure 9D).

**Figure 9.**
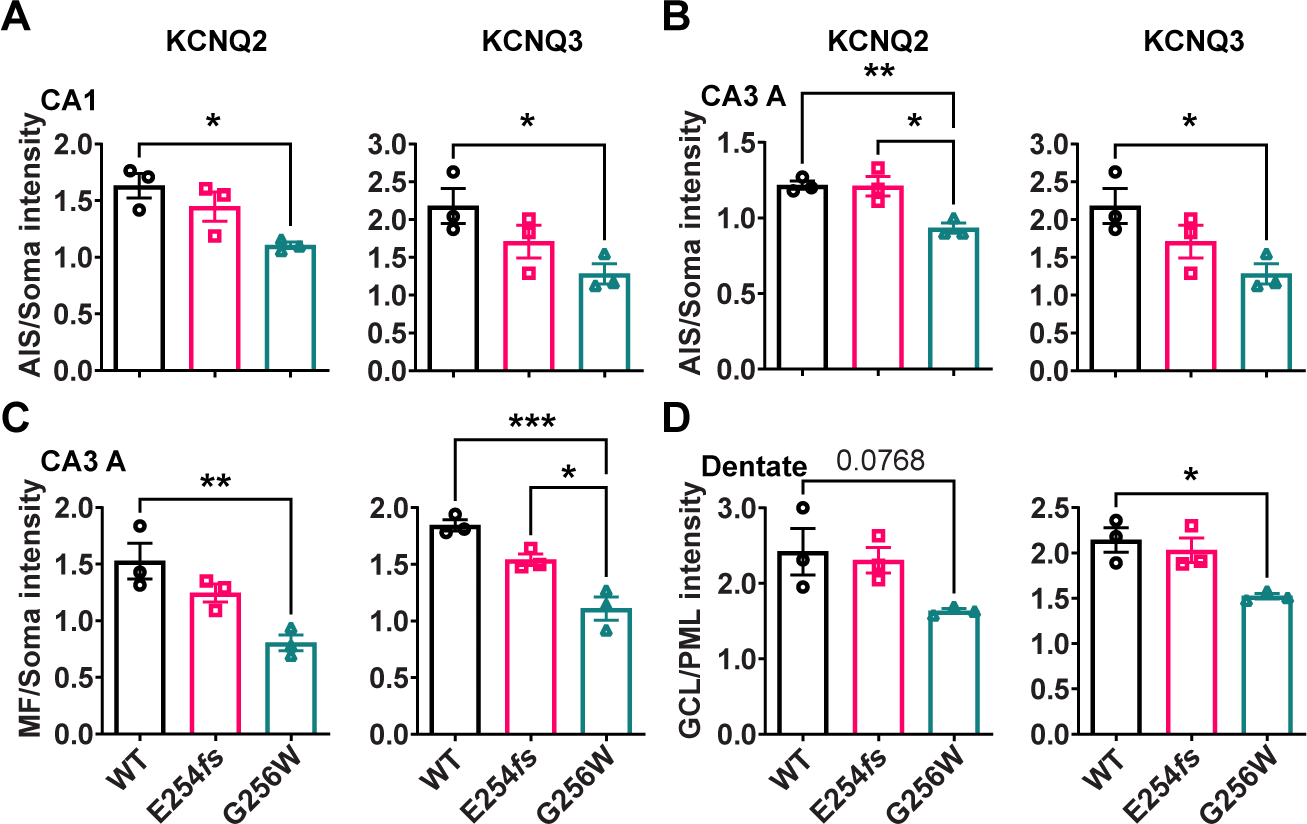
The ratios of axonal to somatic KCNQ2 and KCNQ3 labeling are reduced in CA1 and CA3 in heterozygous G256W mice. **A-B.** The ratios of AIS to somatic immunofluorescence intensity is significantly reduced for KCNQ2 and KCNQ3 in CA1 (**A**) and CA3 (**B**) for G256W/+ but not E254*fs*/+ mice. **C.** The ratio of mossy fiber to somatic KCNQ2 and KCNQ3 immunofluorescence intensity is reduced in the CA3 for G256W/+ but not E254*fs*/+ mice. **D.** In the dentate gyrus, the ratio between GCL and PML intensity is significantly reduced for KCNQ3 but not KCNQ2 in G256W/+ but not E254*fs*/+ mice. n=3 per genotype. One way ANOVA, * = P≤0.05, ** = P≤0.005, *** = P≤0.0005.

In the dorsal hippocampal CA1 region, pyramidal cells outnumber interneurons about 100-fold (Keller et al., 2018). We did not investigate labeling of the hippocampal interneurons extensively, due to their less frequent appearance in the matched sections used for analysis of excitatory neurons. In six CA1 image stacks of G256W/+ mice, 11 putative interneurons showed conspicuous KCNQ2 somatic labeling (e.g., Figure 8 movie, arrow). Only one of these 11 interneurons was co-labeled for KCNQ3 (Figure 8—Figure supplement 3). In an equally sized sample of WT image stacks, none of 19 interneurons identified was somatically labeled for KCNQ2 and not KCNQ3. A small subset of interneuron AISs of both genotypes showed weak labeling of for KCNQ2 and KCNQ3.

We quantified KCNQ2 and KCNQ3 protein levels in neocortical tissue from P21 mice by western blot. As channel proteins localized to AISs and somatic domains could potentially track to different biochemical subcellular fractions, we performed preliminary experiments and developed sample preparation methods that allowed protein from whole tissue homogenates to be denatured and loaded onto SDS gels. We omitted centrifugal fractionation steps to avoid loss of detergent-resistant proteins in low-speed pellets, which might differ by genotype. Across conditions tested, western blots revealed the biochemical properties of KCNQ2 and KCNQ3 to be strikingly different, despite the high homology of the two protein sequences. E254*fs*/+ and G256W/+ samples showed an approximate 50% reduction in KCNQ2 monomer band versus WT, but no change in KCNQ3 (Figure 10). KCNQ2 blots showed additional bands corresponding to candidate dimer and higher oligomeric forms, but KCNQ3 blots showed a predominant monomer (Figure 10—Figure supplement 1). KCNQ2 monomer bands showed microheterogeneity, perhaps reflecting expression of multiple splice isoforms (Cooper et al., 2001; Pan et al., 2001). All KCNQ3 blots showed a single band running near the ∼97 kD mass calculated from its cDNA. To estimate total KCNQ2 protein, we quantified higher molecular weight KCNQ2 bands and the summed intensity of monomeric and higher molecular weight KCNQ2 bands. The sum of all KCNQ2 band intensities showed an approximate 50% loss, like the monomer. This loss was expected for E254*fs*/+ mice, based on prior qPCR and absence of a truncated protein band.. The loss was unexpected for G256W/+, though consistent with the hypothesis that mislocalized KCNQ2 subunits might have shortened half-lives in vivo. Yet, KCNQ3 protein levels in G256W/+ and WT mice were unchanged, despite the similar KCNQ2 and KCNQ3 redistribution from hippocampal pyramidal cell axons to somata exhibited immunohistochemically.

**Figure 10.**
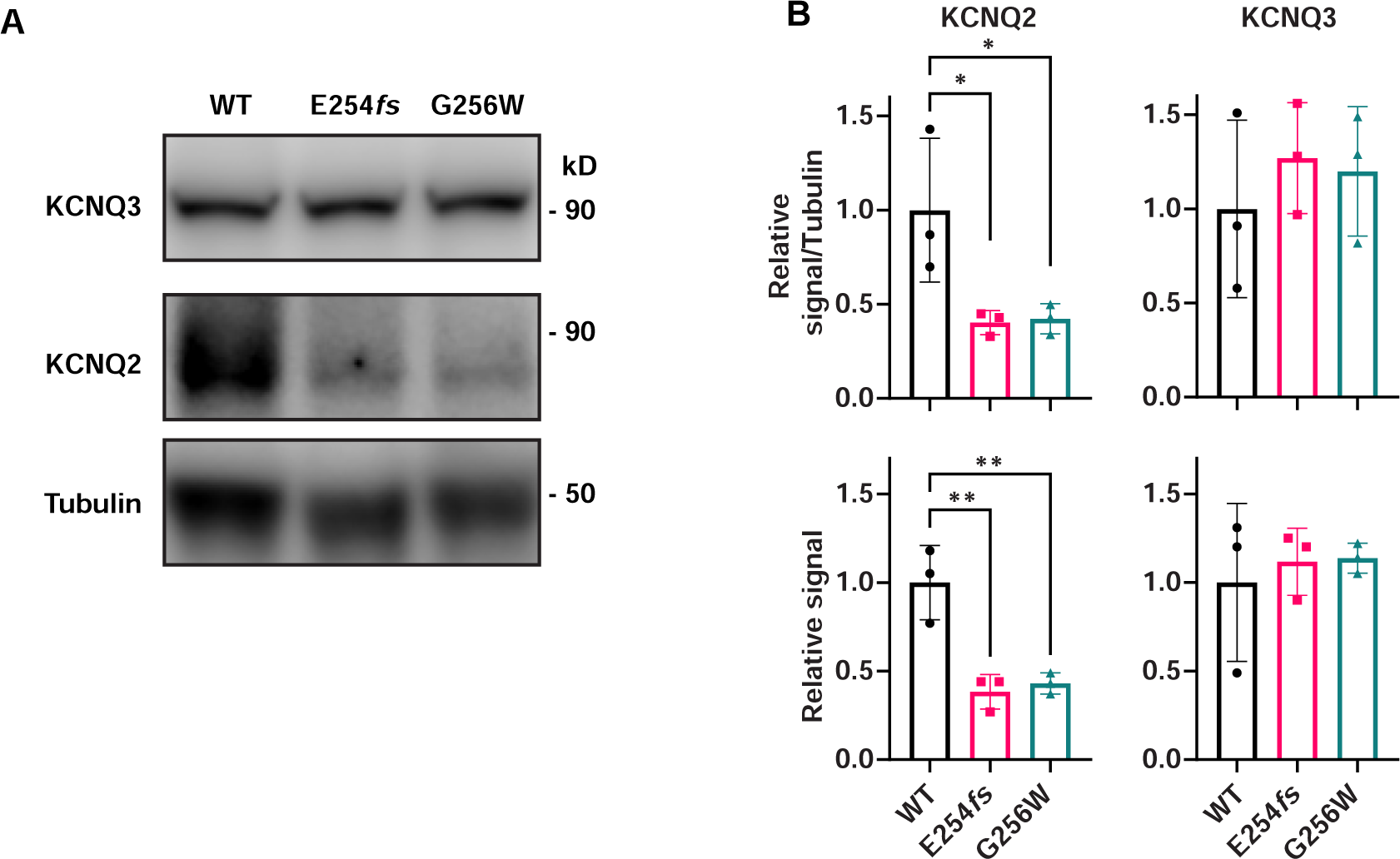
KCNQ2 protein is reduced in neocortex of P21 heterozygous E254*fs* and G256W mice. **A-B.** Representative western blots for all three genotypes probed for KCNQ2 and KCNQ3. **C.** Quantified KCNQ2 signal relative to WT, normalized to protein loaded as assayed by BCA. **D.** Quantified KCNQ3 signal relative to WT, n=3 per genotype, all males. One way ANOVA, * = P≤0.05, ** = P≤0.005.

## Discussion

Learning how an ion channel variant alters human development is challenging and of interest to diverse stakeholders. The studies summarized here bring together clinical, protein structural, in vitro, and in vivo approaches. By carefully considering individual 1’s medical history and then finding three additional affected individuals, we specified phenotype features to be modeled and generated independent clinical evidence of pathogenicity. Our lab studies highlighted three distinct mechanisms contributing to pathogenicity. First, we obtained evidence that Gly256 is part of an arch-shaped extracellular non-covalent bonding network between S5, the turret and the selectivity filter, a divergent and conserved structural feature of neuronal KCNQ2 and closely related subunits. This provided a novel hypothesis why a residue so distant from the pore might be pathogenic. Second, in heterologous cells, we found that expressing G256W with WT KCNQ2 and KCNQ3 subunits reduced currents consistent with the dominant-negative effects, though to an extent intermediate between previously described SLFNE and severe DEE variants. Third, we analyzed G256W/+ mice, learning that KCNQ2 and KCNQ3 proteins were quantitatively redistributed from hippocampal AIS and mossy fiber regions to somatic regions. KCNQ2 protein levels were reduced by about 50%, and CA1 neurons showed hyperexcitability. We performed experiments on E254*fs*/+ and G256W/+ mice in parallel. The contrasting results provide strong evidence that, in contrast to haploinsufficiency-linked SLFNE, in vivo dominant-negative effects of expressed G256W subunits drive DEE severity.

### Salient features of human G256W/+ related illness

Our clinical data, including de novo occurrence in three unrelated individuals and the informative phenotypic pattern observed in the European family including transmission from a mildly-affected mosaic parent (Weckhuysen *et al*., 2012; Myers *et al*., 2018), provide strong clinical evidence that KCNQ2 heterozygous G256W is pathogenic for DEE. Individual 1’s history includes features characteristic of KCNQ2 DEE as previously described but include some issues meriting comment. The patient’s epilepsy had an explosive early neonatal onset, with high seizure frequency and pharmacoresistance in the first 4 weeks of life. The potential that seizures began in utero was raised by the parents but was difficult to verify. Limited prior evidence suggests that, in some individuals born prematurely, onset of KCNQ2-related seizures is delayed until a post-conceptual age near term (Ronen *et al*., 1993). We note the issue, as in a previous report (Berg *et al*., 2021), to motivate broader, controlled survey. Although EEG records of individual 1 showed neither the invariant burst-suppression diagnostic of Ohtahara syndrome or hypsarrhythmia, ACTH was given and remission followed shortly after. Because seizure remission in infancy is characteristic of KCNQ2 DEE, the contribution made by ACTH is unclear and, among approved drugs, early trial of carbamazepine or oxcarbazepine is better supported (Pisano et al., 2015). Individual 1’s neonatal seizures exhibited a range of onset patterns, arising unilaterally from either hemisphere or bilaterally. This is more varied than previously described (Weckhuysen *et al*., 2013; Numis et al., 2014). Postictal voltage attenuation was a feature of nearly all (92%) of unilateral and bilateral onset seizures; this is unusual for neonatal seizures and should be studied in a large sample, as it may be useful diagnostically. Despite achieving seizure remission by 1 month, an age earlier than remission in many individuals with SLFNE (Ronen *et al*., 1993; Grinton et al., 2015), individual 1’s development showed global moderate to severe delay. This contrast between SLFNE and DEE outcomes despite impressive early seizure burden in both, seems to us to fit best within the emerging concept of “developmental encephalopathy” (Scheffer *et al*., 2017; Berg *et al*., 2021), i.e., it is the persistent, strong KCNQ2 loss-of-function after seizure remission that drives developmental impairment, rather than seizure-induced injury.

### Expressed in heterologous cells, KCNQ2 G256W shows intermediate severity dominant-negative effects and ezogabine-responsiveness

How can these clinical features be explained by experiments, and what questions remain? We performed whole-cell patch recordings using two methods. One team performed automated planar patch recordings, another made manual pipette-based patch recordings. The teams were blinded to each other’s work during data collection and analysis. Results showed good agreement, despite differences in sample preparation, cell selection, and recording protocols. Heteromeric KCNQ2/KCNQ3 channels showed no important differences between the methods. Because mean current density was larger using the manual patch method, likely due to selective study of high-expression cells in this method, the smaller KCNQ2 homomeric channel currents were studied by manual patch. Cells with 1:1 co-expression of WT KCNQ2 and G256W plasmids gave currents that were 30.1 +/- 0.1% of control. This is stronger conductance suppression than associated with KCNQ2 missense variants from SLFNE pedigrees (Schroeder et al., 1998; Coppola et al., 2003), but weaker suppression than exhibited by PGD de novo variants associated with very severe and profound impairment phenotypes (Orhan et al., 2014; Tran *et al*., 2020). Our manual-patch homomer recordings were insufficiently powered to achieve P < 0.05 for null hypothesis rejection between the ∼30% current observed and the 50% threshold for a “dominant negative” effect. Because cell-to-cell variability in current density is expected in heterologous transient expression systems, future studies aiming to categorize a spectrum of variant and phenotypic subgroups by this parameter should be designed with this goal in mind.

Our CHO cell transient expression results indicate that inclusion of one G256W subunit per tetramer is insufficient to fully prevent channel activity. Under conditions mimicking G256W heterozygosity, ezogabine shifted voltage-dependence and produced fold increases in currents to an equal extent as in cells expressing WT KCNQ2 only. By contrast, currents under conditions mimicking G256W homozygosity (G256W + WT KCNQ3 co-expression) showed no augmentation by ezogabine. Although the basis for these differences between conditions deserve more study, the most clinically relevant condition is mimicking heterozygosity, and such currents were strongly rescued by acute ezogabine treatment. Our working model (Figure 3A) used simplifying assumptions: that heteromeric currents preferentially reflect activity of channels with 2 KCNQ2 and 2 KCNQ3 subunits, and that (overexpressed from viral promoters in CHO cells) KCNQ2 WT and G256W subunits are equally represented. Future studies using surface biotinylation and tagged and/or concatemerized subunit constructs can extend these results by testing assembly and surface trafficking directly in CHO cells, cultured neurons, and mice. These experiments have therapeutic implications, as highlighted by cystic fibrosis, where effective multidrug therapies combine agents promoting surface expression and with others enhancing open probability (Middleton et al. 2019).

### The utility of mouse KCNQ2 DEE models, and some current limitations

We generated mice expressing G256W under the native mouse *Kcnq2* promoter. Homozygosity led to death at P0. G256W/+ mouse pups showed no differences in *Kcnq2* mRNA levels in hippocampus or cortex at age P21, gained weight and acquired a set of behavioral skills equally with WT controls, but experienced recurrent generalized seizures. We gleaned several lessons. In slices, CA1 pyramidal neurons exhibited normal resting membrane potentials but increased firing responses to current injections. These finding agree with prior studies of pyramidal cells of mice expressing other human pathogenic variants under the native *KCNQ2* promoter (Singh et al., 2008; Biba-Maazou et al., 2022). Both experimental data and modeling suggest that the combination of hyperexcitability and normal somatic biophysical properties may result from KCNQ2’s low somatodendritic and high AIS surface density (Shah et al.2008; Battefeld et al., 2014; Hu and Bean, 2018). The 25-35% increases in CA1 firing rates we observed are similar to prior studies of both SLFNE and DEE variants (Singh et al., 2008; Soh et al., 2014, Biba-Maazou et al., 2022), suggesting to us that the important severity differences between those variant classes involve contributions distant from the soma, from other neuronal types, and/or dynamic network interactions. Indeed, G256W/+ mouse ictal EEGs showed striking similarities with those from human patients, including individual 1 (Figure 1), namely evolution through a phase of high voltage repetitive spiking to a long period of strong voltage attenuation. A similar transition to EEG voltage attenuation was observed in studies of mice with conditional homozygous *Kcnq2* deletion from *Emx1*-expressing (primarily, glutamatergic) cells (Aiba and Noebels, 2021). Direct current EEG methods showed that ictal EEG attenuation in *Emx1-Kcnq2* null mice resulted from cortical spreading depolarization. Mice with deletion of Kv1.1/*Kcna1* had seizures without cortical spreading depression that could be converted to seizures with spreading depolarization by XE-991, the selective KCNQ family blocker. This highlights differences between the Kv1 and Kv7/KCNQ channel families discussed further below. It is expected, but important to learn, that spontaneous seizures of mice with heterozygous DEE variants driven by the native *Kcnq2* promoter will show spreading depolarization. G256W/+ mouse seizures were sometimes fatal (Figure 6, Figure 6—Figure supplement 1-Movie), as found previously in mouse lines with strongly suppressed KCNQ2 current made via diverse genetic strategies (Singh *et al*., 2008; Soh *et al*., 2014; Milh et al., 2020; Kim et al., 2021). Recent surveys show no instances of SUDEP among human KCNQ2 DEE cohorts, in contrast to Dravet syndrome, which has high SUDEP risk (Berg *et al*., 2021; Donnan et al., 2023). Our (JJ, SS, ECC) contacts with over 800 families linked to the KCNQ2 Cure Alliance and over 400 individuals enrolled in RIKEE have reveal no evidence of SUDEP. Study of *Kcnq2* mutant mice may illuminate human SUDEP mechanisms, nonetheless.

Parallel experiments performed on E254*fs*/+ and G256W/+ mice illuminated differences between SLFNE and DEE alleles that may contribute to differences in phenotype severity. In E254*fs*/+ mice, total *Kcnq2* mRNA was ∼30% reduced from WT, but RT-gPCR experiments using a WT transcript-selective assay showed 50% loss. This result indicated that nonsense mediated decay was incompletely efficient for E254*fs* messages. Incomplete nonsense-mediated decay has been described previously, and may show variation between genes, variants, tissues, and cell types (Zetoune et al., 2008; Dyle et al., 2020). Although most *KCNQ2* frameshift alleles have been associated with SLFNE via clinical studies of pedigrees (Millichap et al., 2016; Goto et al., 2019), the potential for some premature termination variants to escape nonsense-mediated decay and exert dominant-negative protein effects is highlighted by our results. This should be considered when stop-gained variants present with DEE phenotypes, especially if de novo and in the c-terminal region. Both E254*fs*/+ and G256W/+ mice showed increases in *Kcnq3* mRNA. In *Kcnq2* haploinsufficiency, increased relative numbers of KCNQ3 subunits may increase the proportion of KCNQ2 subunits forming KCNQ2/KCNQ3 heteromers. Such heteromers are activated at left-shifted voltages and have greater maximal open probability than KCNQ2 homomers. Such increased incorporation of KCNQ3 subunits could compensate for reduced KCNQ2 subunit availability. Increased KCNQ3 expression will do less when half the KCNQ2 subunits are dominant-negative DEE variants that impair in vivo surface trafficking and ion conduction. G256W/+ showed a loss of KCNQ2 but not KCNQ3 brain protein in western blots, even though the subunits were similarly redistributed from hippocampal pyramidal cell AISs to somata. This difference is unexplained and intriguing.

The most prominent shared phenotype of human *KCNQ2* (or *KCNQ3*) loss-of-function variants is frequent recurrent neonatal seizures, usually beginning from the first 2-3 days of life (Grinton *et al*., 2015; Millichap *et al*., 2016), as experienced by individual 1. Our study, in agreement with a recent study of *Kcnq2* T274M/+ knock-in mice (Milh *et al*., 2020), leads us to conclude that as-yet-unknown differences between species render mice much less susceptible to neonatal seizures. Seizures are highly penetrant in human SLFNE pedigrees, but no *Kcnq2* SLFNE mouse model known to date shows spontaneous seizures. G256W/+ and prior DEE model mice all show spontaneous generalized convulsive seizures and seizure-associated death (Milh *et al*., 2020; Kim *et al*., 2021). However, they do not exhibit the very explosive neonatal onset and transience found in children. This difference merits more investigation, including comparative interrogation of human postmortem brain mRNA and protein.

Impaired protein AIS localization resulting from KCNQ2 experimental and pathogenic missense variants in dissociated hippocampal cultures consisting predominantly of glutamatergic cells is long-known (Chung *et al*., 2006; Pan et al., 2006) but has not been previously observed under heterozygous conditions. KCNQ channels have been found in hippocampal parvalbumin and somatostatin interneurons though their roles remain very incompletely understood (Cooper *et al*., 2001; Lawrence et al., 2006; Fidzinski et al., 2015; Soh et al., 2018; Jing et al., 2022). Additional experiments using conditional *Kcnq2* alleles and interneuronal subtype-specific Cre’s, and dedicated studies of G256W/+ mouse interneurons are warranted. Interneurons of WT mice can exhibit somatic KCNQ2 immunolabeling in aldehyde-fixed sections (Cooper *et al*., 2001). Therefore, our detection of potentially increased labeling in G256W/+ interneuron somata (Figure 8 —Figure supplement 3) is best viewed as a motivation for more studies. Our efforts to quantitate changes in KCNQ2 protein levels by western blot again showed the remarkably different biochemical features of native-tissue KCNQ2 and KCNQ3 subunits, previously observed in mouse and human brain (Cooper et al., 2000; Kim et al., 2021). The two subunits have very similar properties when expressed heterologously in CHO or HEK cells (Tran et al., 2020). Mechanisms for the strong detergent-resistance and complex SDS gel migration pattern of brain KCNQ2 are poorly understood.

### Towards better variant pathogenicity prediction

The rapid discovery of novel variants of uncertain clinical significance in *KCNQ2* and other Kv channel genes is a challenge for pathogenicity assessment. *KCNQ2* has been included in efforts to develop high-throughput channel variant prediction methods integrating multiple data types via artificial intelligence (Bosselmann et al., 2022; Brunger *et al*., 2023). Brunger et al. analyzed variants in a very diverse set of channels, finding that pathogenicity was significantly predicted by variant distance from the geometric centers of the ion pore and membrane. KCNQ2 G256 lies about 20 and 30 Å from these locations, respectively, and was therefore of lower predicted risk, yet it is pathogenic. We noticed that KCNQ2 G256 and its turret neighbors were both evolutionarily divergent (from the *Shaker*-like Kv channels, and from closer relative, KCNQ1) and strongly conserved (among amniote KCNQ2s). G256 is part of a co-evolving set of residues and bonding partners, implying shared functional importance. Because specific, long-distance functional coupling mechanisms are well-established for voltage-gated channels, e.g., between residues mediating voltage-sensing, gating, and selectivity (Long et al., 2005; Hoshi and Armstrong, 2013; Zaydman et al., 2013), more detailed structural information including consideration of bond-pairing, gating associated conformational change, conservation, and divergence may improve automated pathogenicity prediction accuracy.

### Conclusion: a model linking molecular mechanisms to pathogenic consequences

Individual 1’s phenotype and variant location indicated that the pore turret could be important for KCNQ2 function in vivo. We tested this using structural modeling, heterologous cell voltage-clamp electrophysiology, and knock-in mice. Our results lead to a unifying model that is useful as it highlights a set of next experiments. The KCNQ2 selectivity filter could be stabilized by an extracellular hydrogen-bonded arch over the turret, and clinical heterozygous pathogenic variants could destabilize the arch and thereby, the pore. Prior studies of Kv channel slow and c-type inactivation highlight needed next experiments, including assessing the effects of [K^+^]_e_ on G256W current density, exploring the functional consequences of experimental mutations at other proposed turret network residues, and determining structures of mutant channels (Hoshi and Armstrong, 2013; Reddi et al., 2022; Tan et al., 2022; Ye et al., 2022; Fernandez-Marino et al., 2023; Wu et al., 2023). Ezogabine binds to a pocket on the pore domain between adjacent subunits, and its ability to correct structural consequence of PGD variants such as G256W can potentially be determined. The prominent EEG attenuation observed after KCNQ2 G256W human and mouse seizures, which is accompanied by spreading depolarization in *Emx1/Kcnq2* null mice (Aiba and Noebels, 2021), may be illuminating a distinctive KCNQ2 cellular and network function, and its basis in a canonical molecular property. Brown and Adams made use of a surprising, inverted voltage-clamp protocol with a holding potential of -30 mV, and measured currents elicited during and after brief steps to more polarized potentials (Brown and Adams, 1980). This protocol isolated I_M_ (primarily, KCNQ2/KCNQ3 current) because many other neuronal currents (notably including Kv1/*Sh* and Kv2 delayed rectifiers) have slow forms of voltage-dependent inactivation. Neurons that experience long-lasting depolarization after high activity or reversible injury must reboot through a temporally ordered process. Lack of voltage-dependent inactivation positions I_M_ to make early contributions to such neuronal recovery and repolarization. By contrast, the contribution of NaV and Kv currents with slow voltage-dependent inactivation will be delayed until after repolarization. The growing set of construct-valid mice including G256W and T274M provide platforms for testing this model in diverse circuits and paradigms of impaired development beyond seizures.

## Materials and Methods

### Human Subjects

US and European patients were enrolled after parental consent in the RIKEE registry, a human subjects research protocol approved by the Institutional Review Board of Baylor College of Medicine. Individual 1 (USA) video-EEG review was performed by a board-certified clinical neurophysiologist on archived recordings made as part of routine care. EEGs began after transfer from the birth hospital to a tertiary care center on the fifth day of life. The archived records included all seizures detected on initial clinical review of the continuous VEEG; interictal background was sampled in saved segments taken about every two hours. Individual 1 received ACTH, 60 units/m^2^, twice daily for two weeks followed by a 4 week taper (Takacs, 2023). Individual 1 underwent clinical Sanger sequencing (GeneDx). Individual 2 was diagnosed using an in-house multigene panel and informatics workflow (case 14; Jang et al., 2019). Individuals 3 and 4 were diagnosed by a 32 gene NGS panel (Familial and Generalized Epilepsy Gene Panel, Center for Human Genetics Tübingen; CHGT). Mosaicism was diagnosed for individual 4 at CHGT by lab-established procedures taking into account deviation from expected 50% read counts and the clinical history.

### Structural modeling Interpretation

We examined structural models of KCNQ1, KCNQ2, KCNQ4 and KcsA using Mol*, UCSF Chimera and ChimeraX (Goddard et al., 2018). We visualized turret main chain bond angles in Pymol. We made pairwise turret alignments in Chimera and used Mol* to identify predicted noncovalent bonds in the turret region using default cut-off parameters (for hydrogen bonding: length 3.5, maximum angle deviation 45). We visualized model B-factor and surface electrostatic potential maps in ChimeraX, and assessed the local resolution and model congruence of the turret region of KCNQ2 in aligned electron density maps (EMD-30443, EMD-30446) using Chimera.

### Mouse Video-EEG monitoring

Using methods previously described in detail, electrode implantation surgeries and EEG recordings were performed in-lab and at the Baylor College of Medicine IDDRC In Vivo Neurophysiology Core (Creson et al., 2019; Jing *et al*., 2022). A post-operative recovery period of 2 days was allowed before commencing video-EEG monitoring. During recordings, mice were able to explore their cages freely and had access to water and food. Seizures counts were made by video and EEG review under band pass filtering (1-59 Hz). Seizure events were clipped and saved.

### Generation and use of KCNQ2 heterozygous G256W and E254*fs* mice

#### Crispr and mouse husbandry

All animal procedures were performed in accordance with protocols reviewed and approved by the Institutional Animal Use and Care Committees of the Jackson Laboratories, Baylor College of Medicine, and the University of Connecticut. KCNQ2 G256 is conserved in all amniotes (Figure 2C), but the codons are not (C57B6/J mouse: GGU, human: GGG). Therefore, we made two base substitutions to introduce W256 (Figure 7—Figure supplement 1, panel D). C57BL/6 single cell zygotes were microinjected with a 123-nt oligonucleotide donor sequence:

5’-tggtacattggcttcctctgcctcatcctggcctcatttctggtgtacttggcagaaaagTgGgagaatgaccactttgacacctacgcagatgcactctggtggggtctggtaagtcctggt-3’ containing two nucleotide differences (capitalized) to change the glycine GGT codon to a tryptophan TGG codon (G256W). Co-injected with the donor oligonucleotide was the *Kcnq2* exon 5 targeting guide RNA; 5’-TCTGGTGTACTTGGCAGAAA-3’ and Cas9. Incorporation of the G256W mutation into the genome would change the AGG PAM recognition sequence to AGT and prevent retargeting of the modified allele. Founders were generated after embryo transfer into pseudopregnant C56BL/6J females and screened by sequence analysis of the *Kcnq2* exon 5 genomic DNA using PCR and sequencing primers 5’-GGGATTCCATCCTCCAAGTC-3’ and 5’-CCAGCCCAGCCTAAAGACA-3’. A single founder female was identified from 40 progeny that carried the desired G256W mutation in trans to a frameshifting indel E254*fs*\*16 mutation (deletion of the AAAGGG nucleotides overlapping with the PAM sequence). Lines were established from both alleles after three successive backcrosses to C57BL/6J mice and designated as Jax Stock numbers 029407 and 029408, respectively. Genotype by sequencing protocols were later replaced with RT-PCR fluorometric probe assays, specific for the WT, G256W and E254*fs*\*16 deletion mutations (Transnetyx).

#### Survival analysis

Only animals that were backcrossed five or more times were included in the survival analysis. During the COVID pandemic, many animals were euthanized in response to mandated colony reduction and research suspension; these animals were also excluded from analysis. Animals were censored if they were euthanized due to other (non-seizure related) health reasons or to not exceed IACUC approved animal usage numbers. All early deaths in G256W mutants appeared to potentially be the result of fatal seizures, as all recovered carcasses exhibited a stereotyped posture with symmetrically flexed forelimbs and extended hindlimbs.

#### Developmental milestone assays and behavioral seizure recording

Developmental milestone assays were performed following previously described methods (Hill et al., 2008). Pups were bred group-timed matings from first time breeders to avoid single litter effects. To test surface righting, an investigator held an individual mouse pups gently on its back, then measured the time required for the pup to flip onto its abdomen after being released. For the negative geotaxis test, an investigator placed the mouse on a 45° sloped surface with its head pointing downhill, then measured the time required for the pup to turn its body 180°. To test cliff aversion, an investigator positioned the mouse pup’s forepaws on the edge of a smooth surface 4” above a table, then recorded the time for the pup to turn its body and take a step away from the edge. The P10 seizure (Figure 4) was observed during a pilot study of pup open field motor behavior conducted as previously described (Bass et al 2020).

#### RT-qPCR

All P100 experiments included the same 3 males and 3 female mice per genotype. For P21, 3 males and 3 females were collected for WT and E254*fs*/+ mice, while 2 males and 3 females were collected for G256W/+ mice. Animals were deeply anesthetized with isoflurane and decapitated. Brains were removed and neocortical and hippocampal regions dissected on ice. Samples were frozen on dry ice and stored at -80°C. RNA was isolated from brain tissue (RNeasy Lipid Tissue kit, Qiagen) and stored at -80°C until use. RNA (1 μg/sample) was used to generate cDNA (SuperScript III Reverse Transcriptase, Invitrogen), which served as template for qPCR using ThermoFisher Scientific TaqMan gene expression assays with the TaqMan Fast Advanced Master Mix. Assays were run on the QuantStudio 3 PCR system (Applied Biosystems), using *Gapdh* as reference and the ΔΔCt quantification method (Livak and Schmittgen, 2001). For measuring total *Kcnq2* and *Kcnq3* mRNA, the PCR amplicon spanned exons 5 and 6 (*Kcnq2* assay) or exons 4 and 5 (*Kcnq3);* primers and probes recognized sequences common to all known splice isoforms (Pan et al., 2001). For the qPCR assay discriminating between *Kcnq2* WT and variant alleles, we developed a custom TaqMan assay (ThermoFisherScientific) where the fluorescent probe binding site included all 8 WT bases deleted or substituted in the mutant alleles. For Figure 7, each (n = 303) individual data point is the mean of three replicate wells. One additional result was determined post hoc to be an outlier by the Grubbs’ test and is not shown (P21 male neocortex, *Kcnq2* G256W/+, allele non-specific probe, Figure 7A; thus n = 1 male, and n = 3 females). Each of the 16 graphs summarizes experiments performed in parallel on a PCR plate. ThermoFisher Scientific TaqMan Assay ID’s are listed in the key Resources table.

#### cDNA Sanger sequencing

cDNA made from 1000 µg of total RNA was amplified using primers spanning *Kcnq2* exons 4 through 7 (Apex Hot Start 2X Master Mix; Blue Apex Buffer 1, ID No. 5200600-1250). cDNA was amplified using the following touchdown protocol: 1. 95°C 15 min 2. 94°C 20 sec 3. 64°C 30 sec (-1°C every cycle) 4. 72°C 30 sec 5. Go to step 2 (repeat 5 times) 6. 94°C 20 sec 7. 59°C 30 sec 8. 72°C 30 sec 9. Go to step 6 (repeat 34 times) 10. 72°C 10 min 11. 4°C hold. The PCR reaction product was cleaned (Zymo DNA Clean & Concentrator Cat. No D4013) and sequenced (Genewiz) using the same primers used for amplification.

### Heterologous expression and CHO cell recording

#### Automated patch

Methods for expression and recording in CHO cells were previously described in detail (Vanoye *et al*., 2022). Human KCNQ2 cDNA (GenBank accession number NM_172108) in pIRES2-EGFP (BD Biosciences-Clontech, Mountain View, CA, USA) was used as template for in vitro mutagenesis. A stable line expressing human KCNQ3 (GenBank accession number NM_004519) was generated as described and maintained under dual selection with Zeocin (100 μg/ml) and hygromycin B (600 μg/ml). Plasmids with WT KCNQ2 or G256W cDNA were introduced into the KCNQ3-expressing CHO cell stable line by electroporation (Maxcyte STX; MaxCyte Inc., Gaithersburg, MD, USA). Automated patch clamp recording was performed using the Syncropatch 768 PE platform using PatchController384 V.1.3.0 software (Nanion Technologies, Munich, Germany). Pulse protocols were performed before and after addition of ezogabine (10 μM, Sigma-Aldrich) and, subsequently, XE-991 (25 μM, Abcam, Cambridge, MA; or TOCRIS, Minneapolis, MN). Currents reported are XE-991-sensitive currents, calculated by subtraction.

#### Manual patch

Mutagenesis, cell culture, transfection, and manual patch recordings were performed as previously described (Tran *et al*., 2020). CHO cells plated on cover slips were recorded at room temperature (20–22°C), 2–3 days post-transfection, using an Axopatch 200B amplifier (Molecular Devices), pCLAMP v.9, a cFlow perfusion controller and mPre8 manifold (Cell MicroControls), and glass micropipettes (VWR International) with 1–4 MΩ resistance. The extracellular solution consisted of (in mM): 138 NaCl, 5.4 KCl, 2 CaC_l2_, 1 MgCl_2_, 10 glucose, 10 HEPES, pH 7.4 with NaOH (Miceli et al., 2013). Pipette solution contained (in mM): 140 KCl, 2 MgCl_2_, 10 EGTA, 10 HEPES, and 5 Mg-ATP, pH 7.4 with KOH. Series resistance was compensated by 70% after compensation using Axopatch 200B fast and slow capacitance controls. Currents were digitally sampled at 10 kHz and filtered at 5 kHz using a low-pass Bessel filter. For voltage-activation experiments, cells were held at −80 mV and depolarized in 10 mV incremental steps from −80 to +40 mV for 1 s, then stepped to 0 mV for 60 ms, followed by a 20 sec, -80 mV interpulse. Tail currents were fitted using the Boltzmann function in Prism to obtain the half-maximum activation voltage (*V*1/2) and the slope factor (*k*).

### Whole Cell CA1 pyramidal cell recording

#### Acute brain slice preparation

For all electrophysiological experiments, we used P12–P15 mice of both sexes. The mice were anesthetized with isoflurane and quickly euthanized through decapitation. Subsequently, their brains were extracted and placed in a chilled cutting solution composed of 26 mM NaHCO_3_, 1.25 mM NaH_2_PO_4_, 2.5 mM KCl, 0.5 mM CaCl_2_, 7 mM MgCl_2_, 10 mM dextrose, and 210 mM sucrose. To record from the hippocampus, 300 μm slices were cut horizontally using a microtome (Leica VT1200S). These slices were then moved to a holding chamber containing artificial cerebrospinal fluid (ACSF), which contained 125 mM NaCl, 2.5 mM KCl, 1.3 mM MgCl_2_, 1 mM NaH_2_PO_4_, 26 mM NaHCO_3_, and 12 mM dextrose, 1.5 mM CaCl_2_ was supplemented the day of the recording. Both the cutting solution and ACSF were consistently saturated with a mixture of 95% O_2_ and 5% CO_2_. The brain slices in ACSF underwent a 30-min incubation in a 37°C water bath, followed by at least one hour at room temperature prior to the actual recordings. Subsequently, the slices were transferred to a recording chamber where the temperature was maintained at 30–32°C using a Warner Instruments TC 324C temperature controller. Continuous perfusion of ACSF into the chamber was achieved through a peristaltic pump.

#### Electrophysiological recordings

We used borosilicate glass electrodes with resistances of 2–4 MΩ for conducting whole-cell recordings. Current clamp recordings were done on ventral CA1 pyramidal neurons of the hippocampus. Recording pipettes were filled with an internal solution composed of 130 mM CH_3_KO_4_S, 10 mM KCl, 4 mM NaCl, 4 mM Tris-phosphocreatine, 10 mM HEPES, 4 mM Mg-ATP, and 0.4 mM Na-ATP. These neuronal recordings took place in the presence of synaptic blockers, including 100 μM picrotoxin to inhibit GABAA receptor-mediated inhibitory responses, 4 μM NBQX to block AMPA-mediated responses, and 10 μM D-AP5 to inhibit NMDA-mediated responses. All current clamp recordings were performed using a Multiclamp 700B amplifier (Molecular Devices) with bridge balancing engaged to compensate for input resistance. Cells with an access resistance of less than 20 MΩ were selected for both recording and subsequent analysis. To evaluate intrinsic excitability and action potential waveform characteristics, a depolarizing current injection ranging from +25 pA to +325 pA was applied in +25 pA increments, each lasting for 1 s with 15 s intervals between sweeps. To determine the cell’s input resistance, a series of hyperpolarizing steps spanning from -100 pA to 0 pA were administered in -25 pA increments, with each hyperpolarizing step maintained for 1 s and no intervals between sweeps. The holding membrane potential was set at -65 mV prior to the step protocols by injecting small DC current through the pipette. Resting membrane potential was measured at I=0 soon after breaking into the cell. AP properties were determined by the 1^st^ action potential in a step protocol. Data were sampled at 50 kHz, with the Bessel filter set at 10 kHz. Data acquisition for all electrophysiology experiments was executed using a Digidata 1440 A system and pClamp software (versions 10.2–11.2; RRID:SCR_011323).

#### Immunohistochemistry

We trialed three alternative tissue preparation protocols: unfixed cryosections with or without ice-cold methanol pre-staining steps (Devaux *et al*., 2004, Pan et al., 2006), and a previously used protocol using weak formaldehyde perfusion fixation followed by microwave/citrate antigen retrieval (Shah *et al*., 2008). The formaldehyde/antigen retrieval method more reliably yielded intact tissue on slides, an outcome that was important for the analysis we planned comparing multiple matched sections per animal and multiple animals per each of 3 genotypes. Accordingly, we adopted that approach for experiments used for quantitation shown herein. After establishment of deep anesthesia (300 mg/kg ketamine/30 mg/kg xylazine IP), mice were transcardially perfused with 20 mL of sterile ice-cold PBS, followed by 20 mL of ice cold 2% formaldehyde in PBS, freshly prepared from paraformaldehyde 20% aqueous solution (Electron Microscopy Sciences). The brain was removed from the skull, post-fixed on ice for 60 min, embedded (OTC Tissue Tek Compound, Sakura Finetek), and stored at 80°C. Coronal 40 μm sections including dorsal hippocampus were cut on a cryostat, transferred to slides (Fisher SuperFrost Plus, Fisher Scientific), and stored at -80°C. Slides bearing brain sections at matched rostrocaudal positions of the dorsal hippocampus were thawed for 10 min at room temperature (RT), subjected to citrate/microwave antigen retrieval, washed with PBS for 10 min, and blocked (PBS, 5% normal goat serum, 0.5% Triton-X 100). Sections were incubated overnight with blocking buffer containing primary antibodies (affinity-purified rabbit anti-KCNQ2 N-terminal antibody, 1:200, KCNQ3 N-terminal antibody, 1:500, and either mouse anti-AnkG IgG2a, clone N106/36, Neuromab/Antibodies, Inc., 75-146, 1:1000, or mouse anti-PanNav IgG1 (hybridoma culture supernatant, 1:200, gift of Jim Trimmer, commercially available as affinity-purified IgG, Sigma K58/35 S8809) using incubation chambers formed with CoverWell Gaskets (ThermoFisher C18150) to prevent drying. Wash steps used PBS with 0.05% Triton-X100. Slides were coverslipped using ProLong Gold with DAPI (ThermoFisher P36931).

#### Quantification of KCNQ2 and KCNQ3 immunolabeling intensity

Confocal imaging was performed on trios of slides differing in genotype (WT, G256W/+ and E254*fs/+*) and immunostained in parallel for KCNQ2. KCNQ3, and PanNaV. The slides were labeled only by numerical codes--the experimenter performing the imaging and quantification analysis was blinded to genotype throughout. Sample trios were imaged under the exact same settings. Image stacks for analysis were collected in CA1B, CA3A, and the dentate. Prior to final image collection, nearby optical fields were used to adjust acquisition parameters to optimize dynamic range and eliminate detector saturation for any sample within the set. Image stacks (pixel XY dimensions: 0.31 μm,1024x1024 pixels/image, 21 images/stack, z-step of 0.500 μm) were generated using a Nikon C2 confocal microscope (20X 0.75 NA Nikon planapo lens, and 2X zoom), running NIS Elements 4.0 AR. Quantitation was performed on maximal projections of the confocal stacks. Regions of interest (ROIs) were marked by DAPI (for somata), and PanNaV (for AISs and mossy fibers in the dentate hilus and CA3). In CA1 and CA3, the somatic ROIs were portions of stratum pyramidale without AISs. In the dentate, the fine AISs could not be excluded as they lie within the granule cell layer (Shah et al., 2008), but compared to CA1, they are very thin and inconspicuous under 20x magnification. KCNQ2 and KCNQ3 color channels were masked during manual marking of somatic and AIS or mossy fiber ROIs. The ROI-setting procedure used with two steps. First, the blinded investigator used the freehand drawing tool, and PanNaV to mark the AIS-and mossy fiber containing ROIs, and used DAPI labeling delimit the remaining portions of the pyramidal cell or granule cell layers excluding zones of overlap. Second, within these regions, 5 smaller, equally sized, rectangular ROIs were placed, avoiding any visible AISs in the soma-containing ROIs. The intensity values for KCNQ2 or KCNQ3 were exported and averaged across all ROIs for each sample. AIS or MF mean ROI intensities were then divided by those of adjoining somata regions (e.g., the average ROI intensity for KCNQ2 or KCNQ3 in the AIS region of CA1 divided by the average intensity in the CA1 pyramidal cell layer from the same image stack). This analysis was performed on two image stacks per animal per region (one per hemisphere), and 3 animals per genotype. Male and female mice were both included since completed qPCR and preliminary immunohistochemical experiments showed no sex differences. Putative CA1 interneurons were identified by size, location, somatic labeling for AnkG and/or KCNQ2, and presence of a neighboring AIS.

### Immunoblotting

For experiments shown in Figure 10 and Figure 10—Figure supplement 1, tissues from individual animals were processed separately, in parallel, to allow for comparison of biological replicates of each genotype. Animals were deeply anesthetized with isoflurane and decapitated. Brains were removed and neocortical and hippocampal regions dissected on ice. Samples were frozen on dry ice and stored at -80°C. Cortex samples were weighed and homogenized with a glass-glass homogenizer in 10 volumes (w/v) of RIPA (150 mM NaCl, 1.0% Triton X-100, 0.5% sodium deoxycholate, 0.1% SDS, 50 mM Tris HCl pH 7.4, Pierce EDTA-free protease inhibitor). Homogenates were sonicated (Branson model 450, Cat. No. 15338553) with a 1/8” microtip (Cat. No. 101063212) at 40% amplitude for 10 sec three times, aliquoted and stored at -80°C. Protein concentration was determined by bicinchoninic acid assay (BCA, Pierce). Homogenate aliquots were thawed on ice, supplemented with SDS sample buffer (Licor) and dithiothreitol (10 mM final concentration), incubated at 37°C for 30 min, mixed, and loaded on 7.5% or 4-15% Mini-Protean TGX gels (Biorad). Resolved proteins were electro-transferred to PVDF. We incubated filters in 5% non-fat dry milk (Carnation) in Tris-buffered saline with 0.2% Tween (TBST) for one hour, then overnight in the same buffer with affinity-purified anti-KCNQ2 or anti-KCNQ3 primary antibodies at 1:400 and 1:1000, respectively. After washing, filters were incubated with HRP conjugated secondary antibodies. Blots were visualized by enhanced chemiluminescence (Amersham Cytiva ECL Prime), and imaged using a CCD-based gel documentation system (LiCor Odyssey XF). Although blots were probed in parallel for tubulin as a loading control (Sigma T6199) protein loading required for channel subunit detection was outside the tubulin linear range (Kirshner and Gibbs, 2018).Quantification was performed using normalization to tubulin and based on equal sample loading (i.e., to the BCA) with similar results. Quantification of band intensities was performed using Image Studio (LiCor, v.5.2). Background was subtracted by the Image Studio software’s individual band 4-sided surround method.

### Statistical analysis

#### Development, qPCR, Immunohistochemistry, and western blot

Statistical tests were applied using Prism version 9.5.1 (GraphPad Software, San Diego, California USA). Data distribution normality was analyzed using the D’Agostino-Pearson test. For comparison of observed E254*fs*/+ *Kcnq2* mRNA levels versus the expected 50%, a one-sample t-test was conducted. For other RT-qPCR, western blot, and immunohistochemistry (AIS and mossy fiber vs somatic intensity) comparisons, group differences were tested by two-way ANOVA to determine if genotype, sex, and/or the interaction between the two had a significant effect. As sex was found not to significantly affect the data, it was dropped as a factor and a one-way ANOVA was performed with genotype as the only factor. A paired t-test was used to compare *Kcnq2* mRNA levels in E254fs/+ mice between allele non-specific and wildtype specific probes. For mouse developmental milestone data, the same approach was used except for using a repeated measures ANOVA. Tukey’s HSD post hoc test was used to assess pairwise significance between genotypes.

#### Electrophysiology

Statistical tests were applied using Prism version 9.5.1 (GraphPad Software, San Diego, California USA). Data distribution normality was analyzed using the D’Agostino-Pearson test. For in vitro patch clamp recordings, I/V and G/V curves were analyzed using a two-way repeated measures ANOVA with the Geissner-Greenhouse correction, matched values from each recorded cell were stacked into a subcolumn. Test potential (voltage) and expression (e.g., Q2 WT + G256W) were defined as factors. Pairwise comparisons were made between expression groups at each test potential, with Tukey’s post hoc analysis correcting for multiple comparisons. For comparing heterozygous homomeric currents to 50% of WT, a t-test was performed between the average current densities at +40 mV. For slice recording data, the same approach was used but with action potential number and current injection being defined as factors.

## Additional Information

## Supporting information

Fig.1-S1 EEG

## Acknowledgements

This work was supported by an American Epilepsy Society Predoctoral Fellowship funded in part by Wishes for Elliot (TJA); by Citizens United for Epilepsy Research, the Jack Pribaz Foundation, the KCNQ2 Cure Alliance, the Miles Family Fund, and R01 NS49119 (ECC), by the NINDS Epilepsy Center without Walls U54 NS108874 (CV, ALG, NV, AT, KS, ECC); by NIH Grants R01 NS101596 and NS108874 (to AVT); by NS096029 and MH126953 (to AM); by NINDS U54 OD020351 (AZ, CL); by R01 NS29709 (JLN); and by K08NS110924 (VK). The Jackson Laboratory scientific services are supported in part through the National Cancer Institute’s Cancer Core Grant P30CA034196. We acknowledge contributions from The Jackson Laboratory Genome Engineering Team and the Transgenic Genotyping Services for assistance and consultation during the generation of the *Kcnq2* mutant mice. The BCM Mass Spectrometry Proteomics Core was supported by the Dan L. Duncan Comprehensive Cancer Center NIH Award P30 CA125123, CPRIT Core Facility Award RP210227, Intellectual Development Disabilities Research Center P50 HD103555, and an NIH High End Instrument Award S10 OD026804. EEG at the Baylor College of Medicine Intellectual and Developmental Disabilities Research Center Neuroconnectivity Core was supported by NICHD P50HD103555. The BCM CryoEM Core and ZW were supported by NIH R01GM143380 and R01HL162842, the Welch Foundation Q-2173-20230405, and by a CPRIT Core Facility Award RP190602.

## Web resources

RIKEE KCNQ variant database, www.rikee.org

PDB, www.rcsb.org

EMDB, www.ebi.ac.uk/emdb

---

**Figure 1—figure supplement 1.**
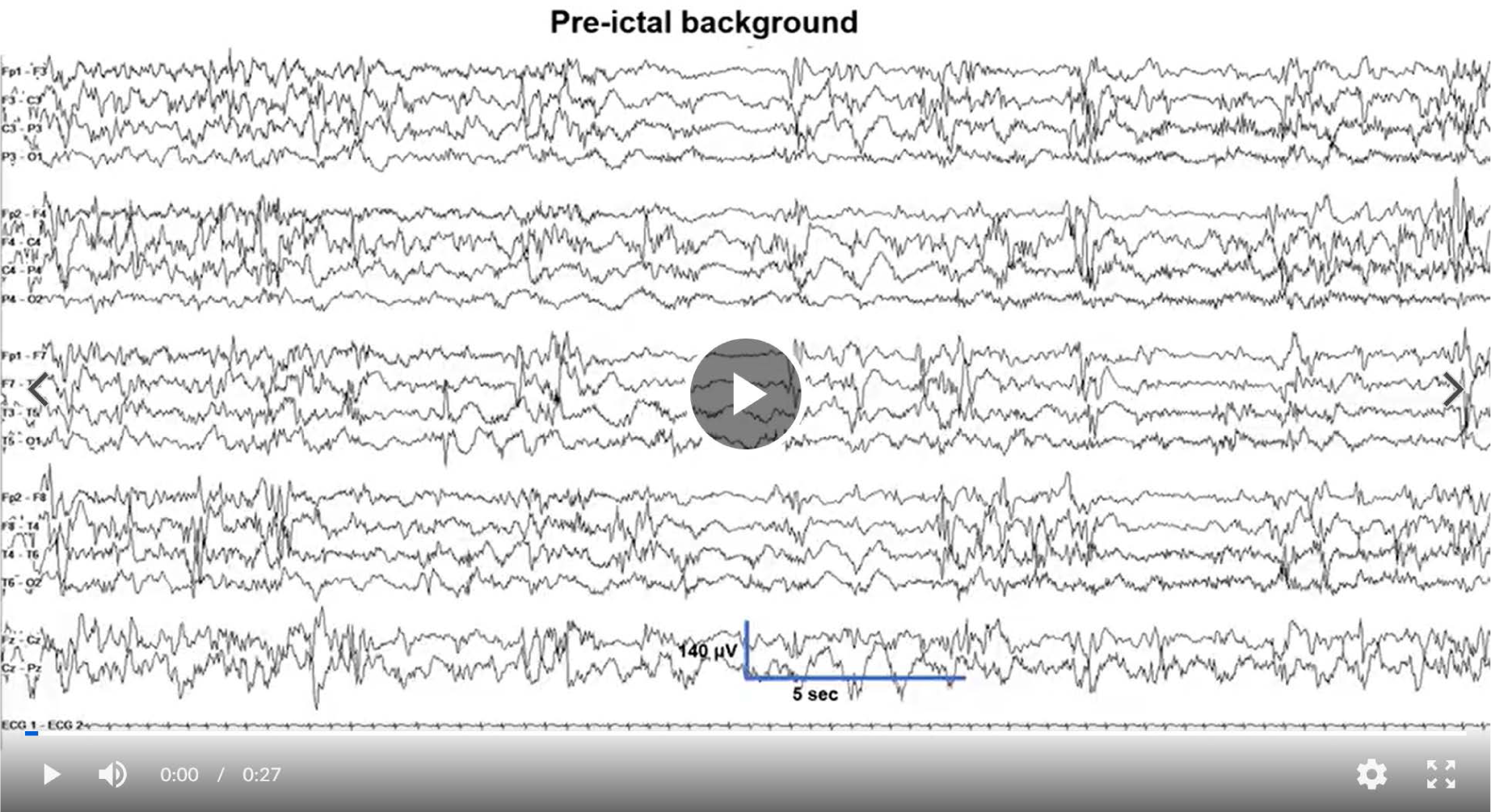
EEG recording including pre-ictal, post-ictal attenuation and recovery of background of seizure excerpted in Figure 1. As labeled, onset was preceded by eyeblink and muscle artifact. The interval of uninterrupted voltage attenuation between the end of the high voltage fast activity to the first epileptiform burst was 61 sec. Interburst length progressively shortened in length, over about 3 min. Settings as in Figure 1. <u>Link to movie F1-S1.</u>

**Figure 1—figure supplement 2.**
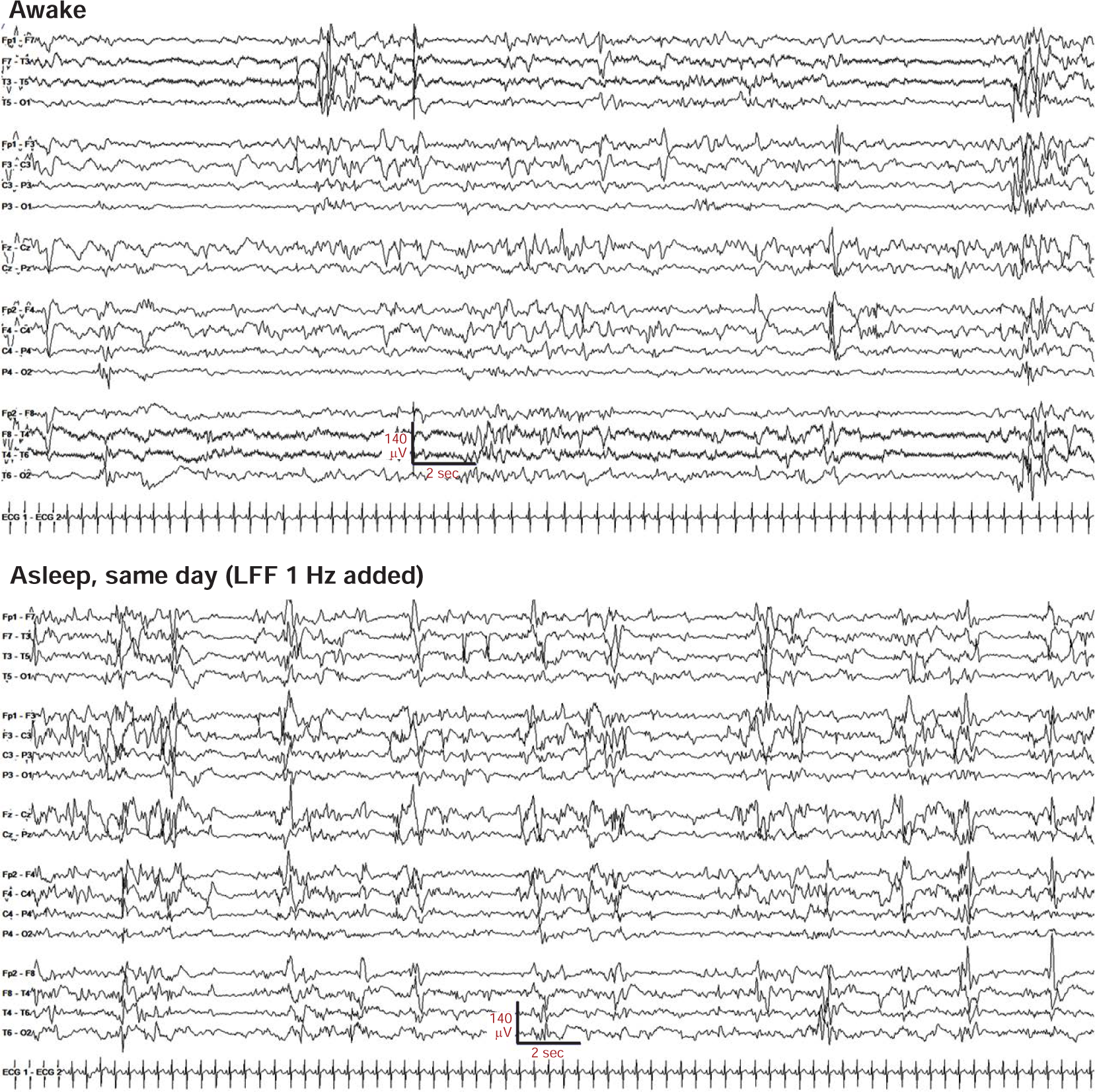
Examples of awake and sleep EEG background. There is evidence of state change, with more discontinuity during sleep. The awake excerpt shows variable frequency composition, and excess multifocal sharps. Excess discontinuity and sharps indicate dysmaturity, but burst-suppression is not seen. Settings as in Figure 1, except LFF changed to 1 Hz.

**Figure 2—figure supplement 1.**
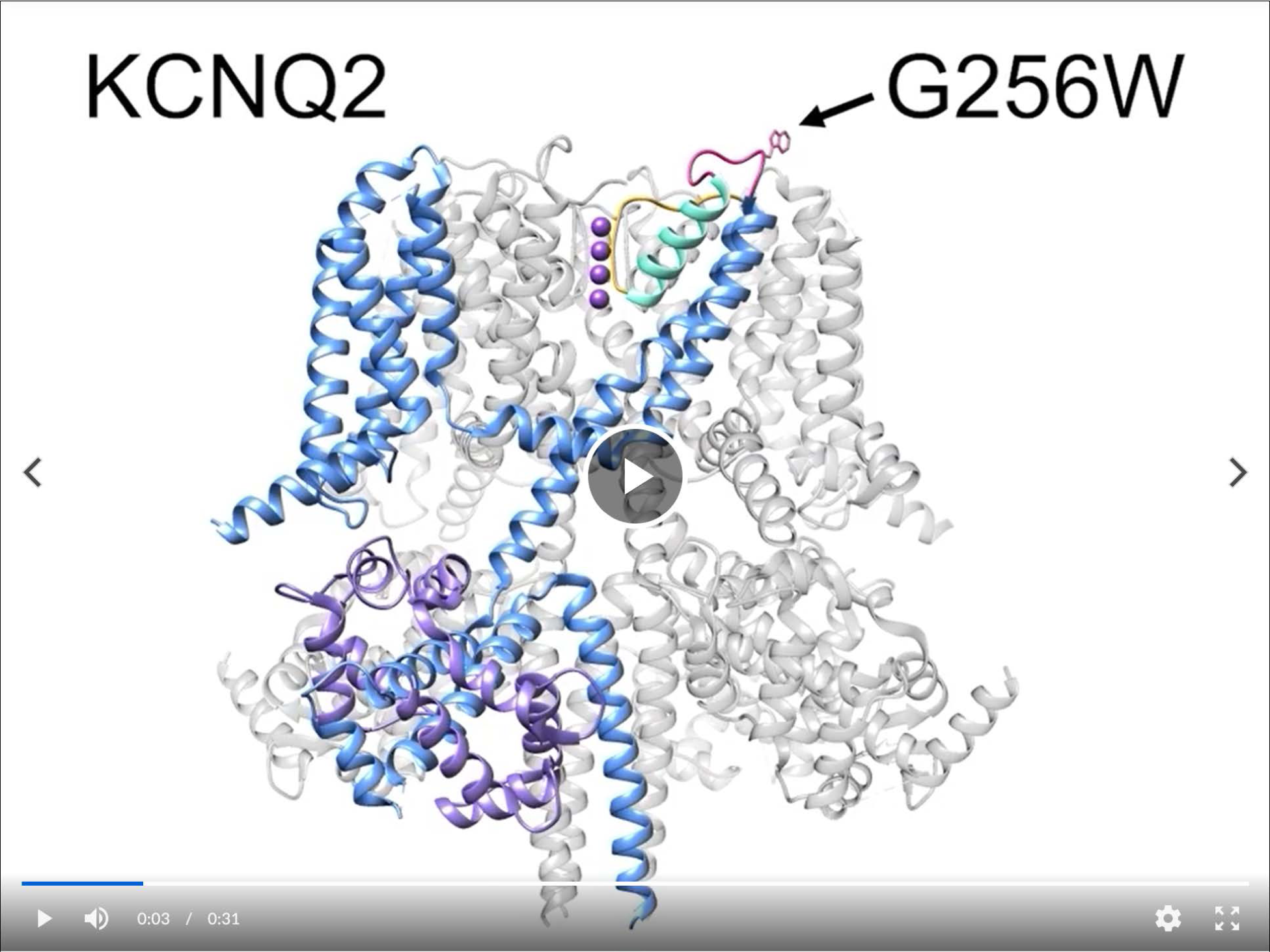
Movie illustrating position of the G256W substitution within the KCNQ2 channel pore turret and its distance to the selectivity filter. <u>Link to movie F2-S1</u>.

**Figure 2—figure supplement 2.**
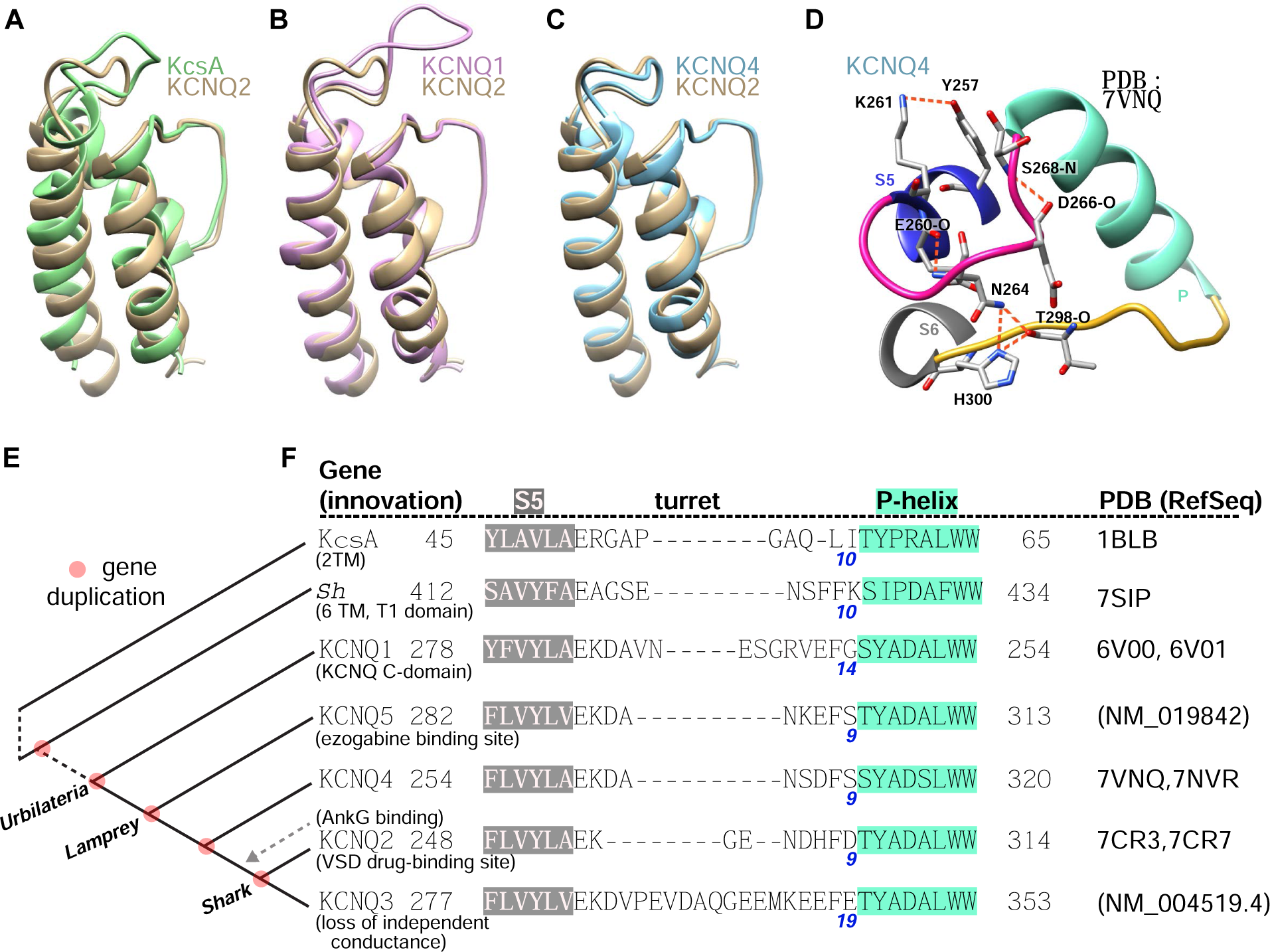
The G256W variant affects a divergent neuronal KCNQ turret structure enabling forming a bonding network linked to the ion selective pore. **A-C.** Aligned structural models of the extracellular portions of PGDs of KCNQ1, KCNQ4 and KcsA with that of KCNQ2. Single subunits are shown. **D.** Cartoon of structural model of turret region of KCNQ4 highlighting the predicted hydrogen bonding network. Several bonds are conserved between KCNQ2 and KCNQ4, but the KCNQ4 network has fewer bonds (compare with Figure 2G). **E.** Cladogram summarizing evolutionary relationships among several voltage-gated potassium channel genes. Gene duplication(s) are indicated by red circles, and are labeled by a common ancestor (or their extant descendant) possessing both duplicate genes. **F.** Sequence alignments of KCNQ1-5 P loops and flanking S5 and S6 regions reveal relative conservation of KCNQ5, KCNQ4, and KCNQ2 (as in Figure 2C), and divergence of KCNQ1 and KCNQ3.

**Figure 2—figure supplement 3.**
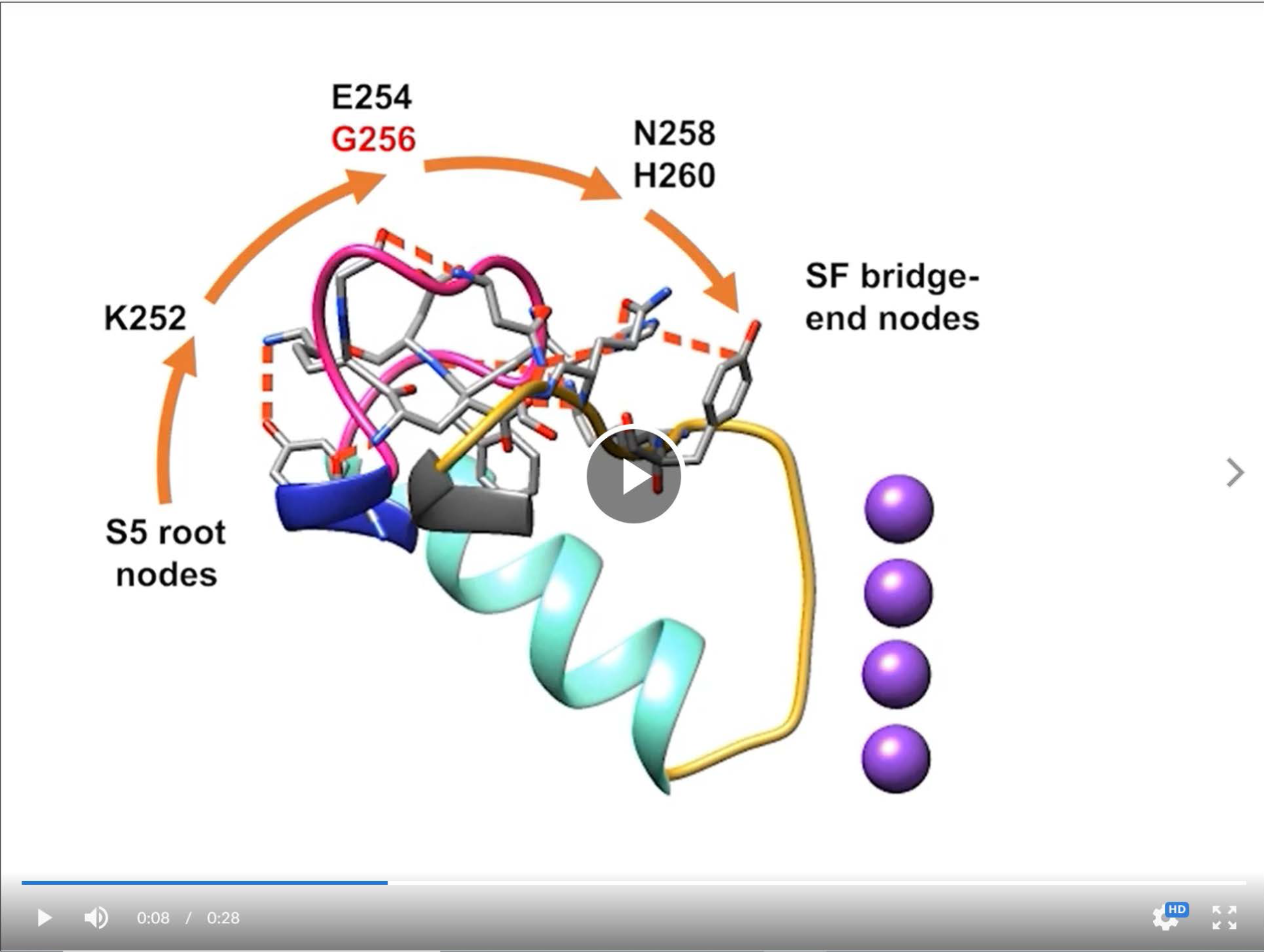
Movie illustrating locations of residues contributing to a non-covalent bonding network extending from S5 to the selectivity filter. <u>Link to movie F2-S3</u>.

**Figure 3—figure supplement 1.**
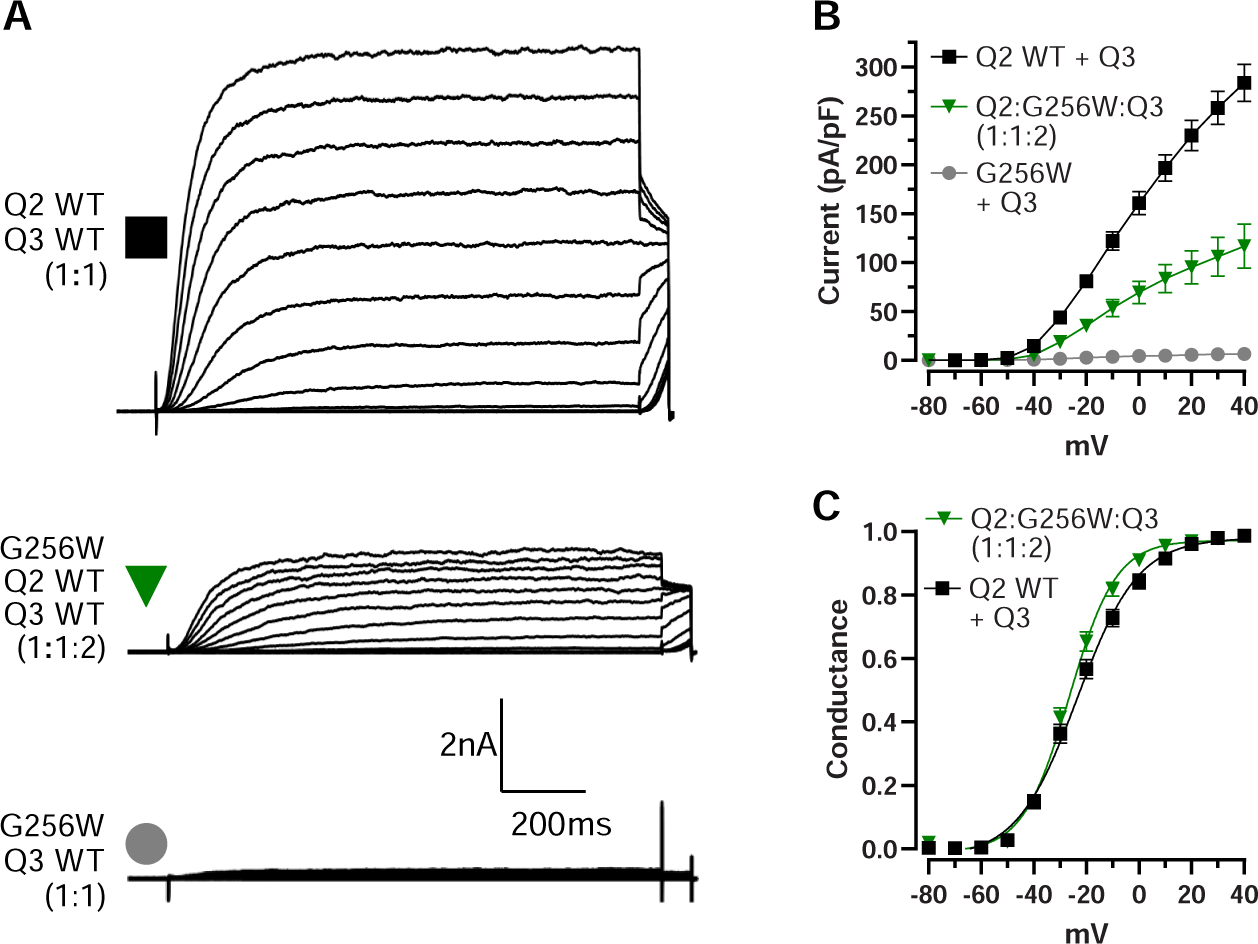
KCNQ2 G256W co-expression suppresses current in KCNQ2/KCNQ3 heteromeric channels recorded by manual patch-clamp. **A.** Representative current families for the indicated ratios of subunits. Note currents are larger than in Figure 3. **B-C.** Current/voltage and conductance/voltage relationships for the indicated WT only and G256W/WT cells.

**Figure 3—figure supplement 2.**
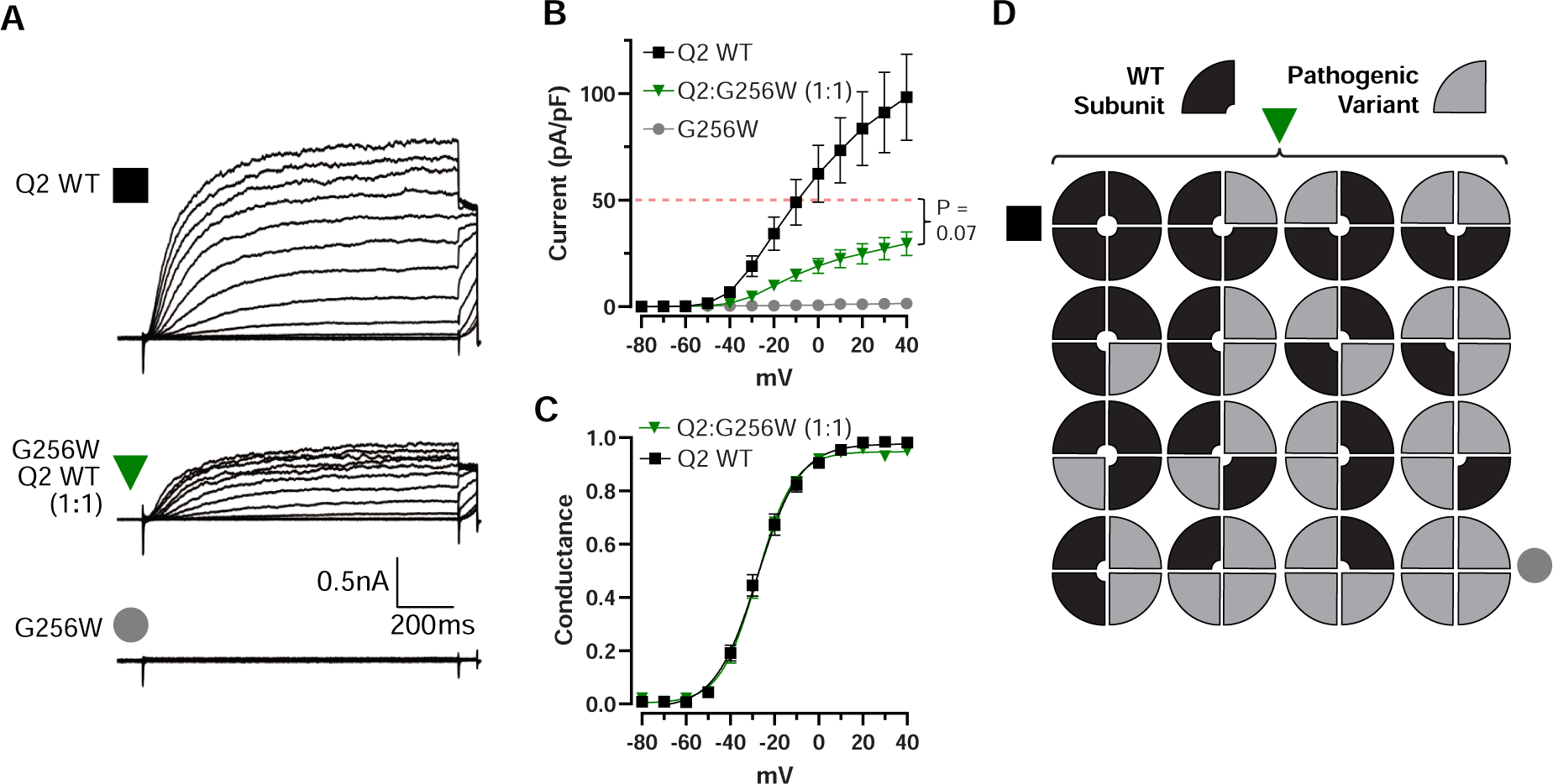
KCNQ2 G256W co-expression suppresses current in KCNQ2 homomeric channels recorded by manual patch-clamp. **A.** Representative current families for the indicated ratios of subunits. Homomeric currents are smaller than in Figure 3—Figure supplement 1. **B-C.** Current/voltage and conductance/voltage relationships for the indicated WT only and G256W/WT cells. **D.** Cartoon showing the expected combinations of WT and G256W subunits under heterozygosity based on a simple random association model. Mutant subunits are included in 15/16 of channel tetramers.

**Figure 4—figure supplement 1.**
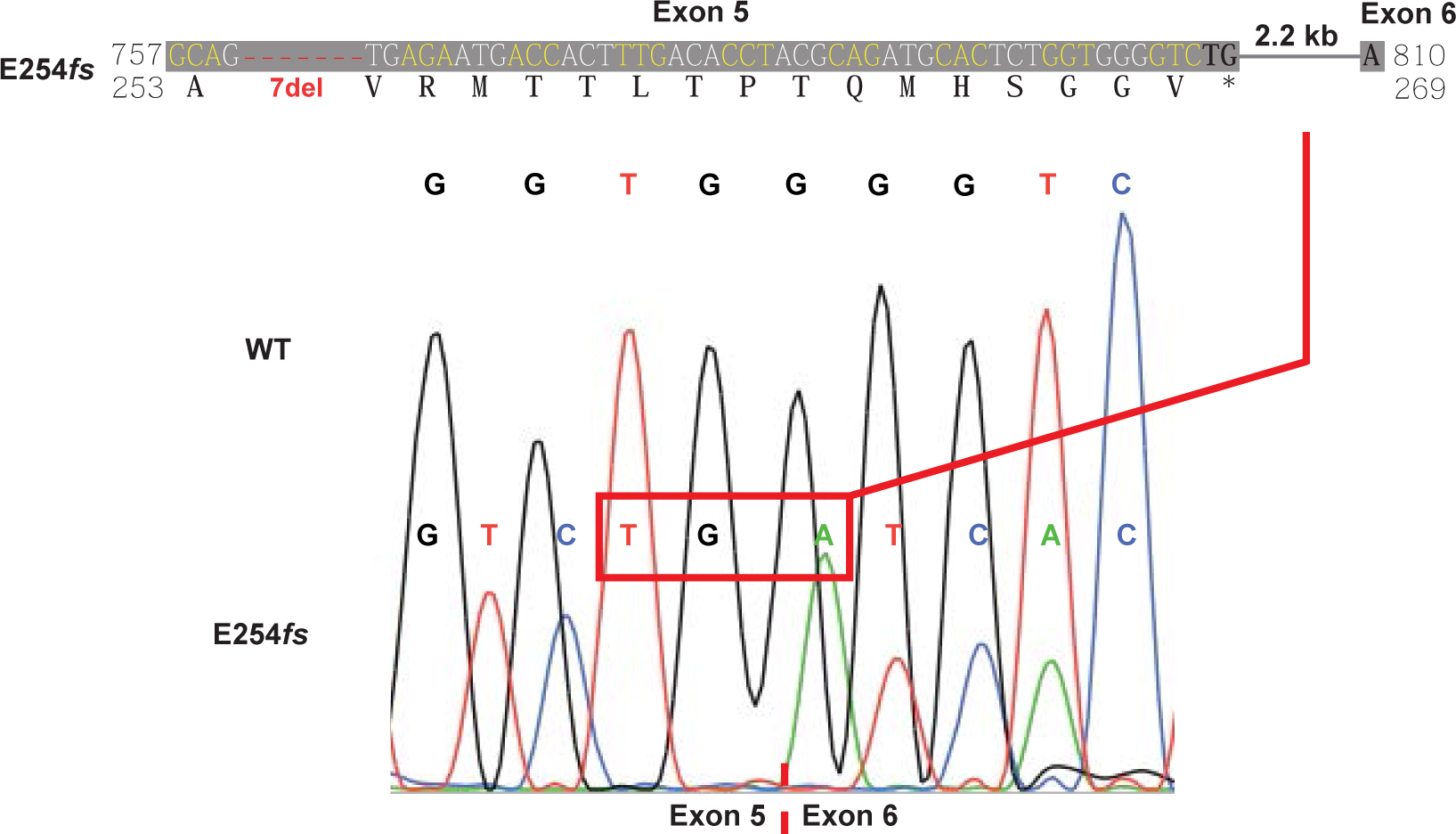
DNA, RNA, and predicted protein consequences of the G256W and E254*fs*\*16 mutations. Upper, DNA and predicted protein alignment of the frameshift mutation. Lower, Sanger sequence for cDNA from hippocampal mRNA of an E254*fs*/+ mouse. Splicing occurs at the WT junction, resulting in the predicted in-frame stop codon.

**Figure 4—figure supplement 2.**
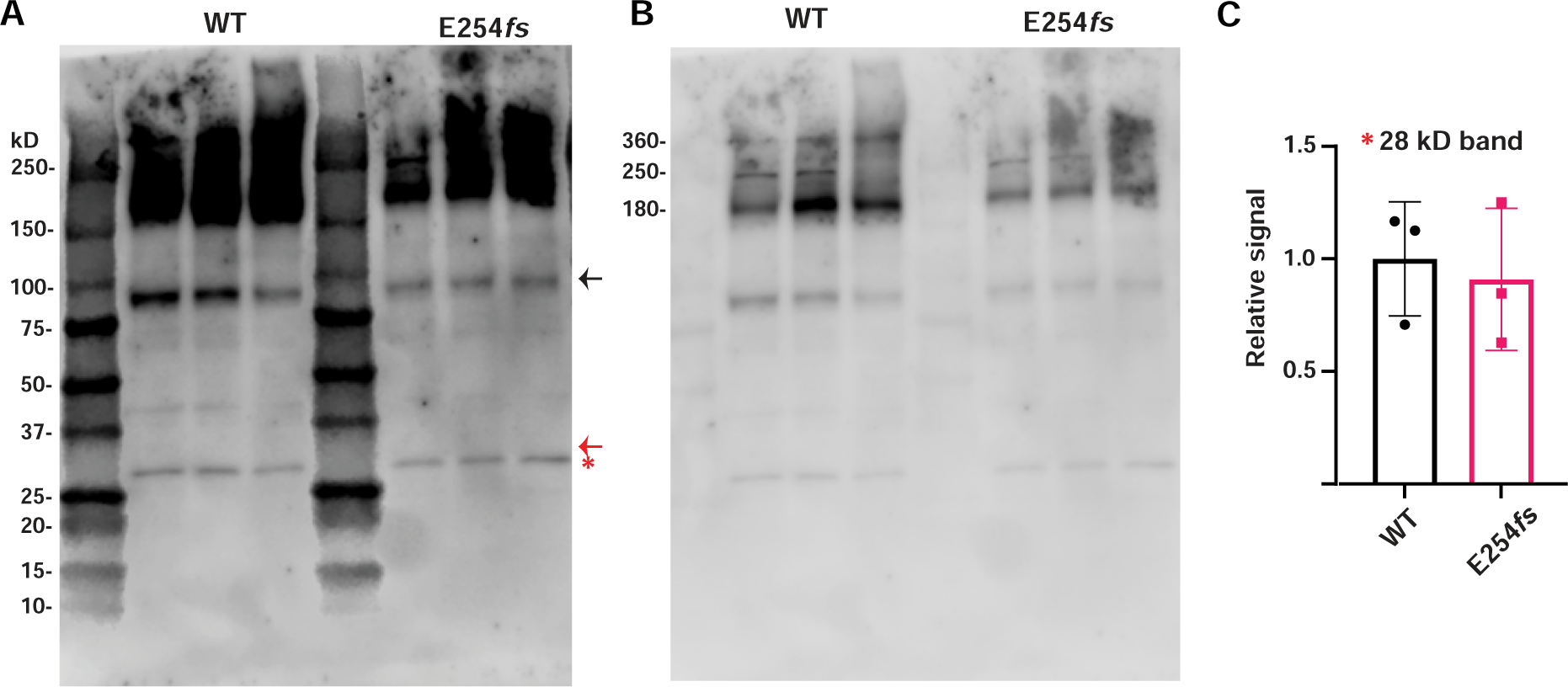
Western blotting reveals no evidence of the predicted E254*fs* truncated protein product. **A.** Western blot of WT and E254*fs*/+ cortical homogenates (3 biological replicates per genotype, all males), probed with KCNQ2 N-terminal antibody. Black arrow indicates the monomer (Mr ∼85kDa), red arrow indicates the estimated relative mobility (∼29.7 kDa), of the truncated protein product made from the E254*fs* allele. Red asterisk indicates a ∼28 kDa band equally detected in both WT and E254*fs*/+. **B.** Same blot as in **A** but windowed to show higher molecular weight bands. Bands at ∼180 kDa and ∼360 kDa consistent with predicted mobility of KCNQ2 dimers and tetramers, respectively. A band at ∼250 kDa appears in all immunoblots of whole brain homogenates using our KCNQ2 N-terminal antibody. Nano-LC tandem mass spectrometry of peptides from an in-gel tryptic digest of this band showed high peptide counts for multiple abundant proteins and few KCNQ2 peptides (Supplementary Data). **C.** Quantification of ∼28 kDa band from **A.**

**Figure 4—figure supplement 3.**
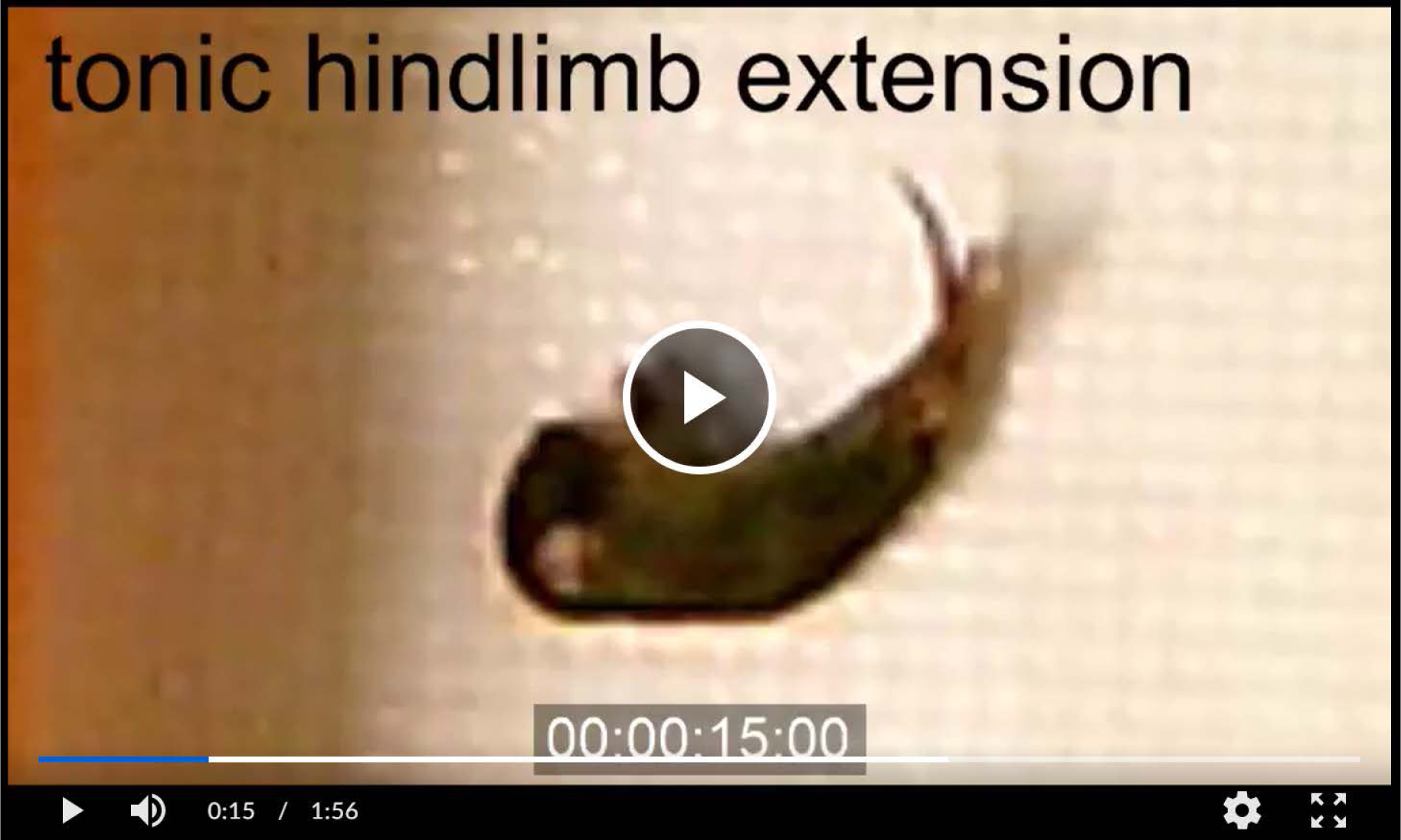
Generalized seizure in a P10 heterozygous G256W mouse. This movie includes from 9:01 to 11:05 of a 15:00 min period of open field observation. Animal recovers upright posture at 1:45 in the clip. <u>Link to movie F4-S3</u>

**Figure 5—figure supplement 1.**
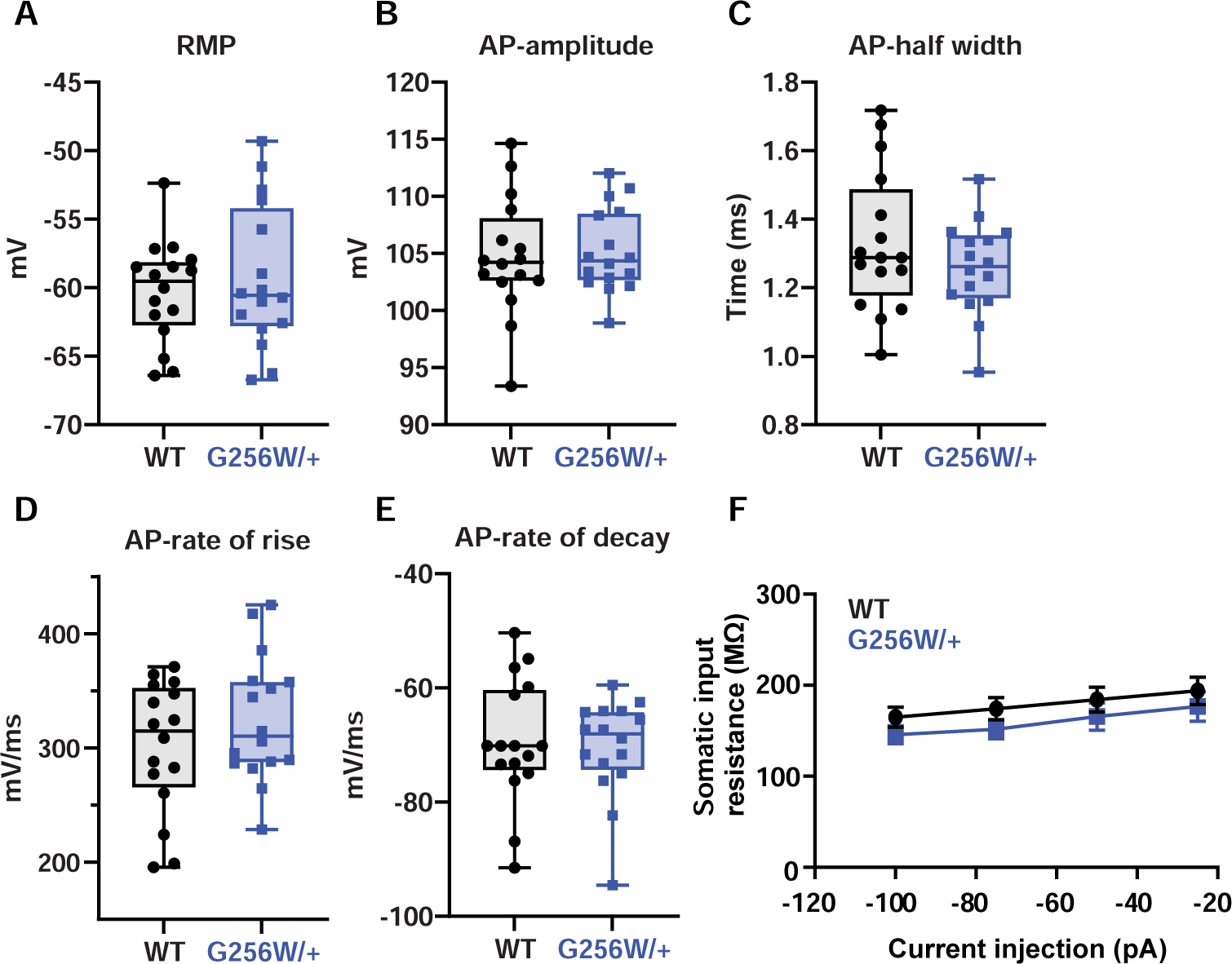
Several neuronal biophysical properties are unchanged. **A.** Resting membrane potential. **B.** Action potential amplitude. **C.** Action potential width. **D.** Action potential rise slope. **E.** Action potential decay. **F.** Input resistance. For all panels, 3 animals per genotype, n=16 cells/genotype).

**Figure 6—figure supplement 1.**
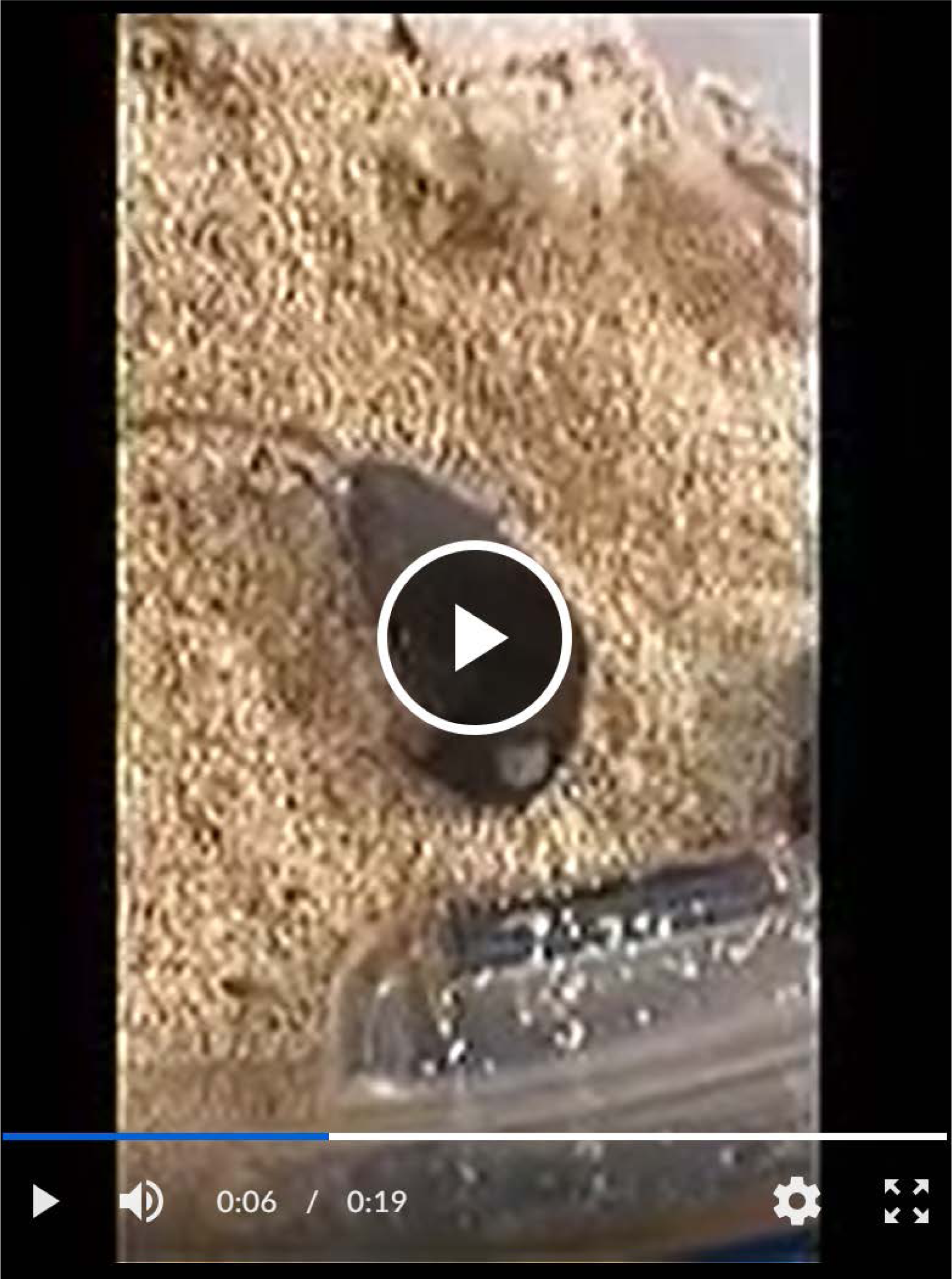
Video of a fatal convulsive seizure in a 4 month old heterozygous G256W mouse. Seizure onset (not shown) occurred 5-10 sec prior to the start of recording with wild running and jumping, followed by arrest, then resumed (start of video). This was again followed by loss of postural control, followed by sustained forelimb flexor/hindlimb extensor posturing. Attempts to resuscitate the animal were begun immediately and were unsuccessful. <u>Link to movie F6-S1</u>

**Figure 6—figure supplement 2.**
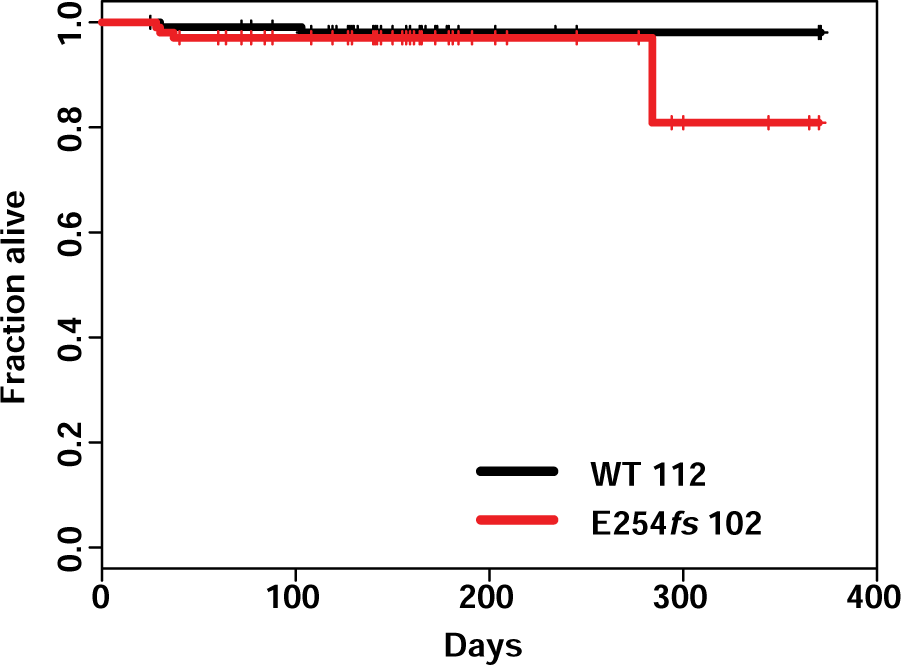
No significant mortality in heterozygous E254*fs* mice. Survival curve of WT vs E254*fs*/+ mice, hashmarks indicate censored mice. E254*fs*/+ mice showed no significant mortality, P = 0.452 Cox propotional hazards model.

**Figure 7—figure supplement 1.**
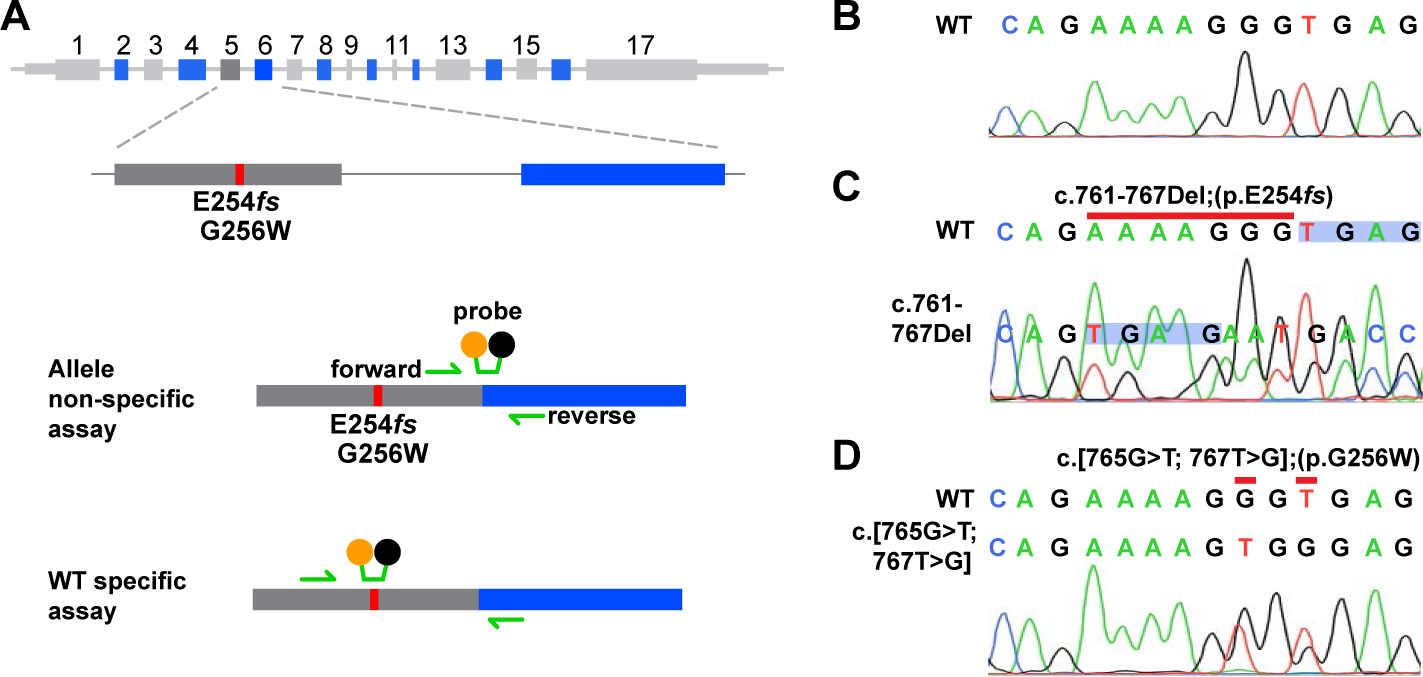
**A.** Probes used for allele non-selective and WT allele-selective RT-qPCR. **B.** Upper, WT sequence (bases 758-771). Lower, Sanger trace of amplified *Kcnq2* cDNA from WT hippocampus. **C.** Upper, WT sequence; the red line indicates bases deleted in the E254*fs* allele. Lower, Sanger trace of cDNA from an E254*fs*/+ mouse. Blue shading highlights the shift following the deletion at positions 761-767. Peaks corresponding to E254*fs* transcripts are labeled and are smaller than WT peaks. **D.** Upper, alignment of WT and missense variant DNA sequences, red lines highlight the two base substitutions at codon 256. Below, Sanger trace of amplified cDNA from G256W/+ hippocampus. Double peaks are visible at the bases mutated by Crispr.

**Figure 8—figure supplement 1.**
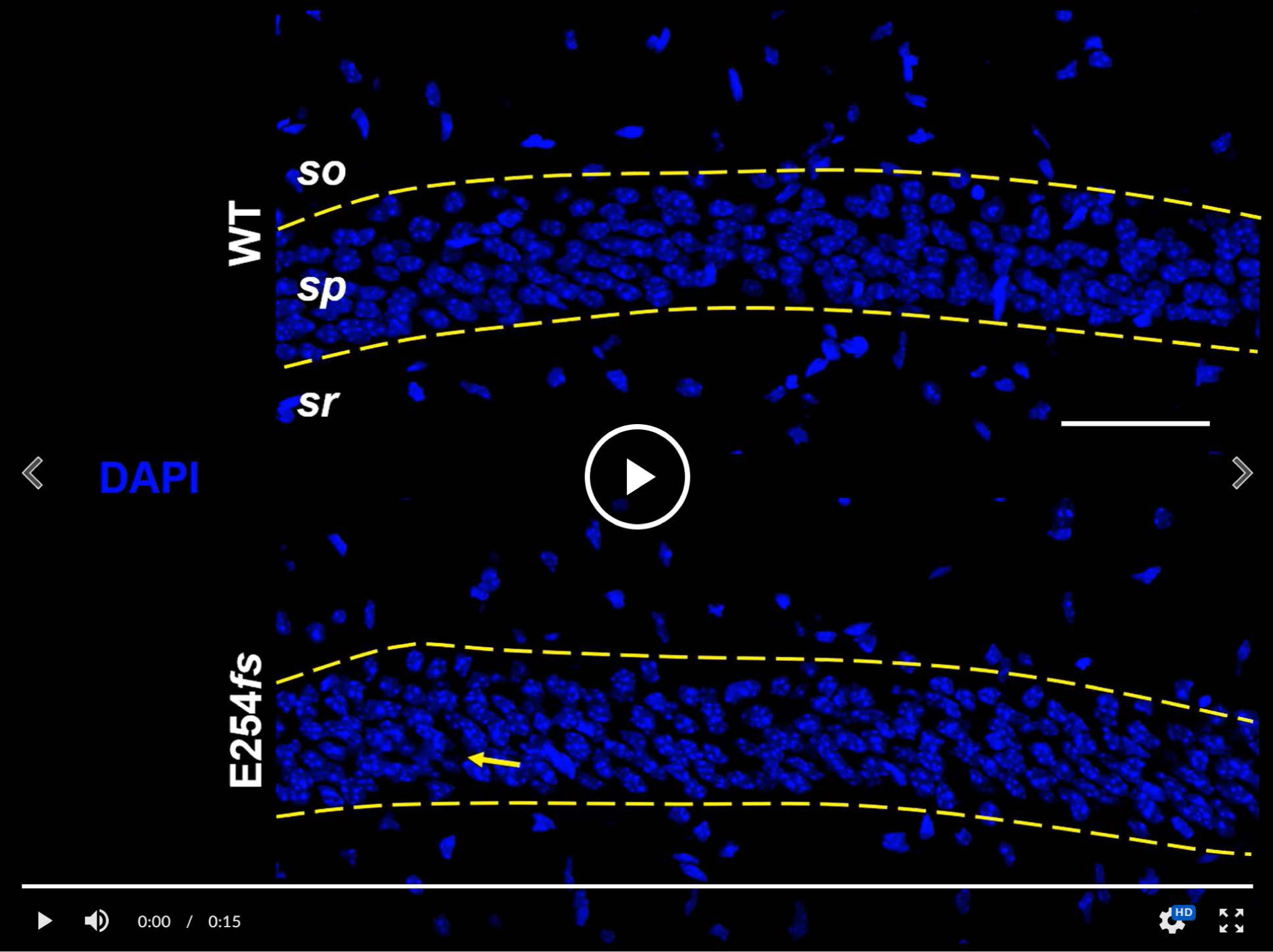
In CA1, the KCNQ2 and KCNQ3 cellular and subcellular immunolabeling patterns appear similar for WT and heterozygous E254*fs* mice. Ankyrin-G marks position of AISs. KCNQ2 and KCNQ3 strongly label CA1 AISs in E254*fs*/+ mice, and do not show increased somatic labeling compared to WT. Highlighted by an arrow is one interneuron in stratum pyramidale that was somatically labeled for KCNQ2 only. Scale: 50 μm. <u>Link to movie F8-</u> <u>S1</u>

**Figure 8—figure supplement 2.**
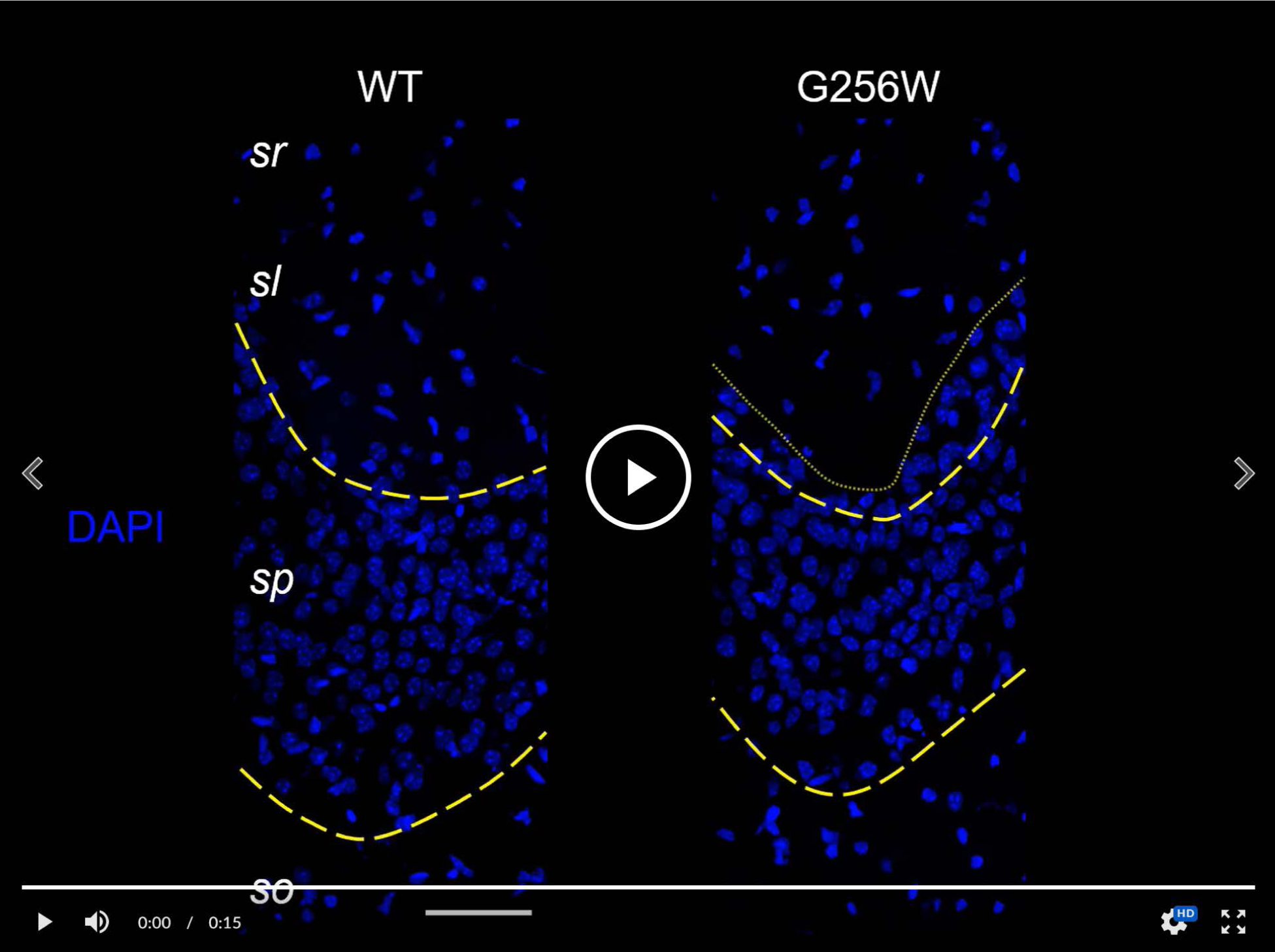
Heterozygous G256W mice show increased CA3 pyramidal cell somatic labeling and reduced mossy fiber labeling for KCNQ2 and KCNQ3. Yellow lines demarcate the borders of sp; the sp-sl border is cut obliquely through the tissue section in the G256W/+ sample. PanNav strongly labels the unmylenated axons of the mossy fibers in stratum lucidum of both samples. PanNav also labels the obliquely cut AISs of pyramidal cell neurons, which are mostly located within sp. Scale: 50 μm. <u>Link to movie F8-S2</u>

**Figure 8—figure supplement 3.**
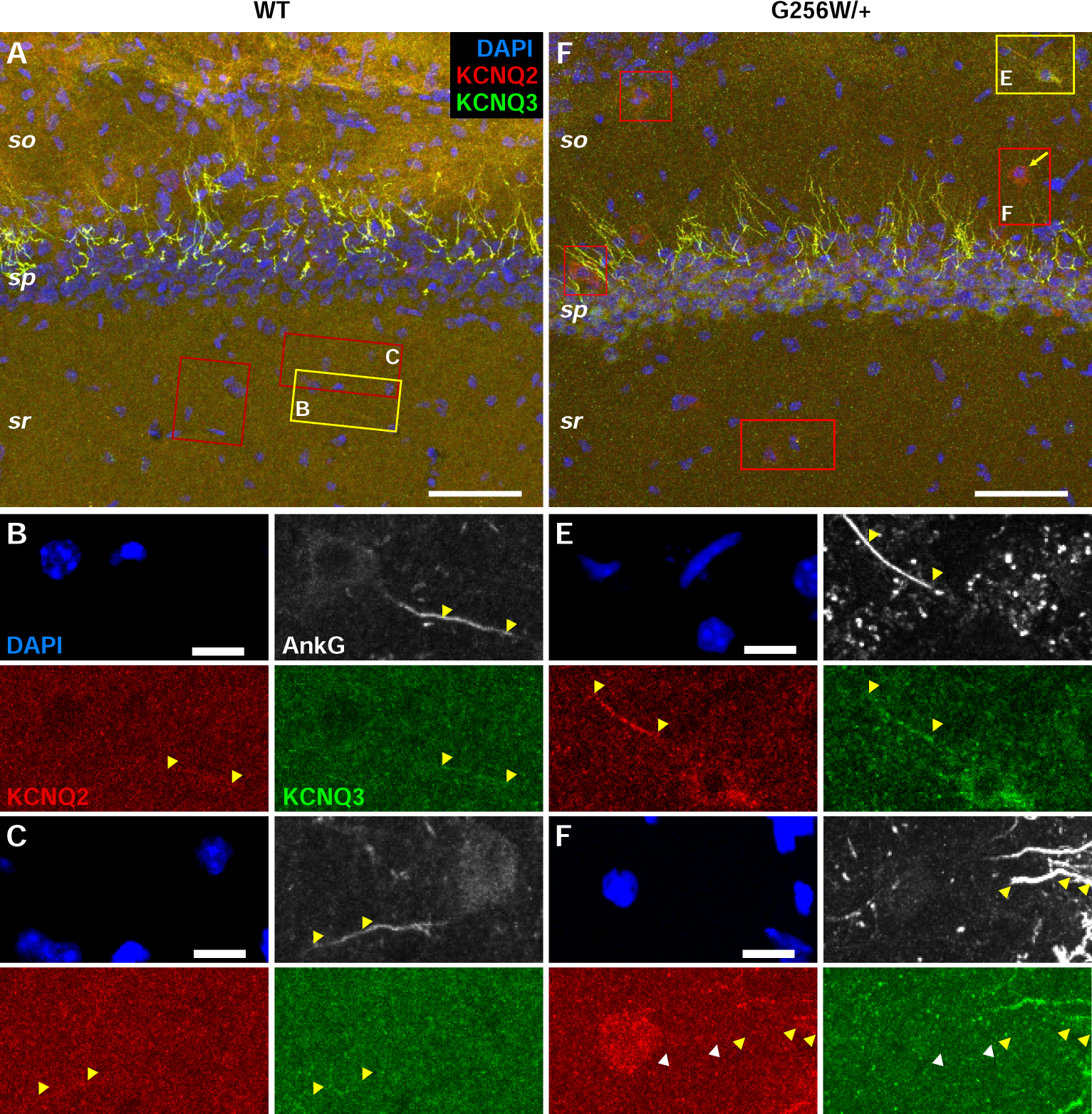
G256W/+ mouse Interneurons in CA1 show somatic KCNQ2 labeling. **A, D.** Wider views of CA, including more of S. radiatum and s. oriens. (**D**, same as shown in Figure 8-Movie). In WT, positions of three S. radiatum interneurons are boxed, but higher magnification (e.g., **B, C**) shows lack somatic labeling for KCNQ2 or KCNQ3 In G256W/+ image (D), four interneurons somatically labelled for KCNQ2 but not KCNQ3 are enclosed by red boxes. The interneuron indicated with an arrow in Figure 8-Movie (KCNQ2 labeled, KCNQ3 unlabeled), is again highlighted. The yellow box encloses an interneuron somatically co-labeled for KCNQ2 and KCNQ3. **E.** Individual laser channels for the interneuron enclosed by yellow box in **D**. The soma is labeled for both KCNQ2 and KCNQ3. The nearby AIS showing AnkG, KCNQ2, and KCNQ3 may arise from this or a different cell, as its origin was not verified by higher resolution re-imaging. **F.** Interneuron somatically labeled for KCNQ2, not KCNQ3. Its AIS appears to arise from a KCNQ2 labeled (white arrowheads). In C-F, yellow arrowheads show distal AISs strongly labeled for AnkG, and weakly for KCNQ2 and KCNQ3. Scales: A, 50 µm; B, 10 µm.

**Figure 8—figure supplement 4.**
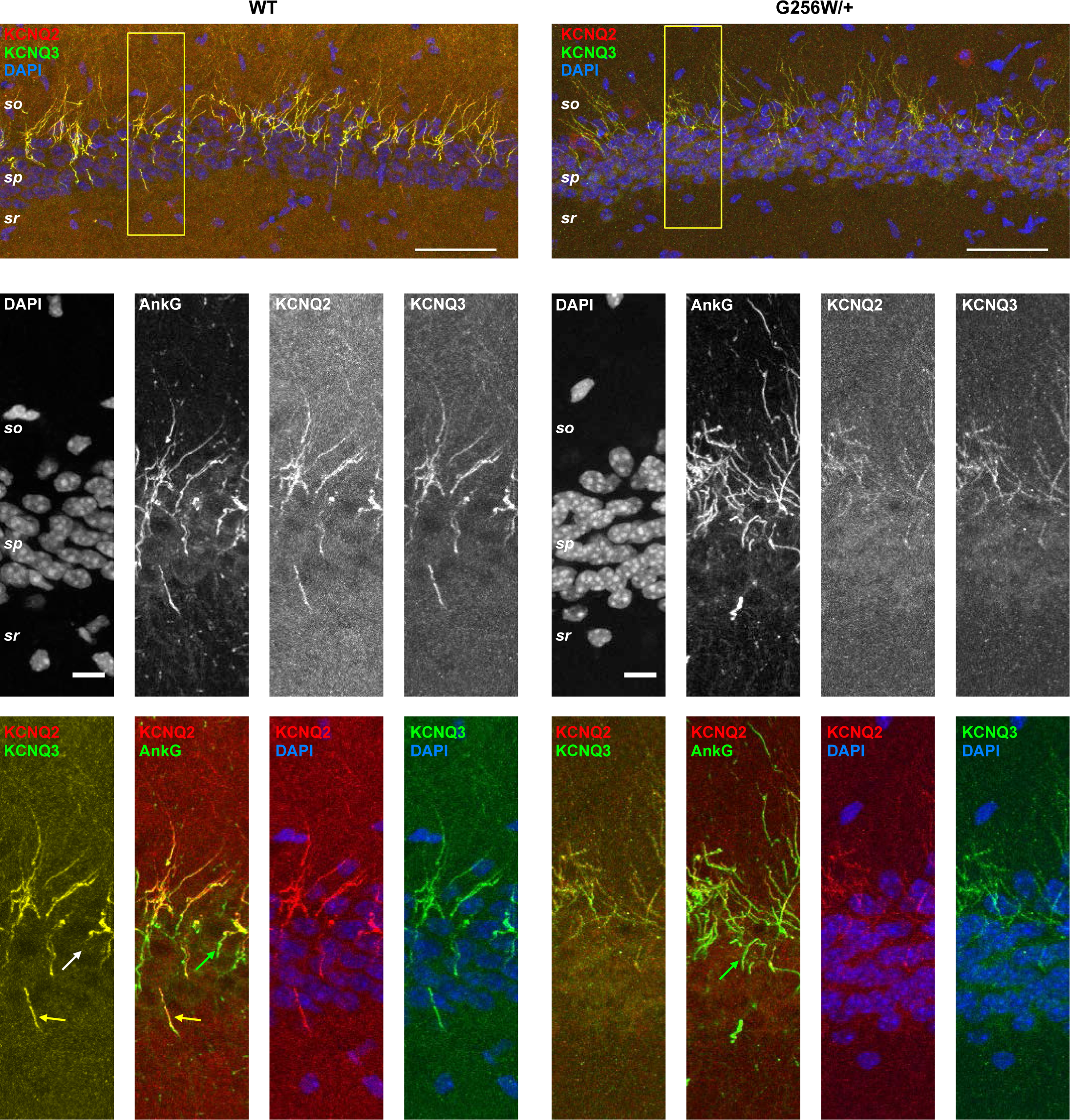
Single marker grey-scale images and selected merged images of CA1, related to Figure 8 movie. In lower merge images, yellow arrows indicate KCNQ2/KCNQ3 overlap, white and green arrows indicate AnkG-only labeling of proximal AIS. Scales: 50 µm, upper; 10 µm, middle and lower.

**Figure 8—figure supplement 5.**
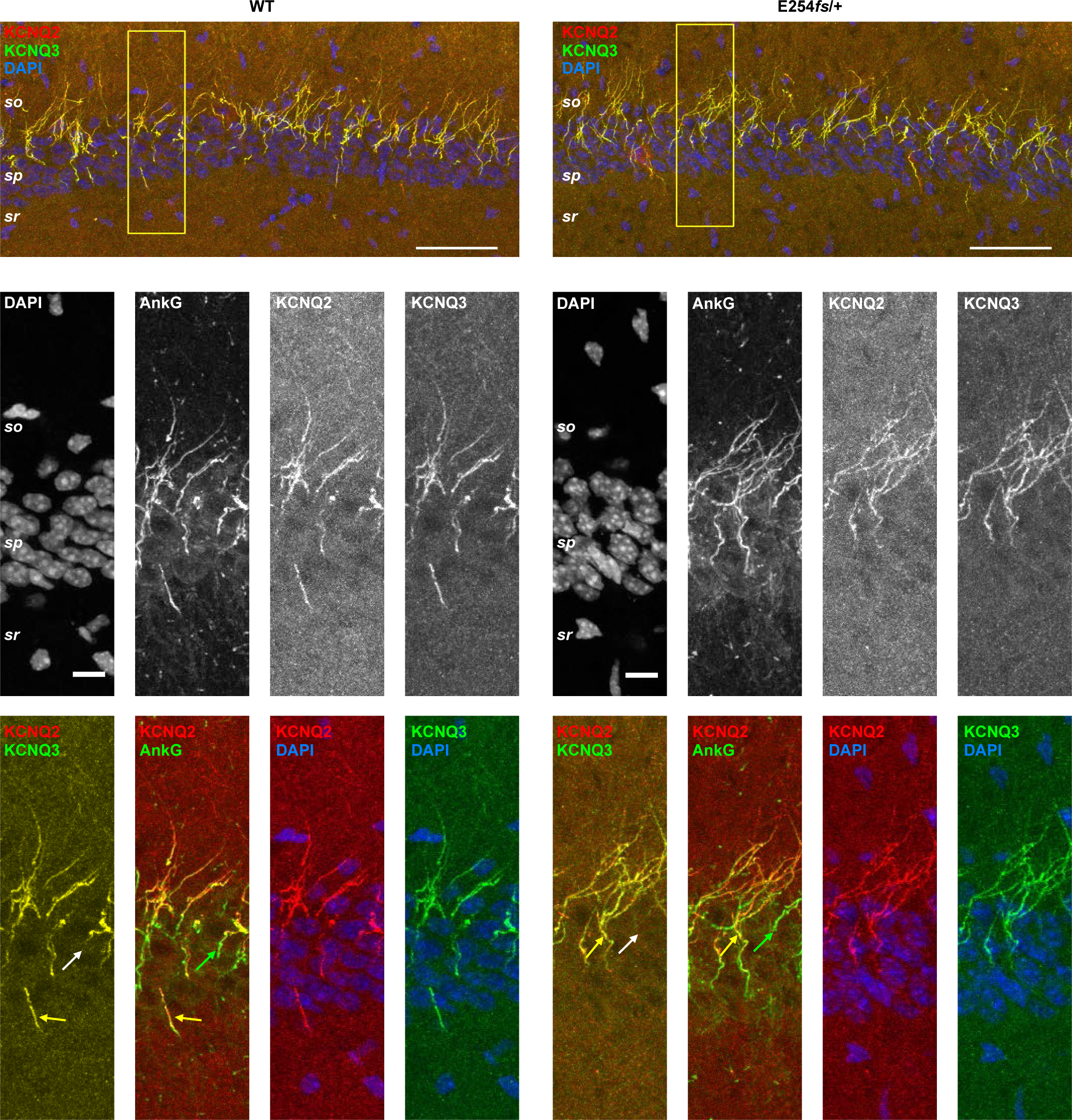
Single marker grey-scale images and selected merged images of CA1, related to Figure 8 —figure supplement 1 movie. In lower merge images, yellow arrows indicate KCNQ2/KCNQ3 overlap, white and green arrows indicate AnkG-only labeling of proximal AIS. Scales: 50 µm, upper; 10 µm, middle and lower.

**Figure 8—figure supplement 6.**
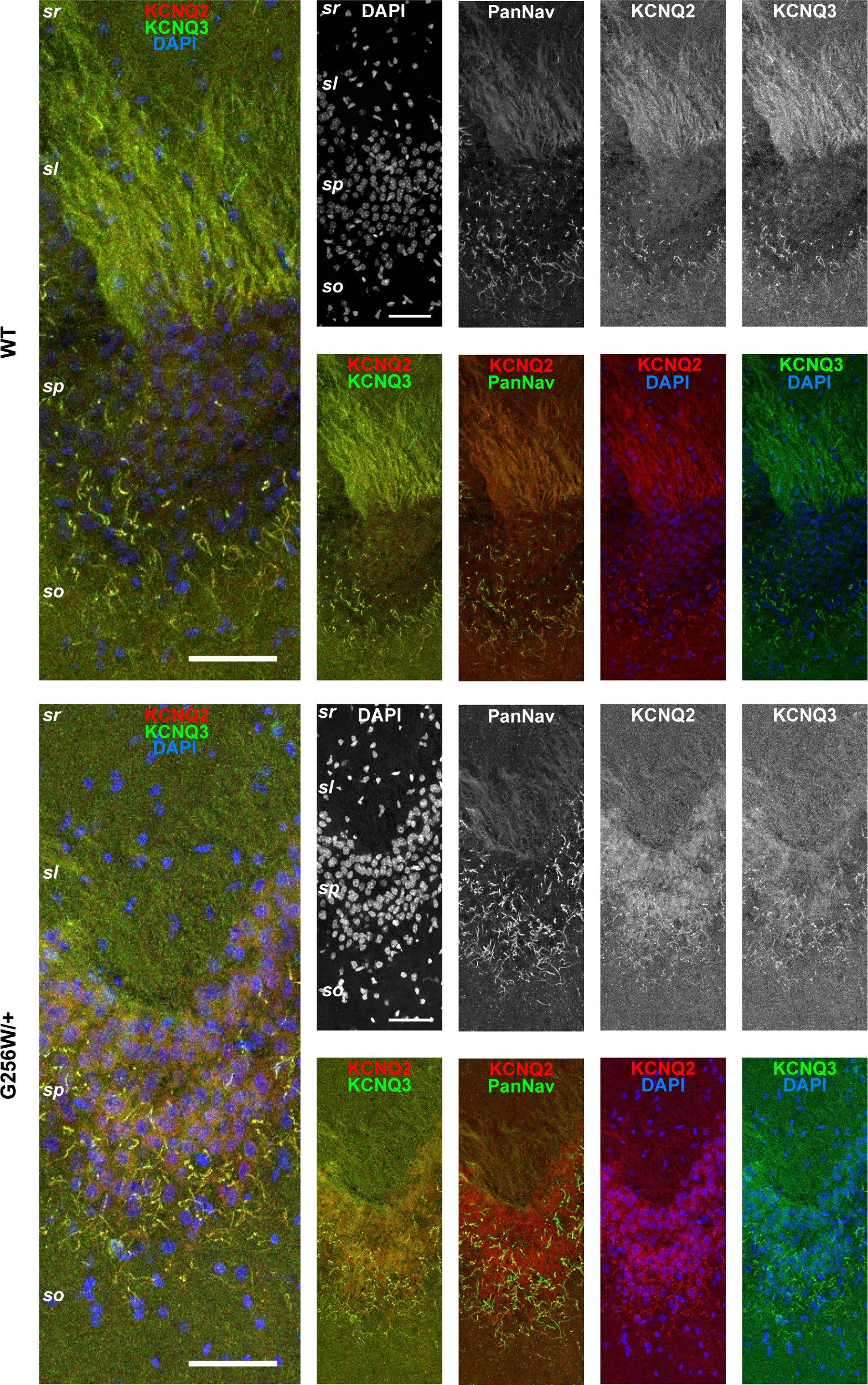
Single marker grey-scale images and selected merged images of CA3, related to Figure 8 —figure supplement 2 movie. Scales: 50 µm.

**Figure 10—figure supplement 1.**
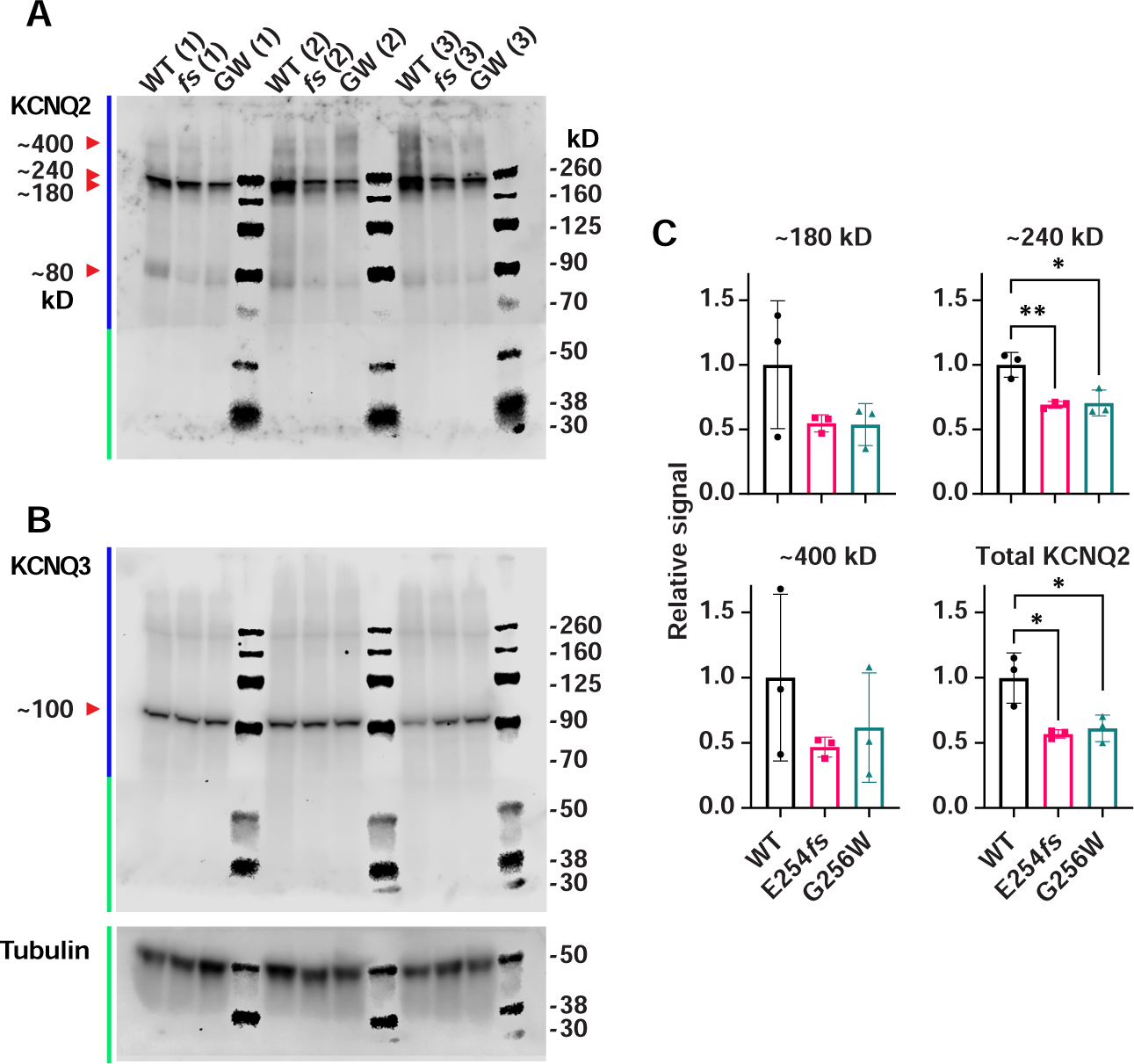
KCNQ2 antibodies, unlike KCNQ3, show complex electrophoretic banding pattern, and reduced levels in E254*fs* and G256W mice. PVDF filter with electrotransferred brain proteins was cut between the 70 and 50 kDa markers. Each lane contains homogenate from an individual animal. Upper portion of the filter (blue bar at L) was initially probed for KCNQ2; lower portion (green bar at L) was initially probed for tubulin, then stripped and reprobed for KCNQ2. Next, the entire filter was stripped and probed again for KCNQ3. **A.** KCNQ2 blots. Arrowheads point to KCNQ2 monomer band (Mr ∼80 kDa), and candidate dimeric and oligomeric bands of Mr ∼180 to ∼400 kDa. **B.** Sequential probe of same filter from A using KCNQ3 and tubulin antibodies. Arrowhead points to predicted KCNQ3 monomer with a Mr ∼100 kDa. **C.** The indicated bands from the KCNQ2 blots, and the total Individual lane intensities, means of genotypes, and SEMs are shown. One way ANOVA, * = P≤0.05.

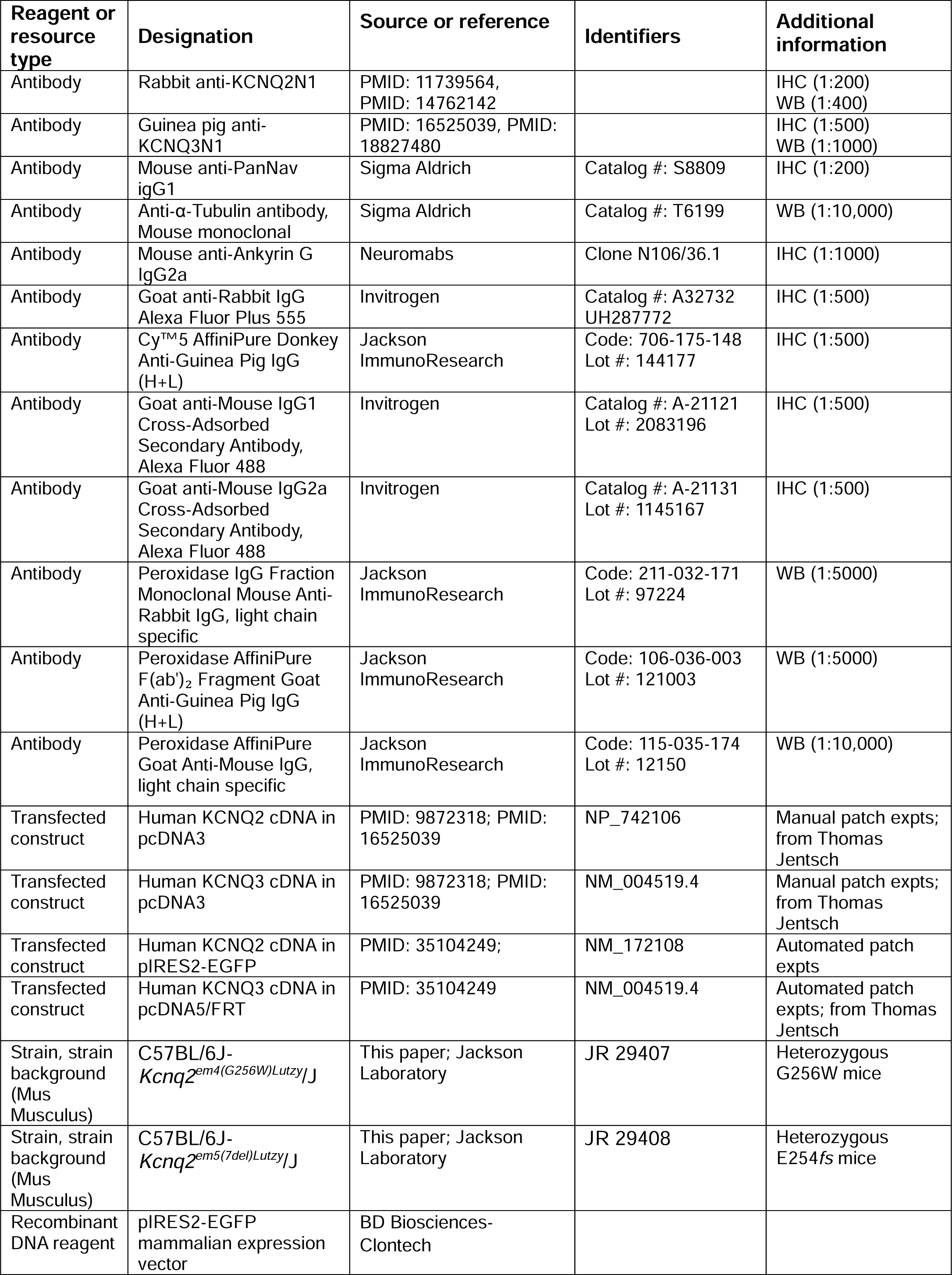
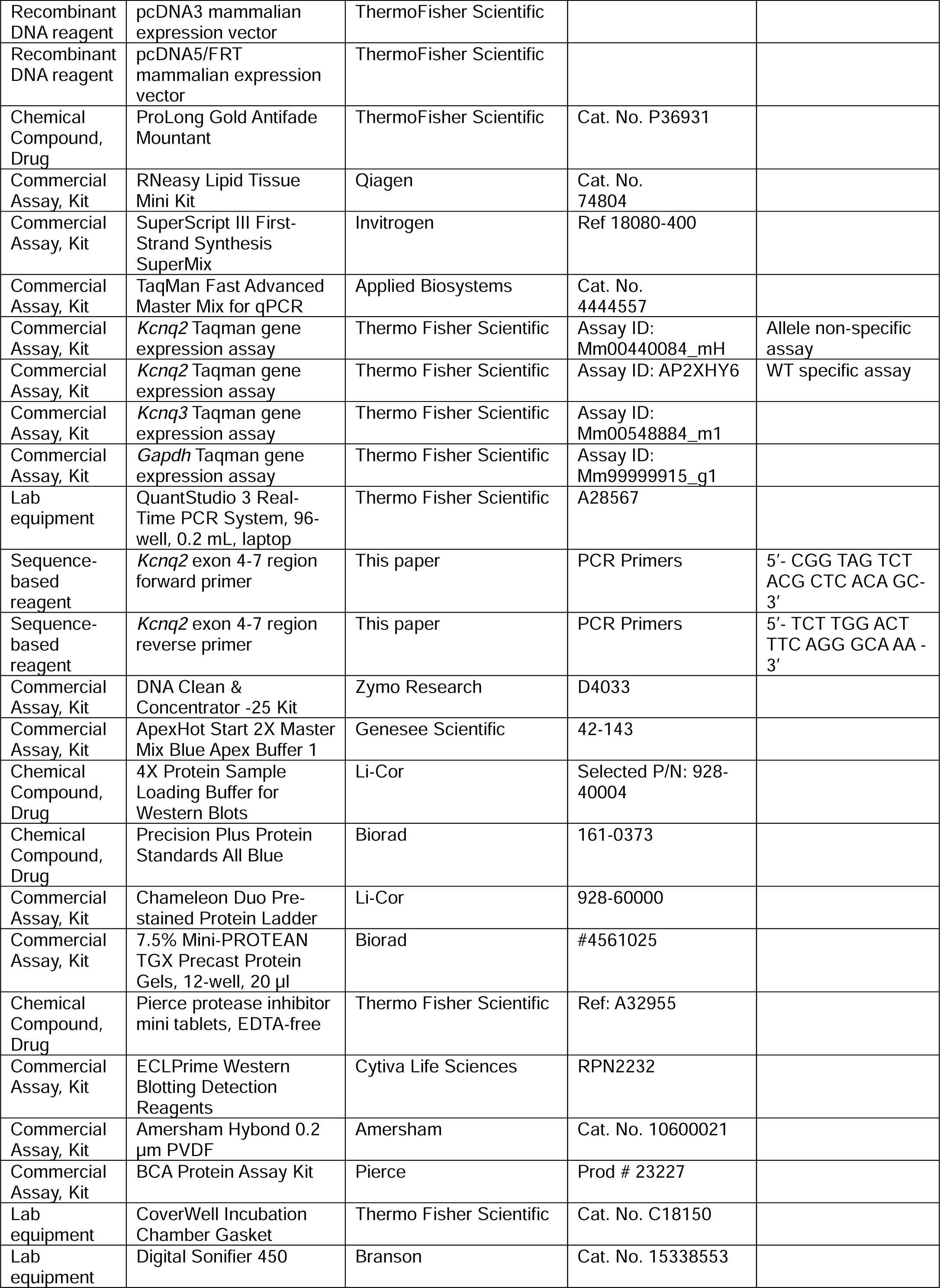
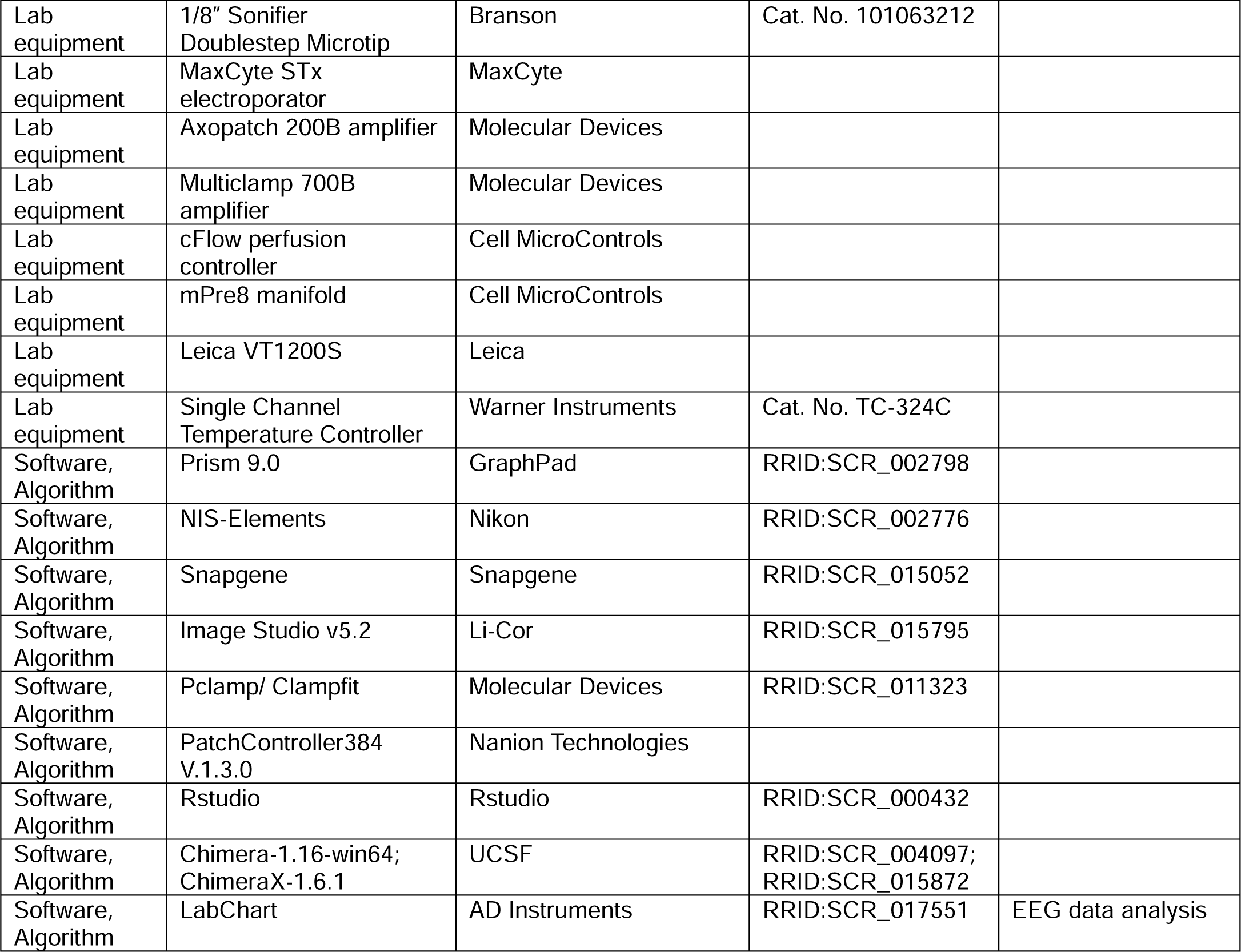

## References

1. Aiba, I., and Noebels, J.L. (2021). Kcnq2/Kv7.2 controls the threshold and bi-hemispheric symmetry of cortical spreading depolarization. Brain 144, 2863–2878. 10.1093/brain/awab141.

2. Banerjee, A., Lee, A., Campbell, E., and Mackinnon, R. (2013). Structure of a pore-blocking toxin in complex with a eukaryotic voltage-dependent K(+) channel. Elife 2, e00594. 10.7554/eLife.00594.

3. Bass, J.S., Tuo, A.H., Ton, L.T., Jankovic, M.J., Kapadia, P.K., Schirmer, C., and Krishnan, V. (2020). On the digital psychopharmacology of valproic acid in mice. Front Neurosci 14, 594612. 10.3389/fnins.2020.594612.

4. Battefeld, A., Tran, B.T., Gavrilis, J., Cooper, E.C., and Kole, M.H. (2014). Heteromeric Kv7.2/7.3 channels differentially regulate action potential initiation and conduction in neocortical myelinated axons. J Neurosci 34, 3719–3732. 10.1523/JNEUROSCI.4206-13.2014.

5. Beal, J.C., Cherian, K., and Moshe, S.L. (2012). Early-onset epileptic encephalopathies: Ohtahara syndrome and early myoclonic encephalopathy. Pediatric neurology 47, 317–323. 10.1016/j.pediatrneurol.2012.06.002.

6. Berg, A.T., Mahida, S., and Poduri, A. (2021). KCNQ2-DEE: developmental or epileptic encephalopathy? Ann Clin Transl Neurol 8, 666–676. 10.1002/acn3.51316.

7. Biba-Maazou, N., Becq, H., Pallesi-Pocachard, E., Sarno, S., Granjeaud, S., Montheil, A., Kurz, M., Villard, L., Milh, M., Santini, P.L., and Aniksztejn, L. (2022). Time-limited alterations in cortical activity of a knock-in mouse model of KCNQ2-related developmental and epileptic encephalopathy. J Physiol 600, 2429–2460. 10.1113/JP282536

8. Bosselmann, C.M., Hedrich, U.B.S., Muller, P., Sonnenberg, L., Parthasarathy, S., Helbig, I., Lerche, H., and Pfeifer, N. (2022). Predicting the functional effects of voltage-gated potassium channel missense variants with multi-task learning. EBioMedicine 81, 104115. 10.1016/j.ebiom.2022.104115.

9. Brown, D.A., and Adams, P.R. (1980). Muscarinic suppression of a novel voltage-sensitive K+ current in a vertebrate neuron. Nature 283, 673–676.

10. Brunger, T., Perez-Palma, E., Montanucci, L., Nothnagel, M., Moller, R.S., Schorge, S., Zuberi, S., Symonds, J., Lemke, J.R., Brunklaus, A., et al. (2023). Conserved patterns across ion channels correlate with variant pathogenicity and clinical phenotypes. Brain 146, 923–934. 10.1093/brain/awac305.

11. Carugo, O., and Djinovic-Carugo, K. (2013). Half a century of Ramachandran plots. Acta Crystallogr D Biol Crystallogr 69, 1333–1341. 10.1107/S090744491301158X.

12. Chung, H.J., Jan, Y.N., and Jan, L.Y. (2006). Polarized axonal surface expression of neuronal KCNQ channels is mediated by multiple signals in the KCNQ2 and KCNQ3 C-terminal domains. Proc Natl Acad Sci U S A 103, 8870–8875.

13. Cooper, E.C. (2011). Made for "anchorin": Kv7.2/7.3 (KCNQ2/KCNQ3) channels and the modulation of neuronal excitability in vertebrate axons. Seminars in cell & developmental biology 22, 185–192. 10.1016/j.semcdb.2010.10.001.

14. Cooper, E.C., Aldape, K.D., Abosch, A., Barbaro, N.M., Berger, M.S., Peacock, W.S., Jan, Y.N., and Jan, L.Y. (2000). Colocalization and coassembly of two human brain M-type potassium channel subunits that are mutated in epilepsy. Proceedings of the National Academy of Sciences 97, 4914. 10.1073/pnas.090092797.

15. Cooper, E.C., Harrington, E., Jan, Y.N., and Jan, L.Y. (2001). M-channel KCNQ2 subunits are localized to key sites for control of neuronal network oscillations and synchronization in mouse brain. J Neurosci 21, 9529–9540.

16. Coppola, G., Castaldo, P., Miraglia del Giudice, E., Bellini, G., Galasso, F., Soldovieri, M.V., Anzalone, L., Sferro, C., Annunziato, L., Pascotto, A., and Taglialatela, M. (2003). A novel KCNQ2 K+ channel mutation in benign neonatal convulsions and centrotemporal spikes. Neurology 61, 131–134. 10.1212/01.wnl.0000069465.53698.bd.

17. Creson, T.K., Rojas, C., Hwaun, E., Vaissiere, T., Kilinc, M., Jimenez-Gomez, A., Holder, J.L., Jr., Tang, J., Colgin, L.L., Miller, C.A., and Rumbaugh, G. (2019). Re-expression of SynGAP protein in adulthood improves translatable measures of brain function and behavior. Elife 8. 10.7554/eLife.46752.

18. Devaux, J.J., Kleopa, K.A., Cooper, E.C., and Scherer, S.S. (2004). KCNQ2 is a nodal K^+^ channel. J Neurosci 24, 1236–1244.

19. Dirkx, N., Miceli, F., Taglialatela, M., and Weckhuysen, S. (2020). The Role of Kv7.2 in Neurodevelopment: Insights and Gaps in Our Understanding. Front Physiol 11, 570588. 10.3389/fphys.2020.570588.

20. Donnan, A.M., Schneider, A.L., Russ-Hall, S., Churilov, L., and Scheffer, I.E. (2023). Rates of Status Epilepticus and Sudden Unexplained Death in Epilepsy in People With Genetic Developmental and Epileptic Encephalopathies. Neurology 100, e1712–e1722. 10.1212/WNL.0000000000207080.

21. Doyle, D.A., Cabral, J.M., Pfuetzner, R.A., Kuo, A., Gulbis, J.M., Cohen, S.L., Chait, B.T., and MacKinnon, R. (1998). The structure of the potassium channel: molecular basis of K^+^ conduction and selectivity. Science 280, 69–77.

22. Dyle, M.C., Kolakada, D., Cortazar, M.A., and Jagannathan, S. (2020). How to get away with nonsense: Mechanisms and consequences of escape from nonsense-mediated RNA decay. Wiley Interdiscip Rev RNA 11, e1560. 10.1002/wrna.1560.

23. FDA, U.S. (2013). FDA Drug Safety Communication: FDA approves label changes for anti-seizure drug Potiga (ezogabine) describing risk of retinal abnormalities, potential vision loss, and skin discoloration. http://www.fda.gov/Drugs/DrugSafety/ucm372774.htm.

24. Fernandez-Marino, A.I., Tan, X.F., Bae, C., Huffer, K., Jiang, J., and Swartz, K.J. (2023). Inactivation of the Kv2.1 channel through electromechanical coupling. Nature 622, 410–417. 10.1038/s41586-023-06582-8.

25. Fidzinski, P., Korotkova, T., Heidenreich, M., Maier, N., Schuetze, S., Kobler, O., Zuschratter, W., Schmitz, D., Ponomarenko, A., and Jentsch, T.J. (2015). KCNQ5 K^+^ channels control hippocampal synaptic inhibition and fast network oscillations. Nat Commun 6, 6254. 10.1038/ncomms7254.

26. Goddard, T.D., Huang C.C., Meng, E.C, Pettersen, E.F., Couch, G.S., Morris, J.H., Ferrin, T.E. (2018). UCSF ChimeraX: Meeting modern challenges in visualization and analysis. Protein Sci 1, 4–25.

27. Goto, A., Ishii, A., Shibata, M., Ihara, Y., Cooper, E.C., and Hirose, S. (2019). Characteristics of KCNQ2 variants causing either benign neonatal epilepsy or developmental and epileptic encephalopathy. Epilepsia 60, 1870–1880. 10.1111/epi.16314.

28. Grinton, B.E., Heron, S.E., Pelekanos, J.T., Zuberi, S.M., Kivity, S., Afawi, Z., Williams, T.C., Casalaz, D.M., Yendle, S., Linder, I., et al. (2015). Familial neonatal seizures in 36 families: Clinical and genetic features correlate with outcome. Epilepsia 56, 1071–1080. 10.1111/epi.13020.

29. Gunthorpe, M.J., Large, C.H., and Sankar, R. (2012). The mechanism of action of retigabine (ezogabine), a first-in-class K+ channel opener for the treatment of epilepsy. Epilepsia 53, 412–424. 10.1111/j.1528-1167.2011.03365.x.

30. Hadley, J.K., Passmore, G.M., Tatulian, L., Al-Qatari, M., Ye, F., Wickenden, A.D., and Brown, D.A. (2003). Stoichiometry of expressed KCNQ2/KCNQ3 potassium channels and subunit composition of native ganglionic M channels deduced from block by tetraethylammonium. J Neurosci 23, 5012–5019.

31. Hill, J.M., Lim, M.A., and Stone, M.M. (2008). Developmental Milestones in the Newborn Mouse. in I. Gozes, Ed., Neuropeptide Techniques Humana Press, Totowa, 131-149.

32. Hille, B., Armstrong, C.M., and MacKinnon, R. (1999). Ion channels: from idea to reality. Nat Med 5, 1105–1109.

33. Hoshi, T., and Armstrong, C.M. (2013). C-type inactivation of voltage-gated K+ channels: pore constriction or dilation? J Gen Physiol 141, 151–160. 10.1085/jgp.201210888.

34. Hu, W., and Bean, B.P. (2018). Differential Control of Axonal and Somatic Resting Potential by Voltage-Dependent Conductances in Cortical Layer 5 Pyramidal Neurons. Neuron 99, 1355. 10.1016/j.neuron.2018.08.042.

35. Jin, Z., Liang, G.H., Cooper, E.C., and Jarlebark, L. (2009). Expression and localization of K^+^ channels KCNQ2 and KCNQ3 in the mammalian cochlea. Audiol Neurootol 14, 98–105. 10.1159/000158538.

36. Jing, J., Dunbar, C., Sonesra, A., Chavez, A., Park, S., Yang, R., Soh, H., Lee, M., Tzingounis, A.V., Cooper, E.C., et al. (2022). Removal of KCNQ2 from parvalbumin-expressing interneurons improves anti-seizure efficacy of retigabine. Exp Neurol 355, 114141. 10.1016/j.expneurol.2022.114141.

37. Karczewski, K.J., Francioli, L.C., Tiao, G., Cummings, B.B., Alfoldi, J., Wang, Q., Collins, R.L., Laricchia, K.M., Ganna, A., Birnbaum, D.P., et al. (2020). The mutational constraint spectrum quantified from variation in 141,456 humans. Nature 581, 434–443. 10.1038/s41586-020-2308-7.

38. Khemaissa, S., Walrant, A., and Sagan, S. (2022). Tryptophan, more than just an interfacial amino acid in the membrane activity of cationic cell-penetrating and antimicrobial peptides. Q Rev Biophys 55, e10. 10.1017/S0033583522000105.

39. Kim, E.C., Zhang, J., Tang, A.Y., Bolton, E.C., Rhodes, J.S., Christian-Hinman, C.A., and Chung, H.J. (2021). Spontaneous seizure and memory loss in mice expressing an epileptic encephalopathy variant in the calmodulin-binding domain of Kv7.2. Proceedings of the National Academy of Sciences 118, e2021265118. 10.1073/pnas.2021265118.

40. Kirshner, Z.Z., and Gibbs, R.B. (2018). Use of the REVERT total protein stain as a loading control demonstrates significant benefits over the use of housekeeping proteins when analyzing brain homogenates by Western blot: An analysis of samples representing different gonadal hormone states. Molecular and Cellular Endocrinology 15:473:156–165. 10.1016/j.mce.2018.01.015.

41. Klinger, F., Gould, G., Boehm, S., and Shapiro, M.S. (2011). Distribution of M-channel subunits KCNQ2 and KCNQ3 in rat hippocampus. Neuroimage 58, 761–769. 10.1016/j.neuroimage.2011.07.003.

42. Kosaka, T. (1980). The axon initial segment as a synaptic site: ultrastructure and synaptology of the initial segment of the pyramidal cell in the rat hippocampus (CA3 region). J Neurocytol 9, 861–882.

43. Landrum, M.J., Lee, J.M., Benson, M., Brown, G.R., Chao, C., Chitipiralla, S., Gu, B., Hart, J., Hoffman, D., Jang, W., et al. (2018). ClinVar: improving access to variant interpretations and supporting evidence. Nucleic Acids Res 46, D1062–D1067. 10.1093/nar/gkx1153. Accessed Aug 14, 2023.

44. Lawrence, J.J., Saraga, F., Churchill, J.F., Statland, J.M., Travis, K.E., Skinner, F.K., and McBain, C.J. (2006). Somatodendritic Kv7/KCNQ/M channels control interspike interval in hippocampal interneurons. J Neurosci 26, 12325–12338.

45. Li, X., Zhang, Q., Guo, P., Fu, J., Mei, L., Lv, D., Wang, J., Lai, D., Ye, S., Yang, H., and Guo, J. (2021). Molecular basis for ligand activation of the human KCNQ2 channel. Cell Res 31, 52–61. 10.1038/s41422-020-00410-8.

46. Livak, K.J., and Schmittgen, T.D. (2001). Analysis of Relative Gene Expression Data Using Real-Time Quantitative PCR and the 2−ΔΔCT Method. Methods 25, 402–408. 10.1006/meth.2001.1262.

47. Long, S.B., Campbell, E.B., and Mackinnon, R. (2005). Voltage sensor of Kv1.2: structural basis of electromechanical coupling. Science 309, 903–908.

48. MacArthur, M.W., and Thornton, J.M. (1996). Deviations from planarity of the peptide bond in peptides and proteins. J Mol Biol 264, 1180–1195. 10.1006/jmbi.1996.0705.

49. MacKinnon, R., and Yellen, G. (1990). Mutations affecting TEA blockade and ion permeation in voltage-activated K^+^ channels. Science 250, 276–279.

50. Martinello, K., Huang, Z., Lujan, R., Tran, B., Watanabe, M., Cooper, E.C., Brown, D.A., and Shah, M.M. (2015). Cholinergic afferent stimulation induces axonal function plasticity in adult hippocampal granule cells. Neuron 85, 346–363. 10.1016/j.neuron.2014.12.030.

51. Martire, M., Castaldo, P., D’Amico, M., Preziosi, P., Annunziato, L., and Taglialatela, M. (2004). M channels containing KCNQ2 subunits modulate norepinephrine, aspartate, and GABA release from hippocampal nerve terminals. J Neurosci 24, 592–597.

52. McGuigan, F.E., and Ralston, S.H. (2002). Single nucleotide polymorphism detection: allelic discrimination using TaqMan. Psychiatric genetics 12, 133–136. 10.1097/00041444-200209000-00003.

53. Middleton, P.G., Mall, M.A., Drevinek, P., Lands, L.C., McKone, E.F., Polineni, D., Ramsey, B.W., Taylor-Cousar, J.L., Tullis, E., Vermeulen, F., et al. (2019). Elexacaftor-Tezacaftor-Ivacaftor for Cystic Fibrosis with a Single Phe508del Allele. N Engl J Med 381, 1809–1819. 10.1056/NEJMoa1908639.

54. Milh, M., Roubertoux, P., Biba, N., Chavany, J., Spiga Ghata, A., Fulachier, C., Collins, S.C., Wagner, C., Roux, J.C., Yalcin, B., et al. (2020). A knock-in mouse model for KCNQ2-related epileptic encephalopathy displays spontaneous generalized seizures and cognitive impairment. Epilepsia 61, 868–878. 10.1111/epi.16494.

55. Miller, C., Moczydlowski, E., Latorre, R., and Phillips, M. (1985). Charybdotoxin, a protein inhibitor of single Ca2+-activated K+ channels from mammalian skeletal muscle. Nature 313, 316–318. 10.1038/313316a0.

56. Millichap, J.J., and Cooper, E.C. (2012). KCNQ2 Potassium Channel Epileptic Encephalopathy Syndrome: Divorce of an Electro-Mechanical Couple? Epilepsy currents / American Epilepsy Society 12, 150–152. 10.5698/1535-7511-12.4.150.

57. Millichap, J.J., Park, K.L., Tsuchida, T., Ben-Zeev, B., Carmant, L., Flamini, R., Joshi, N., Levisohn, P.M., Marsh, E., Nangia, S., et al. (2016). KCNQ2 encephalopathy: Features, mutational hot spots, and ezogabine treatment of 11 patients. Neurol Genet 2, e96. 10.1212/NXG.0000000000000096.

58. Myers, C.T., Hollingsworth, G., Muir, A.M., Schneider, A.L., Thuesmunn, Z., Knupp, A., King, C., Lacroix, A., Mehaffey, M.G., Berkovic, S.F., et al. (2018). Parental Mosaicism in "De Novo" Epileptic Encephalopathies. N Engl J Med 378, 1646–1648. 10.1056/NEJMc1714579.

59. NCBI (2023). ClinVar. https://www.ncbi.nlm.nih.gov/clinvar/variation/2571144/?oq=SCV004011354.

60. Nissenkorn, A., Kornilov, P., Peretz, A., Blumkin, L., Heimer, G., Ben-Zeev, B., and Attali, B. (2021). Personalized treatment with retigabine for pharmacoresistant epilepsy arising from a pathogenic variant in the KCNQ2 selectivity filter. Epileptic Disord 23, 695–705. 10.1684/epd.2021.1315.

61. Numis, A.L., Angriman, M., Sullivan, J.E., Lewis, A.J., Striano, P., Nabbout, R., and Cilio, M.R. (2014). KCNQ2 encephalopathy: delineation of the electroclinical phenotype and treatment response. Neurology 82, 368–370. 10.1212/WNL.0000000000000060.

62. Olson, H.E., Kelly, M., LaCoursiere, C.M., Pinsky, R., Tambunan, D., Shain, C., Ramgopal, S., Takeoka, M., Libenson, M.H., Julich, K., et al. (2017). Genetics and genotype-phenotype correlations in early onset epileptic encephalopathy with burst suppression. Annals of neurology 81, 419–429. 10.1002/ana.24883.

63. Orhan, G., Bock, M., Schepers, D., Ilina, E.I., Reichel, S.N., Loffler, H., Jezutkovic, N., Weckhuysen, S., Mandelstam, S., Suls, A., et al. (2014). Dominant-negative Effects of KCNQ2 Mutations are Associated with Epileptic Encephalopathy. Annals of neurology. 10.1002/ana.24080.

64. Pan, Z., Kao, T., Horvath, Z., Lemos, J., Sul, J.Y., Cranstoun, S.D., Bennett, V., Scherer, S.S., and Cooper, E.C. (2006). A common ankyrin-G-based mechanism retains KCNQ and NaV channels at electrically active domains of the axon. J Neurosci 26, 2599–2613.

65. Pan, Z., Selyanko, A.A., Hadley, J.K., Brown, D.A., Dixon, J.E., and McKinnon, D. (2001). Alternative splicing of KCNQ2 potassium channel transcripts contributes to the functional diversity of M-currents. J Physiol 531, 347–358.

66. Pisano, T., Numis, A.L., Heavin, S.B., Weckhuysen, S., Angriman, M., Suls, A., Podesta, B., Thibert, R.L., Shapiro, K.A., Guerrini, R., et al. (2015). Early and effective treatment of KCNQ2 encephalopathy. Epilepsia 56, 685–691. 10.1111/epi.12984.

67. Reddi, R., Matulef, K., Riederer, E.A., Whorton, M.R., and Valiyaveetil, F.I. (2022). Structural basis for C-type inactivation in a *Shaker* family voltage-gated K^+^ channel. Sci Adv 8, eabm8804. 10.1126/sciadv.abm8804.

68. Robbins, J., Passmore, G.M., Abogadie, F.C., Reilly, J.M., and Brown, D.A. (2013). Effects of KCNQ2 Gene Truncation on M-Type Kv7 Potassium Currents. PLoS ONE 8, e71809. 10.1371/journal.pone.0071809.

69. Ronen, G.M., Rosales, T.O., Connolly, M., Anderson, V.E., and Leppert, M. (1993). Seizure characteristics in chromosome 20 benign familial neonatal convulsions. Neurology 43, 1355–1360.

70. Scheffer, I.E., Berkovic, S., Capovilla, G., Connolly, M.B., French, J., Guilhoto, L., Hirsch, E., Jain, S., Mathern, G.W., Moshe, S.L., et al. (2017). ILAE classification of the epilepsies: Position paper of the ILAE Commission for Classification and Terminology. Epilepsia 58, 512–521. 10.1111/epi.13709.

71. Schroeder, B.C., Kubisch, C., Stein, V., and Jentsch, T.J. (1998). Moderate loss of function of cyclic-AMP-modulated KCNQ2/KCNQ3 K channels causes epilepsy. Nature 396, 687–690.

72. Schwake, M., Athanasiadu, D., Beimgraben, C., Blanz, J., Beck, C., Jentsch, T.J., Saftig, P., and Friedrich, T. (2006). Structural determinants of M-type KCNQ (Kv7) K^+^ channel assembly. J Neurosci 26, 3757–3766.

73. Schwarz, J.R., Glassmeier, G., Cooper, E.C., Kao, T.C., Nodera, H., Tabuena, D., Kaji, R., and Bostock, H. (2006). KCNQ channels mediate IKs, a slow K+ current regulating excitability in the rat node of Ranvier. J Physiol 573, 17–34.

74. Shah, M.M., Migliore, M., Valencia, I., Cooper, E.C., and Brown, D.A. (2008). Functional significance of axonal Kv7 channels in hippocampal pyramidal neurons. Proc Natl Acad Sci U S A 105, 7869–7874.

75. Singh, N.A., Otto, J.F., Dahle, E.J., Pappas, C., Leslie, J.D., Vilaythong, A., Noebels, J.L., White, H.S., Wilcox, K.S., and Leppert, M.F. (2008). Mouse models of human KCNQ2 and KCNQ3 mutations for benign familial neonatal convulsions show seizures and neuronal plasticity without synaptic reorganization. J Physiol 586, 3405–3423. 10.1113/jphysiol.2008.154971.

76. Soh, H., Pant, R., LoTurco, J.J., and Tzingounis, A.V. (2014). Conditional deletions of epilepsy-associated KCNQ2 and KCNQ3 channels from cerebral cortex cause differential effects on neuronal excitability. J Neurosci 34, 5311–5321. 10.1523/JNEUROSCI.3919-13.2014.

77. Soh, H., Park, S., Ryan, K., Springer, K., Maheshwari, A., and Tzingounis, A.V. (2018). Deletion of KCNQ2/3 potassium channels from PV+ interneurons leads to homeostatic potentiation of excitatory transmission. Elife 7. 10.7554/eLife.38617.

78. Sun, J., and MacKinnon, R. (2020). Structural basis of human KCNQ1 modulation and gating. Cell 180, 340–347 e349. 10.1016/j.cell.2019.12.003.

79. Symonds, J.D., Zuberi, S.M., Stewart, K., McLellan, A., O’Regan, M., MacLeod, S., Jollands, A., Joss, S., Kirkpatrick, M., Brunklaus, A., et al. (2019). Incidence and phenotypes of childhood-onset genetic epilepsies: a prospective population-based national cohort. Brain 142, 2303–2318. 10.1093/brain/awz195.

80. Takacs, D.S. (2023). Infantile epileptic spasms syndrome: Management and prognosis. In UpToDate, T.W. Post, ed. (Wolters Kluwer), https://www.uptodate.com/contents/infantile-epileptic-spasms-syndrome-management-and-prognosis.

81. Tan, X.F., Bae, C., Stix, R., Fernandez-Marino, A.I., Huffer, K., Chang, T.H., Jiang, J., Faraldo-Gomez, J.D., and Swartz, K.J. (2022). Structure of the *Shaker* Kv channel and mechanism of slow C-type inactivation. Sci Adv 8, eabm7814. 10.1126/sciadv.abm7814.

82. Tatulian, L., Delmas, P., Abogadie, F.C., and Brown, D.A. (2001). Activation of expressed KCNQ potassium currents and native neuronal M-type potassium currents by the anti-convulsant drug retigabine. J Neurosci 21, 5535–5545. 21/15/5535 [pii].

83. Tran, B., Ji, Z.G., Xu, M., Tsuchida, T.N., and Cooper, E.C. (2020). Two KCNQ2 Encephalopathy Variants in the Calmodulin-Binding Helix A Exhibit Dominant-Negative Effects and Altered PIP2 Interaction. Front Physiol 11, 1144. 10.3389/fphys.2020.571813.

84. Traynelis, J., Silk, M., Wang, Q., Berkovic, S.F., Liu, L., Ascher, D.B., Balding, D.J., and Petrovski, S. (2017). Optimizing genomic medicine in epilepsy through a gene-customized approach to missense variant interpretation. Genome Res 27, 1715–1729. 10.1101/gr.226589.117.

85. Truty, R., Patil, N., Sankar, R., Sullivan, J., Millichap, J., Carvill, G., Entezam, A., Esplin, E.D., Fuller, A., Hogue, M., et al. (2019). Possible precision medicine implications from genetic testing using combined detection of sequence and intragenic copy number variants in a large cohort with childhood epilepsy. Epilepsia Open 4, 397–408. 10.1002/epi4.12348.

86. Vanoye, C.G., Desai, R.R., Ji, Z., Adusumilli, S., Jairam, N., Ghabra, N., Joshi, N., Fitch, E., Helbig, K.L., McKnight, D., et al. (2022). High-throughput evaluation of epilepsy-associated KCNQ2 variants reveals functional and pharmacological heterogeneity. JCI Insight 7. 10.1172/jci.insight.156314.

87. Varghese, N., Moscoso, B., Chavez, A., Springer, K., Ortiz, E., Soh, H., Santaniello, S., Maheshwari, A., and Tzingounis, A.V. (2023). KCNQ2/3 Gain-of-Function Variants and Cell Excitability: Differential Effects in CA1 versus L2/3 Pyramidal Neurons. J Neurosci 43, 6479–6494. 10.1523/JNEUROSCI.0980-23.2023.

88. Watanabe, H., Nagata, E., Kosakai, A., Nakamura, M., Yokoyama, M., Tanaka, K., and Sasai, H. (2000). Disruption of the epilepsy KCNQ2 gene results in neural hyperexcitability. J Neurochem 75, 28–33.

89. Wechkuysen, S., and George, A.L., Jr., eds. (2022). KCNQ2-and KCNQ3-associated Epilepsy. Cambridge Elements series on Genetics in Epilepsy. Cambridge University Press. DOI: 10.1017/9781009278270

90. Weckhuysen, S., Ivanovic, V., Hendrickx, R., Van Coster, R., Hjalgrim, H., Moller, R.S., Gronborg, S., Schoonjans, A.S., Ceulemans, B., Heavin, S.B., et al. (2013). Extending the KCNQ2 encephalopathy spectrum: clinical and neuroimaging findings in 17 patients. Neurology 81, 1697–1703. 10.1212/01.wnl.0000435296.72400.a1.

91. Weckhuysen, S., Mandelstam, S., Suls, A., Audenaert, D., Deconinck, T., Claes, L.R., Deprez, L., Smets, K., Hristova, D., Yordanova, I., et al. (2012). KCNQ2 encephalopathy: emerging phenotype of a neonatal epileptic encephalopathy. Annals of neurology 71, 15–25. 10.1002/ana.22644.

92. Wu, Y., Yan, Y., Yang, Y., Bian, S., Rivetta, A., Allen, K., and Sigworth, F.J. (2023). Cryo-EM structures of Kv1.2 potassium channels, conducting and non-conducting. bioRxiv. 10.1101/2023.06.02.543446.

93. Yang, Y., Beyer, B.J., Otto, J.F., O’Brien, T.P., Letts, V.A., White, H.S., and Frankel, W.N. (2003). Spontaneous deletion of epilepsy gene orthologs in a mutant mouse with a low electroconvulsive threshold. Hum Mol Genet 12, 975–984.

94. Ye, W., Zhao, H., Dai, Y., Wang, Y., Lo, Y.H., Jan, L.Y., and Lee, C.H. (2022). Activation and closed-state inactivation mechanisms of the human voltage-gated K(V)4 channel complexes. Mol Cell 82, 2427–2442.e2424.

95. Zaydman, M.A., Silva, J.R., Delaloye, K., Li, Y., Liang, H., Larsson, H.P., Shi, J., and Cui, J. (2013). Kv7.1 ion channels require a lipid to couple voltage sensing to pore opening. Proc Natl Acad Sci U S A 110, 13180–13185. 10.1073/pnas.1305167110.

96. Zetoune, A.B., Fontaniere, S., Magnin, D., Anczukow, O., Buisson, M., Zhang, C.X., and Mazoyer, S. (2008). Comparison of nonsense-mediated mRNA decay efficiency in various murine tissues. BMC Genet 9, 83. 10.1186/1471-2156-9-83.

97. Zhang, J., Kim, E.C., Chen, C., Procko, E., Pant, S., Lam, K., Patel, J., Choi, R., Hong, M., Joshi, D., et al. (2020). Identifying mutation hotspots reveals pathogenetic mechanisms of KCNQ2 epileptic encephalopathy. Sci Rep 10, 4756. 10.1038/s41598-020-61697-6.

98. Zhao, Y., Chen, Z., Cao, Z., Li, W., and Wu, Y. (2019). Diverse Structural Features of Potassium Channels Characterized by Scorpion Toxins as Molecular Probes. Molecules 24. 10.3390/molecules24112045.

99. Zheng, Y., Liu, H., Chen, Y., Dong, S., Wang, F., Wang, S., Li, G.L., Shu, Y., and Xu, F. (2022). Structural insights into the lipid and ligand regulation of a human neuronal KCNQ channel. Neuron 110, 237–247 e234. 10.1016/j.neuron.2021.10.029.

